# Improving orthologous signal and model fit in datasets addressing the root of the animal phylogeny

**DOI:** 10.1101/2022.11.21.517274

**Authors:** Charley GP McCarthy, Peter O Mulhair, Karen Siu-Ting, Christopher J Creevey, Mary J O’Connell

## Abstract

There is conflicting evidence as to whether Porifera (sponges) or Ctenophora (comb jellies) comprise the root of the animal phylogeny. Support for either a Porifera-sister or Ctenophore-sister tree has been extensively examined in the context of model selection, taxon sampling and outgroup selection. The influence of dataset construction is comparatively understudied. We re-examine five animal phylogeny datasets that have supported either root hypothesis using an approach designed to enrich orthologous signal in phylogenomic datasets. We find that many component orthogroups in animal datasets fail to recover major animal lineages as monophyletic with the exception of Ctenophora, regardless of the supported root. Enriching these datasets to retain orthogroups recovering ≥3 major lineages reduces dataset size by up to 50% while retaining underlying phylogenetic information and taxon sampling. Site-heterogeneous phylogenomic analysis of these enriched datasets recovers both Porifera-sister and Ctenophora-sister positions, even with additional constraints on outgroup sampling. Two datasets which previously supported Ctenophora-sister support Porifera-sister upon enrichment. All enriched datasets display improved model fitness under posterior predictive analysis. While not conclusively rooting animals at either Porifera or Ctenophora, our results indicate that dataset size and construction as well as model fit influence animal root inference.

## Introduction

Animals comprise five major phyla: Bilateria, Cnidaria, Placozoa, Ctenophora and Porifera (Halanych, 2004; King and Rokas, 2017). There remain a number of conflicting hypotheses regarding the evolutionary history of the animals. Primary among these is the placement of either Porifera (sponges) or Ctenophora (comb jellies) as the sister to all other animals (**Fig. 1**) (King and Rokas, 2017). Porifera possess rudimentary epithelia and neural-like signalling while lacking muscle cells or a complete digestive system (King and Rokas, 2017; Nielsen, 2019). Ctenophora exhibit more complex morphology, e.g. a nervous system and true tissues, but possess different epithelial organization from other animals (King and Rokas, 2017; Belahbib et al., 2018; Nielsen, 2019). Although both phyla lack more-derived innovations found in Cnidaria and Bilateria, some sponges possess microRNA-like machinery and ctenophore genomes may retain phylum-specific copies of genes (Wheeler et al., 2009; Pastrana et al., 2019; Pett et al., 2019). The “Porifera-sister” hypothesis, which places sponges as sister to all other animals, has been the traditionally-accepted view of animal evolution based upon morphological and molecular observations (Halanych, 2004; Pisani et al., 2015; King and Rokas, 2017; Fernández et al., 2019; Nielsen, 2019) (**Fig. 1**). Since the advent of large-scale phylogenomics, several studies have supported an alternative “Ctenophora-sister” hypothesis where ctenophores are placed sister to all other animals (Dunn et al., 2008; Philippe et al., 2009; Ryan et al., 2013; Moroz et al., 2014; Chang et al., 2015; King and Rokas, 2017) (**Fig. 1**).

**Figure 1.**
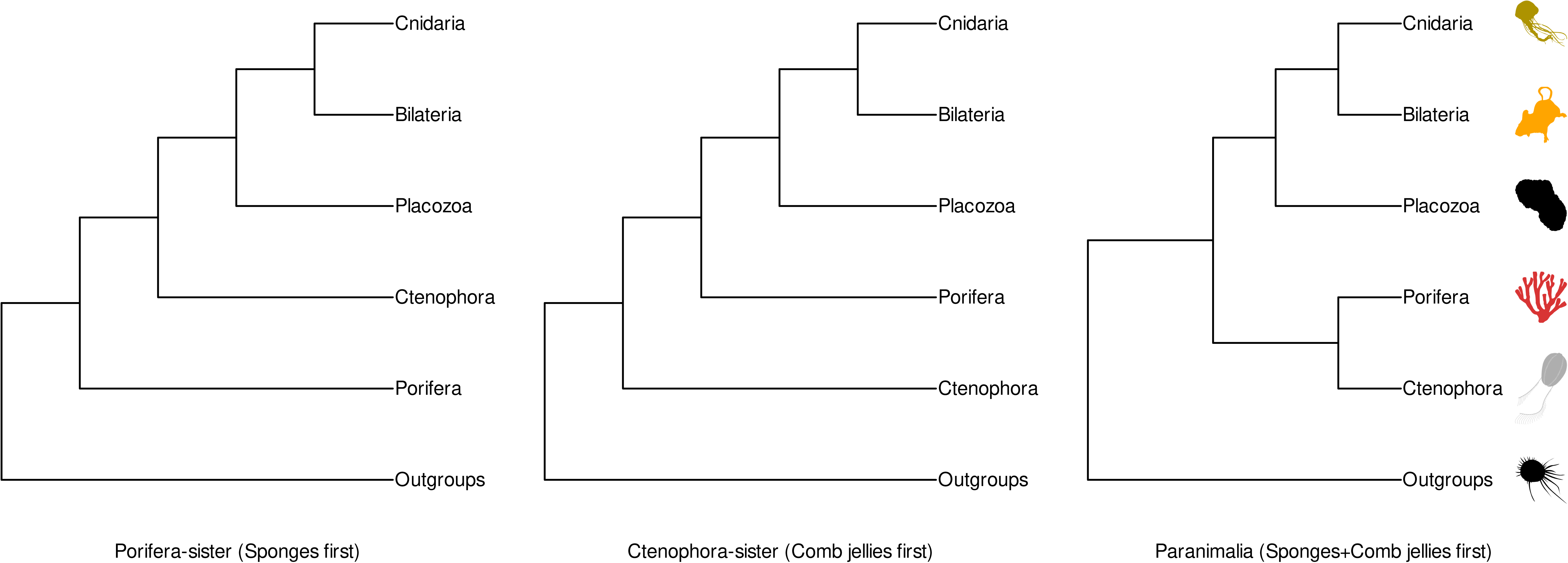
Overview of alternative hypotheses for the root of animals. Topologies shown describe the two major hypotheses for animal rooting (Porifera-sister and Ctenophore-sister), alongside an alternate hypothesis consisting of a monophyletic Porifera+Ctenophora-sister root (“Paranimalia”). Topologies for Porifera- and Ctenophora-sister roots as described in King and Rokas (2017). Nomenclature and topology for Paranimalia root as described in Francis and Canfield (2020). All silhouettes obtained from Phylopic (http://phylopic.org). Placozoa silhouette (*Trichoplax adhaerens*) by Oliver Voigt under a Creative Commons licence <u>CC BY-SA 3.0</u>; Bilateria silhouette (*Mus musculus*) by David Liao under a Creative Commons licence <u>CC BY-SA 3.0</u> and Porifera silhouette (*Siphonochalina siphonella*) by Mali’o Kodis, photograph by Derek Keats (http://www.flickr.com/photos/dkeats/) under a Creative Commons licence <u>CC BY 3.0</u>. Filozoa outgroup (*Capsaspora owczarzaki*), Cnidaria silhouette (Medusazoa sp.), and Ctenophora silhouette (*Hormiphora californensis*) under public domain.

Recent studies arguing in favour of either hypothesis have focused on the use of models which assume homogeneous or hetereogenous site evolution processes (Francis and Canfield, 2020). “Site-homogeneous” models assume similar amino acid equilibrium frequencies amongst partitions of a data matrix (Moran et al., 2015; Li et al., 2021). “Site-heterogeneous” models allow for heterogeneity in amino acid equilibrium frequencies across sites (Moran et al., 2015; Li et al., 2020). Both approaches have advantages and disadvantages. Site-homogeneous partitioned approaches can be readily implemented in maximum-likelihood frameworks, but may be vulnerable to long-branch attraction (LBA) artefacts (Li et al., 2021; Redmond and McLysaght, 2021). Site-heterogeneous approaches better reflect actual sequence evolution, but current implementations in Bayesian frameworks are computationally expensive. When applied in animal phylogenomics site-homogeneous approaches invariably recover a Ctenophora-sister tree (Borowiec et al., 2015; Chang et al., 2015; Pisani et al., 2015; Whelan et al., 2015, 2017; Simion et al., 2017; Laumer et al., 2018, 2019), whereas site-heterogeneous approaches have shown support for either Ctenophora-sister (Borowiec et al., 2015; Chang et al., 2015; Whelan et al., 2015, 2017; Laumer et al., 2018, 2019; Li et al., 2021) or Porifera-sister (Pisani et al., 2015; Feuda et al., 2017; Simion et al., 2017; Kapli and Telford, 2020; Redmond and McLysaght, 2021). It has been suggested that site-heterogeneous support for Porifera-sister may be dependent on data recoding strategies, selection of outgroup taxa, or the number of frequency classes afforded by site-heterogenous models (Feuda et al., 2017; Whelan and Halanych, 2017; Kapli and Telford, 2020; Li et al., 2021). Alternative approaches based on gene content have supported Porifera-sister (Pett et al., 2019).

The influence of individual gene families (or orthogroups) within animal datasets on root inference is comparatively underexplored. Gene loss, gene duplication, and genome duplication events have occurred extensively across animals (Dehal and Boore, 2005; Meyer and Van de Peer, 2005; Fernández and Gabaldón, 2020; Guijarro-Clarke et al., 2020). Both sponges and ctenophores have undergone gene loss after their divergence from other animals (Belahbib et al., 2018; Pett et al., 2019), and Ctenophora are thought to possess faster-evolving genomes than other animals (Pick et al., 2010; Philippe et al., 2011; Feuda et al., 2014). These factors can complicate ortholog detection within the animals and may contribute to conflicting placements of Porifera and Ctenophora (Glover et al., 2019; Deutekom et al., 2020; Fernández and Gabaldón, 2020). Some studies have examined the effect of different classes of sequences on animal root inference (Philippe et al., 2011; Nosenko et al., 2013; Whelan et al., 2015). Most animal phylogeny datasets are constructed from combinations of genomic and transcriptomic data using automated/semi-automated ortholog detection pipelines, and include rigorous protocols to remove orthogroups which may contain LBA or compositional heterogeneity (CH) artefacts (Whelan et al., 2015; Simion et al., 2017; Whelan et al., 2017; Laumer et al., 2018, 2019). There is evidence that component orthogroups in these datasets may favour Ctenophora-sister (Shen et al., 2017), but a recent study found that the removal of a small fraction of sites from one such dataset recovered a monophyletic Ctenophora+Porifera root (“Paranimalia”) under multiple models of evolution (**Fig. 1**) (Francis and Canfield, 2020).

Ortholog misidentification has been shown to have a substantial impact on final species tree inference within the animals (Brown and Thomson, 2017; Siu-Ting et al., 2019; Natsidis et al., 2021). Assessment of internal congruence within these datasets (i.e. the ability of individual orthogroups to recover internal relationships in a tree) has only been applied in limited instances (Simion et al., 2017). One approach to reducing artefacts arising from ortholog detection is to assess the ability of orthologs to recover uncontroversial relationships within a species tree. Such assessment has previously been applied in similar conflicts within Lissamphibia lineages prone to hidden paralogy, where paralogous genes are misidentified as orthologous (Kuraku 2010; Siu-Ting et al. 2019). For datasets constructed to answer questions of placement of deeper nodes in a tree (e.g. Porifera-sister vs. Ctenophora-sister), it may be prudent to ask whether these datasets can recover at least a proportion of these nodes themselves (Doolittle and Brown, 1994; Smith and Hahn, 2021; Hime et al., 2021).

We have examined the effects of dataset incongruence on conflicting animal root hypotheses, using an approach to enrich orthologous signal as implemented in the software clan_check (Siu-Ting et al, 2019). We re-examined five previously-published animal datasets which have variously supported either a Porifera-sister or Ctenophora-sister root (**Table 1**) (Chang et al., 2015; Whelan et al., 2015; Simion et al., 2017; Whelan et al., 2017). We find that for all but one of these datasets, component orthogroups overwhelmingly recover Ctenophora as a monophyletic clan when ≥2 ctenophore taxa are present. By contrast, <10% of orthogroups across all datasets recover a monophyletic Porifera and <50% recover monophyletic Cnidaria or Bilateria. Filtering these datasets to retain orthogroups which recover ≥3 clans reduces dataset size by up to 50% while retaining taxon sampling and underlying phylogenetic information. We then reconstructed an animal phylogeny for each dataset using the site-heterogeneous CAT-GTR+G4 model as implemented in PhyloBayes-MPI. We find that filtered datasets exhibit improved model fit under posterior predictive analysis relative to original datasets, while maintaining sufficient amino acid substitution and transition rate information. As for the animal root, we recover both Porifera-sister and Ctenophora-sister trees across the five datasets examined and across additional analyses restricting outgroup sampling. Notably, two datasets previously demonstrated to support Ctenophora-sister instead support Porifera-sister after such reanalysis. This study does not conclusively resolve the animal root. However, our findings illustrate the importance of assessing ortholog suitability and dataset congruence in animal phylogeny and may indicate that smaller enriched datasets contain sufficient information to reconstruct complex evolutionary histories while facilitating better model fit than larger datasets.

**Table 1.** Animal phylogeny datasets used. . Table includes number of taxa and orthogroups sampled per dataset, the number of sites across the entire dataset and information as to the phylogenetic approaches and rooting for each dataset in each original study (note: other datasets in Whelan et al. (2015) were analyzed under CAT-GTR+G4). The datasets chosen column includes our nomenclature for each dataset and, for studies with multiple datasets, which dataset was selected. OG: orthogroups, ML: maximum-likelihood, CAT: CAT mixture model, GTR: generalised time reversible model, G4: discrete gamma distribution with 4 categories.

| Study | Dataset chosen | # Taxa | # OGs | # Sites | Rooting in original study | Model(s) used in original study | Data availability |
| --- | --- | --- | --- | --- | --- | --- | --- |
| Chang et al. (2015) | Chang2015 | 77 | 200 | 51,940 | Ctenophora-sister | ML partitioning CAT | https://treebase.org/treebase-web/search/study/summary.html?id=17743 |
| Whelan et al. (2015) | Whelan2015_D10 ("Dataset 10") | 70 | 210 | 59,733 | Ctenophora-sister | ML partitioning CAT-GTR+G4 (other datasets) | http://dx.doi.org/10.6084/m9.figshare.1334306.v3 |
| Whelan et al. (2015) | Whelan2015_D20 ("Dataset 20") | 70 | 178 | 47,632 | Ctenophora-sister | ML partitioning CAT-GTR+G4 (other datasets) | http://dx.doi.org/10.6084/m9.figshare.1334306.v3 |
| Simion et al. (2017) | Simion2017 | 97 | 1,719 | 401,632 | Porifera-sister | ML partitioning Jack-knifed CAT+G4 | https://github.com/psimion/SuppData_Metazoa_2017 |
| Whelan et al. (2017) | Whelan2017_MCRS ("Metazoa_Choano_RCFV_strict") | 76 | 127 | 49,388 | Ctenophora-sister | ML partitioning CAT-GTR+G4 | http://dx.doi.org/10.6084/m9.figshare.4484138.v1 |

## Results

### Lack of orthologous signal at major nodes in animal datasets

The five animal phylogeny datasets analyzed in this study were chosen to reflect competing animal root hypotheses, and differences in taxon sampling and dataset construction approaches (**Tables 1 and S1**) (Chang et al., 2015; Whelan et al., 2015; Simion et al., 2017; Whelan et al., 2017; Siu-Ting et al., 2019). The latter can also be observed by the relative lack of overlap in human gene content across all five datasets (Francis and Canfield 2020) (**Fig. S1**). We filtered these datasets to enrich for orthologous signal at major animal nodes using an approach implemented in the software clan_check (Siu-Ting et al., 2019). Clan_check takes user-defined sets of taxa and tests gene trees on their ability to recover these sets as clans *sensu* Wilkinson et al. (2007), provided that ≥2 taxa from a set are present in a tree (Wilkinson et al., 2007; Siu-Ting et al., 2019) (**Fig. 2a**). This approach has previously been demonstrated to improve resolution of problematic nodes within the Lissamphibia (Siu-Ting et al., 2019). We defined six groups to test as clans: the five major animal phyla and an additional clan of all outgroup taxa for each dataset. As Placozoa was solely represented by *Trichoplax adhaerens* in each dataset, in practice we assessed orthologous signal of five clans in our study. For the Chang et al. (2015) and Simion et al. (2017) studies, the original datasets were obtained for analysis and are henceforth referred to as “Chang2015” and “Simion2017” (**Table 1**). For the two Whelan et al. (2015, 2017) studies, the datasets chosen were those presented in the main text of each paper: Dataset 10 for Whelan et al. (2015) and Metazoa_Choano_RCFV_strict for Whelan et al. (2017) (**Supplementary Information**). Additionally, Dataset 20 from Whelan et al. (2015) was chosen as it has been used in other animal phylogenomics studies (Feuda et al. 2017, Li et al. 2020). These datasets are henceforth referred to as “Whelan2015_D10”, “Whelan2015_D20” and “Whelan2017_MCRS” (**Table 1**).

**Figure 2.**
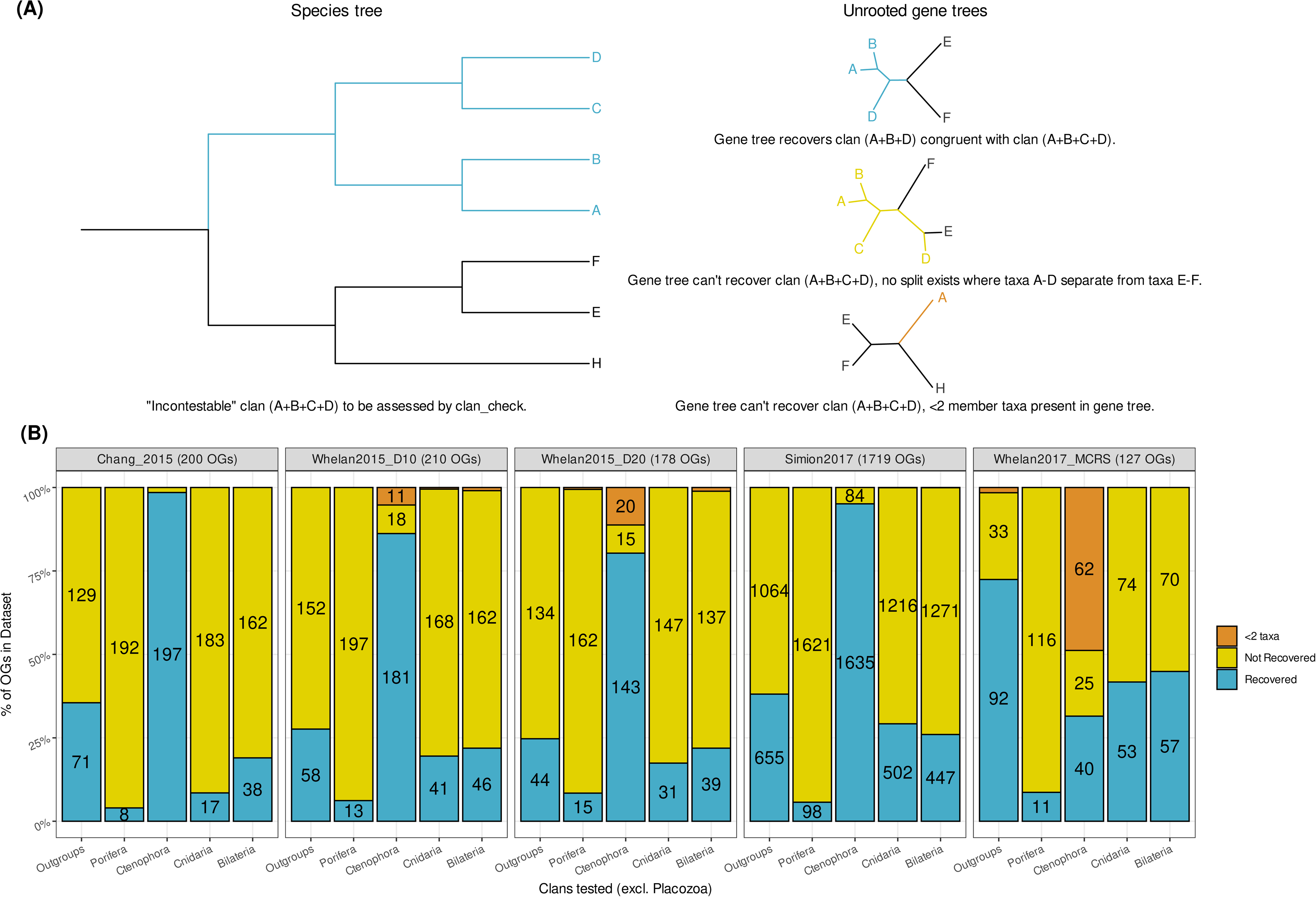
**(a) Simplified illustration of clan_check approach to assessing congruence**. In unrooted trees a “clan”, *sensu* Wilkinson et al. (2007), is analogous to monophyletic groups or clades in rooted trees. Clan_check assesses whether user-defined “incontestable” clans are violated in a set of unrooted gene trees. These “incontestable” clans can be thought of as phylogenetic relationships that should be recapitulated by gene trees to the exclusion of all other taxa if ≥2 taxa representing that relationship are present. In our study, we refer to clans which are not violated as “recovered” and clans which are violated as “not recovered”. In the example given, the relationship A+B+C+D (blue) is considered incontestable within a rooted species tree (left) and is assessed across a group of unrooted gene trees (right). In the first gene tree (top right), A+B+C+D is recovered as a clan because A+B+D (blue) group together to the exclusion of E and F. In the second gene tree (middle right), A+B+C+D is not recovered as a clan as no bipartition exists in which those taxa group to the exclusion of E and F (yellow). In the third gene tree (bottom right), A+B+C+D cannot be recovered as a clan as only taxon A is present in the gene tree (orange). **(b) Clan_check results for 5 animal phylogeny datasets**. Stacked bar plot representing proportions of orthogroups per dataset which can recover/cannot recover/do not contain enough taxa (2 or more) to recover a given clan from five animal clans defined in this study excluding the singleton clan Placozoa. Numbers of orthogroups in each of the three categories per clan given inside bars, values less than 8 not shown. OG: orthogroups.

Clan_check analysis indicates a lack of orthologous signal for deeper relationships within animal phylogeny datasets (**Fig. 2b**). In four of the five datasets analyzed, 86-98% of component orthogroups recover Ctenophora as a clan. This contrasts with recovery of the remaining three animal clans; <10% of orthogroups can recover Porifera, ∼9-30% can recover Cnidaria and ∼19-26% can recover Bilateria. While outgroup composition varies across these four datasets, ∼26-38% of orthogroups recover an outgroup clan separate from the remaining animal taxa. One exception to this trend is the lower proportion of Ctenophora recovery observed in the Whelan2017_MCRS dataset, where 48% of orthogroups have <2 ctenophores present and 35% lack any ctenophores whatsoever. There is also greater recovery of the outgroup clan across Whelan2017_MCRS (∼73%) compared to the other datasets, a reflection of Choanoflagellate-only outgroup sampling in Whelan2017_MCRS. **Table 2** indicates the number of clans (excluding Placozoa) recovered by each dataset; nearly three quarters of orthogroups across all five datasets cannot recover >2 defined clans.

**Table 2.** Clan recovery across five animal datasets using clan_check. Table lists the number of orthogroups in each dataset, and then how many orthogroups within each dataset can recover *n* number of the defined clans given in Figure 2. The numbers of orthogroups which recover ≥3 clans is emphasised in-table for *n* = 3-5, and the total number of orthogroups which passed this filter and were retained in our filtered datasets is given on the right. OG: orthogroups.

| Dataset | OGs in original dataset | # of OGs which recovered $n$ number of clans (excluding Placozoa) | | | | | | OGs retained after clan_check filtering |
| --- | --- | --- | --- | --- | --- | --- | --- | --- |
| | | $n = 5$ | 4 | 3 | 2 | 1 | 0 | |
| Chang2015 | 200 | 1 | 8 | 25 | 56 | 107 | 3 | 34 |
| Whelan2015_D10 | 210 | 1 | 15 | 24 | 51 | 100 | 19 | 40 |
| Whelan2015_D20 | 178 | 2 | 8 | 19 | 45 | 83 | 21 | 29 |
| Simion2017 | 1719 | 25 | 125 | 307 | 583 | 625 | 54 | 457 |
| Whelan2017MCRS | 127 | 3 | 19 | 20 | 33 | 36 | 16 | 42 |

### The effect of clan_check filtering on animal phylogeny datasets

We retained orthogroups that could recover ≥2 clans defined in our clan_check analysis for downstream analysis (**Table 2**). Applying this filter reflects a reasonable assumption that orthogroups of sufficient sampling and quality should be capable of recapitulating most major internal relationships within a species tree. This filter retained between 17-33% of orthogroups from the original datasets and 25-52% of the original dataset size (**Tables 1-3**), while retaining taxon sampling distribution across each major clan (**Fig. S2**). The filtered datasets also display similar or reduced levels of data heterogeneity relative to the original datasets (**Fig. S3, Table S2**) and similar distributions of gene ontology categories (**Table S3**), with some exceptions that can be attributed to dataset construction approaches.

**Table 3.** PhyloBayes-MPI CAT-GTR reconstructions of five filtered animal datasets. Table includes number of taxa and orthogroups sampled per filtered dataset, the number of sites across each dataset, details on the convergence/divergence of each set of runs in tree and component space, and supported root hypothesis. OG: orthogroups, diff: discrepancy, maxdiff: maximum discrepancy observed across all bipartitions, meandiff: mean discrepancy observed across all bipartitions, rel_diff: relative discrepancy between chain components.

| Dataset<br>(Iterations per chain) | #<br>Taxa | # OGs | # Sites | Tree convergence | Conflicting<br>bipartitions<br>(diff > 0) | Divergent<br>components<br>(rel_diff < 0.3) | Root hypothesis |
| --- | --- | --- | --- | --- | --- | --- | --- |
| Chang2015_filtered<br>(12,768 / 14,720) | 77 | 34 | 15,983 | Maxdiff: 0.12<br>Meandiff: 0.00144368 | 10 | Tree length<br>(0.447) | Ctenophora-sister |
| Whelan2015_D10_filtered<br>(18,587 / 19,295) | 70 | 40 | 15,125 | Maxdiff: 0.234<br>Meandiff: 0.00375807 | 19 | None | Ctenophora-sister |
| Whelan2015_D20_filtered<br>(12,258 / 12,418) | 70 | 29 | 10,0016 | Maxdiff: 0.204<br>Meandiff: 0.00281874 | 12 | None | Porifera-sister |
| Simion2017_filtered<br>(8,776 / 9,471) | 97 | 457 | 152,324 | Maxdiff: 1<br>Meandiff: 0.0104712 | 4 | Log-likelihood<br>(0.85), mean<br>site entropy<br>(0.73), alpha<br>(0.76) | Porifera-sister |
| Whelan2017_MCRS_filtered<br>(15,714 / 12,226) | 76 | 42 | 22,820 | Maxdiff: 0.038<br>Meandiff: 0.000906953 | 33 | None | Porifera-sister |

We assessed the impact of clan_check filtering on dataset construction using PhyKIT, which performs several sequence- and tree-based evaluations of phylogenetic information and potential biases (Steenwyk et al., 2021). For each dataset, we evaluated phylogenetic information across orthogroups which passed or failed our clan_check filter using seven criteria following Steenwyk et al. (2021) (**Fig. 3**). Orthogroups which passed our clan_check filtering exhibit significantly higher alignment lengths and numbers of parsimony informative and variable sites (Wilcoxson test, p ≤ 0.05) - all measures associated with higher-quality phylogenetic information (Shen et al., 2016). Mean bipartition support, or internal branch support, is also significantly improved in orthogroups that passed our clan_check filter (**Fig. 3**). Other metrics, such as long branch score or treeness divided by relative compositional variability, show no significant difference before or after clan_check filtering in most datasets - indicating that there is still heterogeneity present in these datasets (**Fig. 3**).

**Figure 3.**
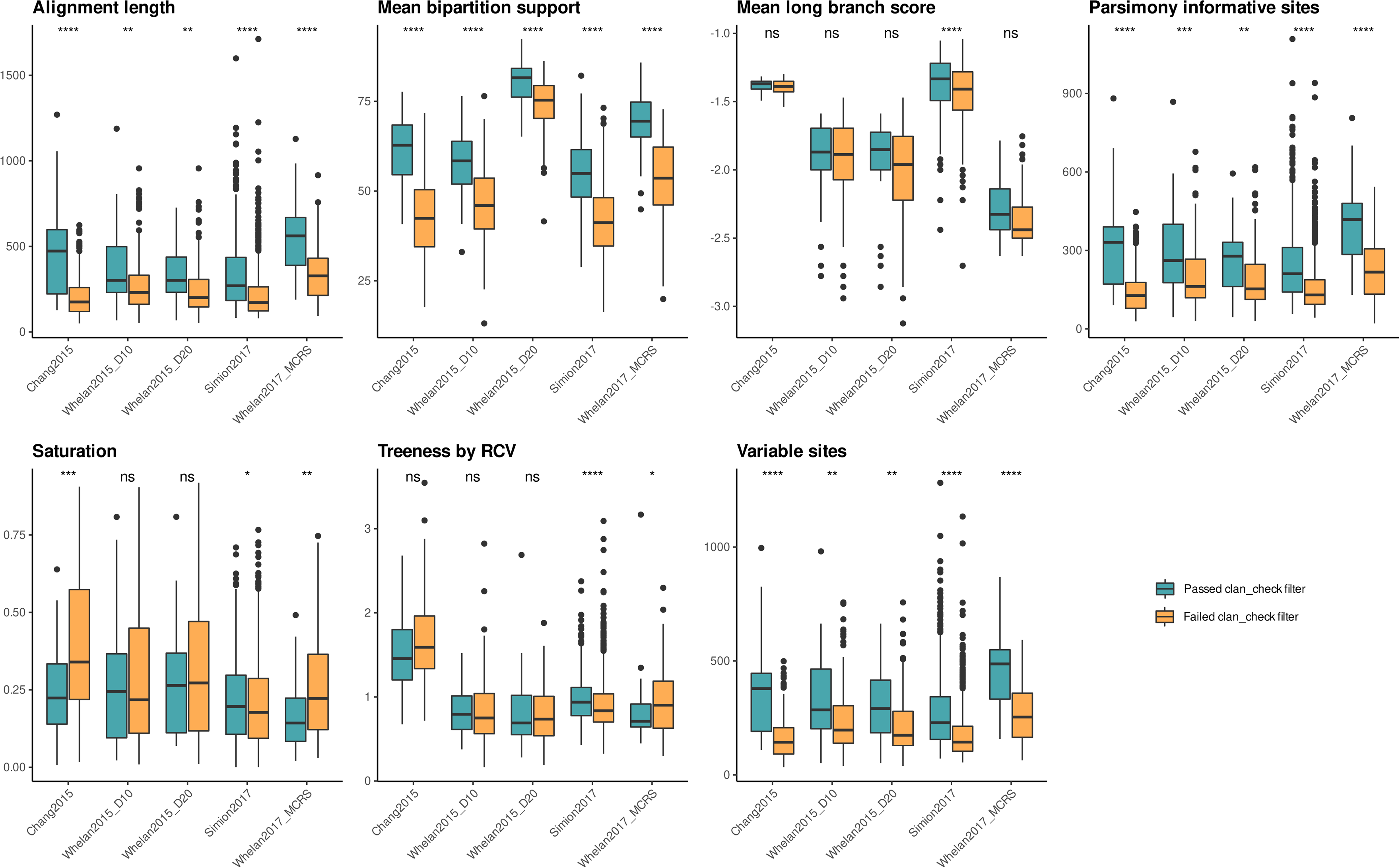
Comparison of seven phylogenetic information criteria in orthogroups passing or failing clan_check filter. Sequence- and tree-based phylogenetic information was assessed using PhyKIT (Steenwyk et al. (2021)). Alignment length: longer alignments are associated with stronger phylogenetic information; Mean bipartition support: higher values imply greater certainty in tree topology; Mean long branch score: lower values indicate less long branch attraction artefacts; Parsimony informative sites: higher number of sites associated with improved phylogenetic information; Saturation: higher scores indicate less sequence saturation; Treeness by Relative Compositional Variability (RCV): higher values indicate less susceptibility to composition bias; Variable sites: higher number of sites associated with stronger phylogenetic information content. Box plot representing distribution of values for each criterion in original and filtered dataset. Wilcoxson test results: **** = p ≤ 0.0001, *** = p ≤ 0.001, ** = p ≤ 0.01, * = p ≤ 0.05, ns = not significant.

We also examined the effect of our clan_check filtering on the overall gene- and site concordance factors in each dataset, using methods implemented in IQTREE (Nguyen et al. 2015; Minh et al. 2020). For a given branch in a species tree, these factors represent the percentage of gene trees or alignment sites in a dataset containing or supporting that branch for all gene trees or sites “decisive” for that branch – i.e. those capable of accepting or rejecting that branch (Minh et al. 2020). Gene concordance factors (gCFs) in some respects recapitulate the results of our clan_check analysis. For example, in the Chang2015 dataset we observe 8 orthogroups recovering a Porifera clan out of 200 orthogroups containing ≥2 sponges (**Fig. 2b**) – the branch representing the ancestral Porifera node is given a gCF of 4 (8 / 200) in our concordance analysis of the same dataset (**Fig. S4**). Comparing the original and filtered datasets, we observe an increase in gCFs across most branches in the filtered datasets – as would be expected given the reduction in the number of orthogroups in the latter (**Figs. S4-S8**). Site concordance factor (sCF) analysis indicates more incremental change between original and filtered datasets, which reflects that our approach filters on the gene-level as opposed to site-level (**Figs. S4-S8**). The sCFs of many deep branches in both original and filtered datasets are below 50%, suggesting that there is substantial conflicting signal for deeper animal relationships present in these datasets at the site-level. Notably, the sCF of the Ctenophora branch is consistently higher (∼70-75%) than other major animal nodes across all original and filtered datasets (**Fig. S4-S8**).

### Improved model fit observed in datasets filtered using clan_check

Phylogenomic reconstruction of each enriched animal dataset was performed using PhyloBayes-MPI under a CAT-GTR+G4 model, running two independent chains for at least 10,000 iterations or in the case of Simion2017_filtered, at least 7,500 iterations (**Table 3**) (Lartillot and Philippe 2004; Lartillot et al. 2013). Visual and quantitative assessment of chain convergence of each run indicated that all parameters had acceptably converged, with exceptions in Chang2015_filtered and Simion2017_filtered (**Table 3**, **Figs. S9-13**, **Supplementary Information**). We assessed model fitness for each filtered dataset using five posterior predictive analysis (PPA) statistics implemented in PhyloBayes-MPI (Lartillot et al., 2013). Each statistic tests a model’s ability to accommodate site- or branch-heterogeneity by comparing observed values from phylogenomic datasets with data simulated under the model. This comparison is computed as a |Z|-score, with the null hypothesis that the model adequately estimates these preferences. |Z| < 2 indicates adequate model fitness, whereas |Z| > 5 indicates strong rejection of the null hypothesis (Feuda et al., 2017; Lartillot, 2020). PPAs for each filtered dataset were compared with PPAs for the original dataset, which were obtained from Feuda et al. (2017) for Chang2015 and Whelan2015_D20 or generated from additional PhyloBayes-MPI runs for all other datasets.

In line with reductions in dataset size and enrichment of phylogenetic information, PPAs for each filtered dataset indicate improved fit to CAT-GTR+G4 over the original datasets (**Fig. 5a**). Datasets constructed with strict paralogy and/or heterogeneity filtering criteria, particularly Whelan2015_D20 and Whelan2017_MCRS, show smaller improvements relative to other datasets (**Fig. 4a**). Of note is the substantial improvement in scores for PPA-MAX, which tests for adequate modelling of maximal compositional heterogeneity, observed for Chang2015_filtered and Whelan2017_MCRS_filtered (|Z| < 2). This may reflect the reduction of compositional outliers inferred from comparing RCFV values in original and filtered datasets (**Fig. S3**). Overall, CAT-GTR+G4 fails to adequately model most compositional heterogeneity in filtered animal datasets despite the improvements in PPA results (**Fig. 4a, Fig. S26**). This is in line with observations by Feuda et al. (2017), although the same authors demonstrated CAT-GTR+G4 displays substantially better model fitness than site-homogeneous approaches (Feuda et al. 2017).

**Figure 4.**
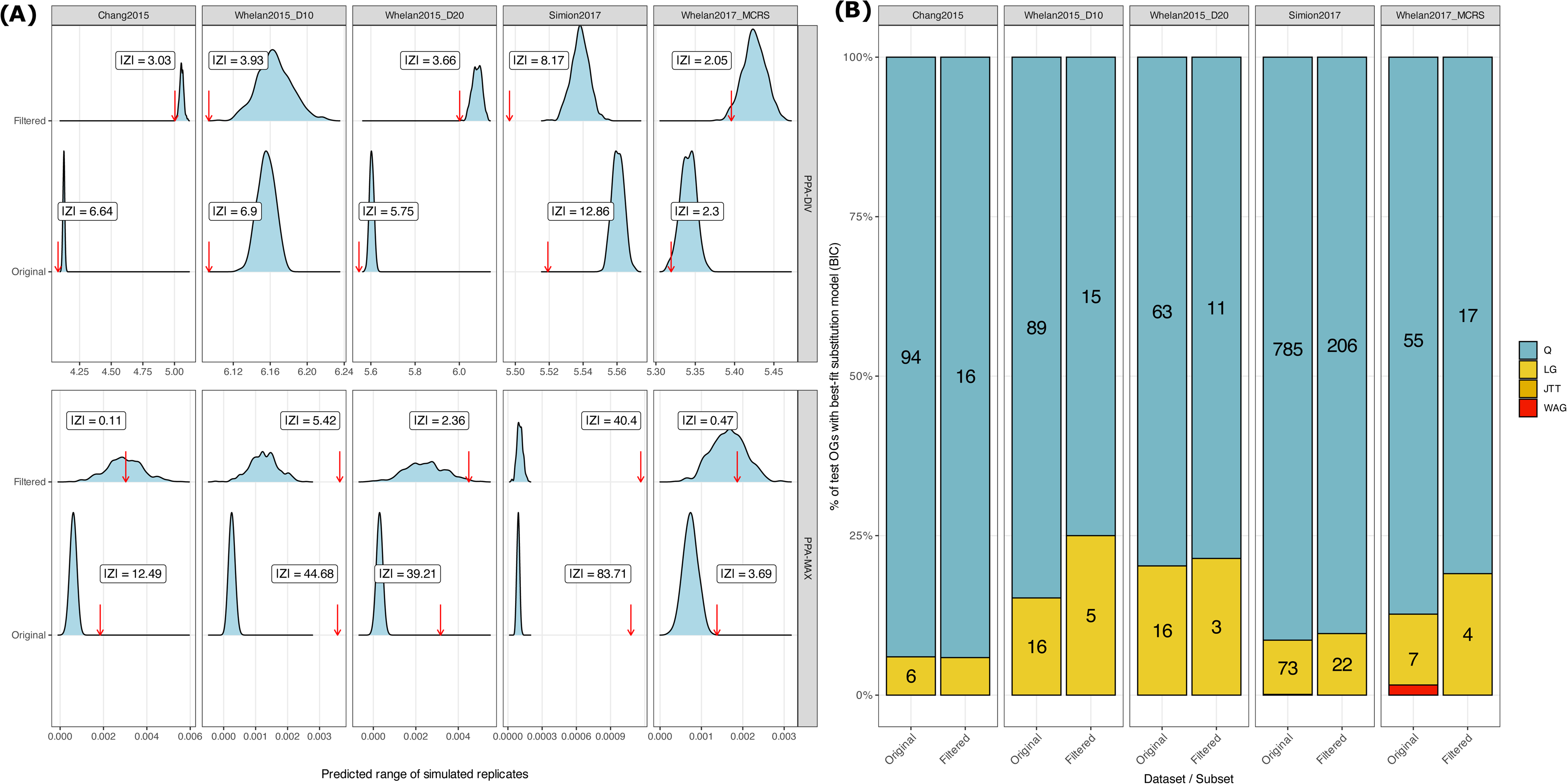
Illustration of assessment of model fit for: (a) CAT-GTR+G reconstructions of original and filtered animal phylogeny datasets. Model fit assessed using posterior predictive analysis (PPA) as implemented in PhyloBayes-MPI. Two PPA statistics shown: mean amino acid diversity (PPA-DIV) and maximal compositional heterogeneity (PPA-MAX). Red arrows indicate the observed mean for each PPA calculated for original and filtered datasets. Blue ridgeline curves represent the range of values for each PPA estimated from the CAT-GTR+G model using 500 simulated replicates, given a predicted mean and standard deviation estimated from PhyloBayes-MPI. |Z| represents the deviation of the predicted values from the observed value and hence the fit of the model to data, with the null hypothesis that the model adequately fits the data. |Z| < 2 indicates adequate model fit for a given statistic, |Z| > 5 indicates inadequate model fit. PPA data for the original Chang2015 and Whelan2015_D20 taken from Feuda et al. (2017). PPA data for the remaining original datasets generated from single chain PhyloBayes-MPI analyses run for either ∼1,000 iterations (Simion2017) or ∼5,000 iterations (Whelan2015_D10, Whelan2017_MCRS). (b) estimated empirical amino acid substitution models for original and filtered animal phylogeny datasets. Empirical amino acid substitution models (*Q*) estimated for original and filtered datasets using QMaker approach as implemented in IQTREE (Minh et al. 2020). Models estimated using 50% of orthogroups as training sets, and fit of *Q* models vs. standard empirical models (LG, WAG, JTT) assessed across remaining 50% of orthogroups as test sets under Bayesian information criterion. Stacked bar plot representing proportions of best-fit models across test orthogroups in original and filtered animal datasets, numbers of orthogroups per category given in bars (values under 3 not shown). OG: orthogroups.

**Figure 5.**
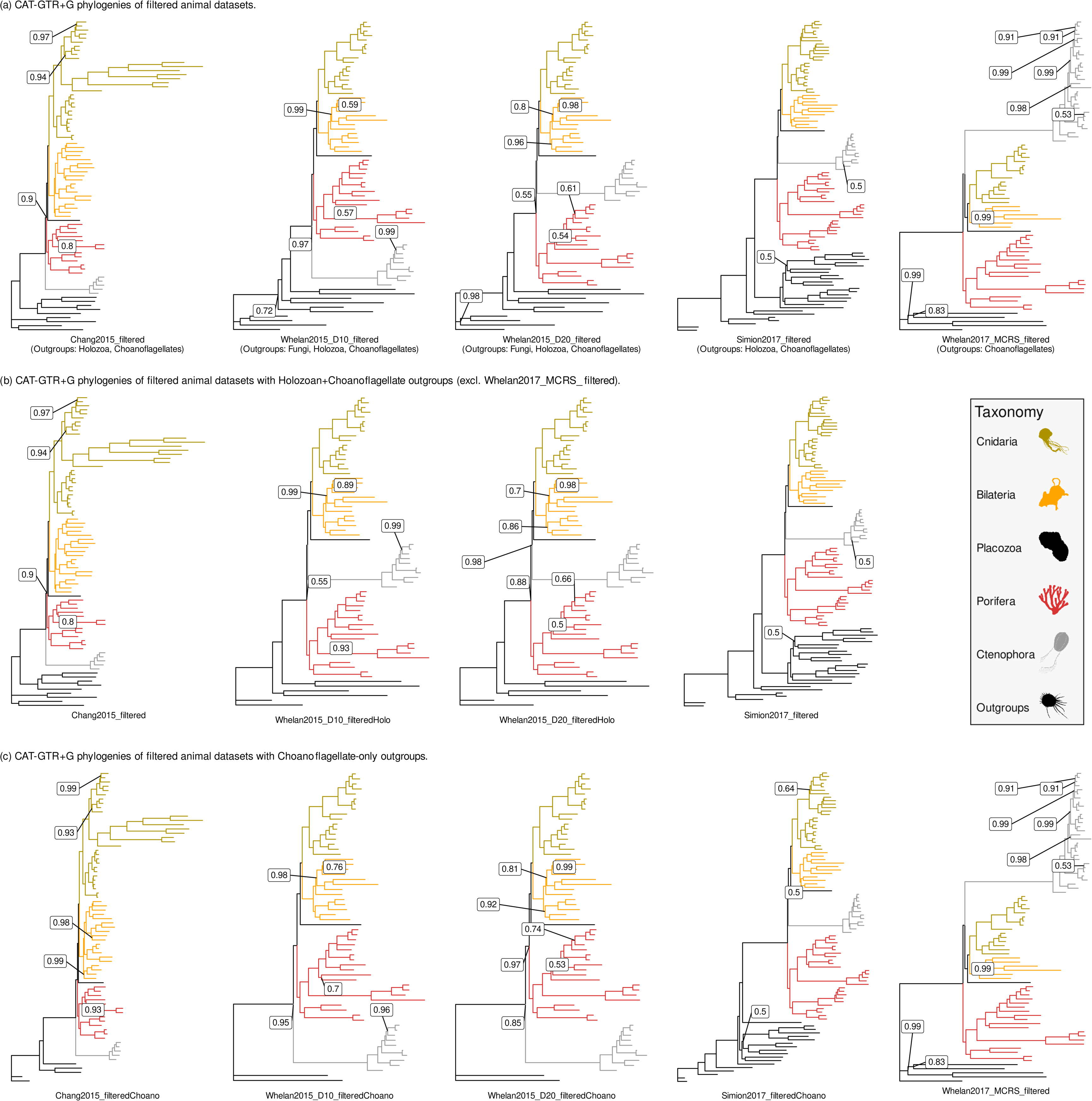
**(a**) **Bayesian CAT-GTR+G reconstructions of filtered animal phylogeny datasets.** Posterior consensus trees generated from PhyloBayes-MPI runs using bpcomp with a burn-in of 5,000 iteration and sampling every 10 iterations up to 10,000 iterations (or 7,500 iterations for Simion2017_filtered). Posterior probabilities (PP) = 1 for all branches, unless indicated. All trees rooted at most distantly-related outgroup in dataset: Ichthyosporea+Filasterea (Chang2015_filtered), Fungi (Whelan2015_D10_filtered and Whelan2015_D20_filtered), Ichthyosporea (Simion2017_filtered) or Choanoflagellata (Whelan2017_MCRS_filtered). **(b) Bayesian CAT-GTR+G reconstructions of filtered animal phylogeny datasets with outgroup sampling restricted to Holozoa and Choanoflagellata.** Whelan2017_MCRS_filtered not included in comparison due to only containing Choanoflagellata as outgroup taxa, Chang2015_filtered and Simion2017_filtered trees as in Figure 5a above. Posterior probabilities (PP) = 1 for all branches, unless indicated. All trees rooted at most distantly-related outgroup in dataset: Ichthyosporea+Filasterea (Chang2015_filtered) or Ichthyosporea (all other trees). **(c) Bayesian CAT-GTR+G reconstructions of filtered animal phylogeny datasets with outgroup sampling restricted to Choanoflagellata.** Whelan2017_MCRS_filtered tree as in Figure 5a above. Posterior probabilities (PP) = 1 for all branches, unless indicated. All trees rooted at Choanoflagellata. All silhouettes obtained from Phylopic (http://phylopic.org). Placozoa silhouette (*Trichoplax adhaerens*) by Oliver Voigt under a Creative Commons licence <u>CC BY-SA 3.0</u>; Bilateria silhouette (*Mus musculus*) by David Liao under a Creative Commons licence <u>CC BY-SA 3.0</u> and Porifera silhouette (*Siphonochalina siphonella*) by Mali’o Kodis, photograph by Derek Keats (http://www.flickr.com/photos/dkeats/) under a Creative Commons licence <u>CC BY 3.0</u>. Filozoa outgroup (*Capsaspora owczarzaki*), Cnidaria silhouette (Medusazoa sp.), and Ctenophora silhouette (*Hormiphora californensis*) under public domain.

As a complementary analysis, we also assessed model fitness in animal datasets using the QMaker approach implemented in IQTREE (Minh et al. 2020). Custom substitution matrices (*Q*) were generated for each original and filtered dataset using 50% of component orthogroups as a training set, and the fitness of each *Q* matrix was compared with three empirical substitution matrices (LG, JTT and WAG) across the other 50% of orthogroups as a test set (Minh et al. 2020). *Q* matrices were overwhelmingly selected as the best-fit model for original and filtered datasets under Bayesian information criterion, in line with observations in the Minh et al. (2020) study across different phylogenomic datasets (**Fig. 4b**), and the proportions of orthogroups selecting *Q* in original and filtered datasets is broadly similar. Following a rule of thumb from Minh et al. (2020), the high proportions of support for each *Q* matrix across filtered test set indicate that filtered animal datasets may still contain sufficient information in terms of amino acid frequencies and transition rates to conduct phylogenomic analysis, even for datasets <50 genes in size (**Fig. 4b**).

### CAT-GTR+G4 reconstruction can recover both animal root hypotheses

All initial PhyloBayes-MPI runs except for Simion2017_filtered converged in tree space upon generating posterior consensus trees (maxdiff < 0.3) (**Table 3**). For Simion2017_filtered, lack of convergence was the result of divergence between chains in resolving internal branches within Choanoflagellata and Ctenophora which became fixed early in each chain (**Supplementary Information**) and these divergences do not impact phylum-level reconstruction of the animal tree. Each filtered dataset recovered a tree which supported either the Porifera-sister or Ctenophora-sister hypothesis (**Fig. 5a, Fig. S15-S19**). An alternative “Paranimalia” hypothesis (Porifera+Ctenophora sister to all other animals) was not recovered in this analysis (Francis and Canfield 2020).

In three of the five CAT-GTR+G4 reconstructions we performed on filtered animal datasets, we recovered the same animal root as inferred in the original studies (**Fig 5a**, **Tables 1 & 3**). The Chang2015_filtered and Whelan2015D10_filtered datasets both supported a Ctenophora-sister tree (**Fig. S15-16**) (Chang et al., 2015; Whelan et al., 2015), and Simion2017_filtered supported a Porifera-sister tree (**Fig. S18**) (Simion et al., 2017). Notably, Simion2017_filtered recovers a Porifera-sister tree with holozoan outgroups (Simion et al. 2017; Feuda et al. 2017). This contradicts assumptions that recovering Porifera-sister under CAT-GTR+G4 was dependent on restricting outgroup sampling to Choanoflagellates (Halanych et al., 2016; Li et al., 2020). As for Whelan2015_D20_filtered and Whelan2017_MCRS_filtered, we recovered a Porifera-sister tree for both datasets whereas the original datasets recovered Ctenophora-sister trees (**Fig. 5a**, **Fig. S17 & S19**) (Whelan et al. 2015; Whelan et al. 2017). For Whelan2015_D20_filtered, there is noticeably poor support at the node separating Porifera from all other animals (PP = 0.55) – this may reflect gene-level discordance with the placement of Ctenophora next to the remaining animal phyla in the Bayesian tree (gCF = 0, sCF = 31) (**Figs. S6 & S17**). As with Simion2017_filtered, this represents a Porifera-sister tree obtained under CAT-GTR+G4 analysis with larger outgroup sampling. For Whelan2017_MCRS_filtered, Porifera branch closest to the five Choanoflagellate outgroup taxa (**Fig. S19**). In total two of the five datasets analyzed in this approach supported a Ctenophora-sister tree, and three supported a Porifera-sister tree (**Fig. 5a**).

Overall topologies are largely congruent between original and filtered trees for each dataset, with exceptions in some internal branches in major animal groups (see **Supplementary Information** for details on intraphylum relationships in these trees). Both Chang2015_filtered and Whelan2017MCRS_filtered fail to recover deuterostomes (Bilateria) as monophyletic, Whelan2015D10_filtered fails to resolve a singular node within demosponges (Porifera) and Simion2017_filtered deviates in the placement of stalked jellies within Cnidaria (**Supplementary Information**). Statistical supports in the filtered phylogenies are congruent with the original phylogenies with slight variations in internal branch support, and each major phyla and the relationships between phyla receives maximum or near-maximum support (**Supplementary Information**). The minor changes observed in the filtered phylogenies are likely the results of unavoidable informational loss due to our filtering approach, but the overall retention of major animal phyla and most internal relationships within these phyla indicate that this approach is phylogenetically sound.

Part of the debate surrounding the animal phylogeny root has focused on the effect of outgroup sampling strategies on root inference, particularly whether restricting outgroup sampling to Choanoflagellates induces either a Porifera-sister or Ctenophora-sister tree due to artefacts such as LBA (Halanych et al. 2016; Li et al. 2020). To examine how outgroup inclusion affects root inference for our filtered datasets, we repeated PhyloBayes-MPI runs for several datasets restricting outgroups to 1) Holozoans and Choanoflagellates (**Fig. 5b, Figs. S20-S21**) and 2) Choanoflagellates-only **(Fig. 5c, Figs. S22-S25**). For Holozoan+Choanoflagellate outgroup comparisons, Whelan2015_D10_filtered and Whelan2015_D20_filtered were reanalyzed with fungal outgroups removed and compared with all previously filtered datasets except Whelan2017_MCRS_filtered which lacked Holozoan outgroup taxa (**Fig. 5b**). These new filtered datasets (Whelan2015_D10_filteredHolo and Whelan2015_D20_filteredHolo) both support a Porifera-sister tree. Support at the node separating Porifera from Ctenophora is poor (PP = 0.55) in Whelan2015_D10_filteredHolo, improving to PP = 0.88 in Whelan2015_D20_filteredHolo (**Fig. 5b, S20-S21**). Comparing these two restricted trees with Chang2015_filtered and Simion2017_filtered, three support a Porifera-sister tree and one supports a Ctenophore-sister tree. Finally, for Choanoflagellate-only outgroup comparisons, all datasets with the exception of Whelan2017_MCRS_filtered were reanalyzed with all non-Choanoflagellate outgroups removed (**Fig. 5c**, **S22-S25**). This time, three datasets support a Ctenophore-sister tree with high statistical support (PP ≈ 0.85-1), whereas Simion2017_filteredChoano and the previously generated Whelan2017_MCRS_filtered trees support Porifera-sister (**Fig. 5c**, **S22-25**).

## Discussion

For this study, we chose animal phylogeny datasets that have supported either the Porifera-sister or Ctenophore-sister root hypothesis (**Fig. 1**) and examined the effect of enriching orthologous signal on animal root inference. Assessing orthologous signal using clan_check (Siu-Ting et al., 2019) demonstrates that orthogroups in these datasets are largely capable of recovering monophyletic Ctenophora but struggle to recover other major animal groups (**Fig. 2b**). This inconsistent orthologous signal could be due to several factors; hidden paralogy leading to ortholog misidentification, gene-species tree conflict arising from incomplete lineage sorting or divergent rates of sequence evolution driving incongruent gene trees, among others. Given the evolutionary timespan of the animals, it is possible that a combination of these confounding factors is diminishing orthologous signal in these datasets in differing proportions – i.e. hidden paralogy may drive incongruence across the tree while incomplete lineage sorting may exacerbate that incongruence in different parts of the tree. Further examination would be required to identify the driving factor(s) in this lack of signal outside of this conjecture.

Siu-Ting et al. (2019) also highlight that the use of transcriptomic data alongside genomic data in datasets like those examined in this study may exacerbate these artefacts, as transcriptomes may reflect time- and tissue-dependent complements of a genome as opposed to the full genome itself. Recent studies comparing “phylotranscriptomic” trees with phylogenomic trees have found comparable phylogenetic resolution in both, but only where transcriptomic data is of sufficient quality and sampling consistency which may be lineage-dependent (Cheon et al., 2020; Spillane et al., 2021). Improving assembly quality and the gradual standardization of data assembly protocols may help to reduce these errors in future studies (Cheon et al., 2020).

More broadly, inherent challenges in ortholog detection on kingdom-wide level may have some influence on the lack of orthologous signal we observed at major animal nodes (Natsidis et al. 2021). This is not to suggest that the datasets analyzed here have been mismanaged, as many have either been manually curated or strictly filtered to remove sources of common phylogenetic errors such as LBA or compositional bias (Chang et al., 2015; Whelan et al., 2015; Simion et al., 2017; Whelan et al., 2017). Rather, in constructing datasets intended to resolve deep nodes such as the animal root, it may be prudent to also assess genes on their ability to recover deeper relationships within a larger species tree. In one of the datasets we reanalyzed (Simion2017) internal dataset congruence was inferred from the percentage of bipartitions that were identical in individual gene trees and a corresponding species tree (Simion et al., 2017). By comparison, our approach in effect functions as a filter on congruence at individual nodes in gene trees using information derived from a species tree rather than comparing all bipartitions. For an analysis of deep nodes within a species tree, the ability of gene trees to recapitulate deeper relationships may be more relevant than their ability to recapitulate relationships closer to the tips.

Reconstructing animal phylogenies from these filtered datasets under a Bayesian CAT-GTR+G4 model recovered either Porifera-sister and Ctenophora-sister trees. No tree recovered the “Paranimalia” hypothesis, where a monophyletic Porifera+Ctenophora clade branch sister to all other animals. Francis and Canfield (2020) recovered Paranimalia from another dataset from Whelan et al. (2015) under maximum-likelihood and CAT-GTR+G4 analysis after removing sites which strongly-favoured either Porifera-sister or Ctenophora-sister, even though these sites comprised <2% of the full dataset. It is worth noting that the authors recovered the Paranimalia tree as part of a larger study addressing data management and interpretation in phylogenomic analysis, and several of the issues highlighted in their study (Francis and Canfield 2020) are relevant to this study.

Our assessment of model fit using posterior predictive analysis shows that filtered animal datasets display consistently improved fit to CAT-GTR+G4 over their original counterparts (**Fig. 5a**). This reflects the reduction in dataset size that our approach entails, but as PPA-MAX and RCFV analysis suggests our approach may indirectly reduce some data heterogeneity artefacts (**Fig 5a**, **Fig. S3**). As |Z| < 2 for many PPA statistics estimated in this study, there are still issues regarding data heterogeneity across animals datasets even with this relative improvement in fit (**Fig. 4a**, **Fig. S26**) (Feuda et al. 2017). Though CAT-GTR+G4 better accommodates heterogeneity across sites than site-homogeneous approaches, it is not designed to accommodate heterogeneity across branches (Lartillot and Philippe, 2004). Given the faster rates of evolution observed in different animal lineages (e.g. Ctenophora), this is undoubtedly a major stumbling block in accurately reconstructing animal evolutionary history (Jékely et al., 2015). Models which can better accommodate branch heterogeneity have existed for some time, but generally require computation costs infeasible on the scale of datasets analyzed here (Foster, 2004; Blanquart and Lartillot, 2008; Moran et al., 2015). The limits of current analysis using software like PhyloBayes-MPI can be observed in previous animal studies through the removal of taxa or alternative data management approach being necessary to facilitate reconstruction (Simion et al., 2017; Whelan et al., 2017). Larger datasets are not guaranteed to resolve outstanding phylogenetic conflicts any better than smaller datasets (Philippe et al., 2011; Franco et al., 2021). It may be more appropriate to prioritise smaller datasets with more careful, higher-quality gene/taxon sampling. Our complementary assessment of model fitness using the QMaker approach in IQTREE may provide some indication of the feasibility of smaller phylogenomic datasets (**Fig. 4b**) (Minh et al. 2020).

In two of the five filtered phylogenies we reconstructed under CAT-GTR+G4 we recovered a different root to that recovered by the original dataset (**Fig. 5a**). When we repeated this analysis for several datasets with different outgroups removed, we still found that it was possible to recover both Porifera-sister and Ctenophora-sister trees in varying proportions (**Fig. 5b, Fig. 5c**). Our findings indicate that at the very least it is possible to recover Porifera-sister trees under CAT-GTR+G4: 1) without applying data recoding techniques, and 2) with either Choanoflagellate or non-Choanoflagellate outgroups. This appears to contradict previous assumptions that Porifera-sister could only be recovered under recoding techniques or with certain approaches to outgroup inclusion (Halanych et al., 2016; Li et al., 2021; Hernandez and Ryan, 2021). While this study does not examine the merits or otherwise of data recoding, it appears a reasonable recourse to handling data heterogeneity if applied carefully. As for outgroup inclusion, we contend that the animal portions of these datasets may be as much of an influence on rooting the animal tree of life as the outgroup portions. In this regard, the repeated recovery of Ctenophora at greater numbers than other animal groups in all but one dataset analyzed here merits some further examination, although it should not be correlated with ultimate root inference (**Fig. 2b**). The elevated congruence of Ctenophora across these datasets is likely a reflection of faster sequence evolution amongst the comb jellies (Moroz et al. 2014), making it more likely that ctenophore taxa branch together amongst gene trees. This can also be observed by the long branch leading to Ctenophora in each phylogeny generated in this study (**Fig. 5**). It is beyond the scope of the study to assess why this might be the case, but it may be prudent to examine whether this strong congruence is a fair reflection of actual animal evolution or some unexpected bias in ortholog detection or some other step in phylogenomic dataset construction.

The question of whether the animal tree should be rooted at Porifera or Ctenophora remains to be resolved. It may be the case that we are approaching the limit of our ability to resolve this conflict with current approaches to dataset construction and sequence evolution modelling. With this in mind, we are cautious not to assert the legitimacy of one rooting over the other given our findings and the uncertainty that remains within the field (King and Rokas 2017). From a morphological perspective, the Porifera-sister hypothesis has been assumed to be the more parsimonious hypothesis as it requires fewer evolutionary changes to have occurred over the course of time (Nielsen, 2019). Other morphological characteristics may contradict this assumption however (Telford et al., 2016), and other evolutionary trends such as the independent evolution of neurons in Ctenophores and Cnidaria+Bilateria are thought to be equally possible under both hypotheses (Francis et al., 2017). There are other outstanding issues in animal phylogenomics, such as the placement of Cnidaria. Many phylogenomic studies, including those reassessed in this study, support Cnidaria+Bilateria over previous hypotheses grouping Cnidaria and Ctenophora together (Zapata et al. 2015; King and Rokas 2017). Two recent studies incorporating greater sampling of placozoan transcriptomes have supported grouping Cnidaria and Placozoa together (Laumer et al., 2018, 2019). Similarly, the position of Placozoa within animals has yet to be fully resolved (Laumer et al., 2018). In some instances protein sequence data alone may not be sufficient for resolution of problematic nodes; additional sources of information such as microRNA data have been required to resolve relationships within reptiles and placental mammals (Field et al., 2014; Tarver et al., 2016). Rare genomic events such as gene fusion/fission data may prove resourceful in clarifying some of the deeper animal relationships (Leonard and Richards, 2012; Jékely et al., 2015). Future work should seek to continue to refine current approaches whilst embracing new and complementary datatypes and methods.

## Conclusions

We examined orthologous signal across five phylogenomic datasets designed to resolve the root of the animal phylogeny as either Porifera-sister or Ctenophora-sister. Regardless of which root a dataset originally supported, we find orthogroups in these five datasets largely recover a monophyletic Ctenophora but violate most other major animal groups. We show that retaining orthogroups which can recover ≥3 major uncontroversial clans in each dataset reduces dataset size while retaining underlying phylogenetic information and taxon sampling. Bayesian CAT-GTR+G4 reconstruction of these filtered datasets recovers both root positions, in some cases supporting a different root from the original dataset. Further analysis of these datasets with outgroup restrictions applied also recovers both roots. Datasets enriched for orthologs generally exhibit better fit to the CAT-GTR+G4 model relative to the original datasets, although some data heterogeneity remains. Our findings do not definitively resolve the root of animals, but indicate that dataset size and construction can influence root inference as has been previously demonstrated for model selection and outgroup selection. These findings highlight areas of improvement in current animal phylogenomics such as dataset composition and modelling of data heterogeneity, which may help to resolve this elusive aspect of animal evolutionary history.

## Methodology

### Animal phylogeny dataset selection

Five published datasets were chosen on the basis of their differing approaches to dataset construction and support for different animal root hypotheses (Chang et al., 2015; Whelan et al., 2015; Simion et al., 2017; Whelan et al., 2017). Many of these datasets have previously been analyzed in other animal phylogeny studies (Pisani et al., 2015; Feuda et al., 2017; Shen et al., 2017). Details for each selected dataset are provided in **Table 1**, and further information on taxon sampling and construction strategies for each dataset is provided in **Supplementary Information.**

### Gene content overlap across animal phylogeny datasets

Gene content overlap across the five selected animal datasets was inferred following Francis & Canfield (2020). Where present, human sequences were extracted from all orthogroups from each dataset and queried against 20,386 human sequences from Swiss-Prot using BLASTp (e-value = 1e^-4^) (Camacho et al., 2009; Francis and Canfield, 2020; The UniProt Consortium, 2021). The sequence identifier of each top hit per seed sequence was extracted for each dataset, and was used to generate a presence-absence matrix for human Swiss-Prot sequence hits across all four datasets. This matrix was visualized as a UpSet plot representing overlap between datasets using the R package UpSetR (**Fig. S1**) (Conway et al. 2017, R Core Team 2021).

### Orthogroup alignment and tree reconstruction

Component orthogroups were extracted from their datasets using available partition information. Best-fit alignments for each orthogroup were chosen using the following procedure:

- Each orthogroup was first aligned using three different multiple sequence alignment software: MUSCLE, MAFFT and PRANK (Katoh et al., 2002; Edgar, 2004; Löytynoja, 2014). For MUSCLE and PRANK the default parameters were used for sequence alignment, and for MAFFT the most-appropriate alignment strategy was automatically selected using the “--auto” flag.
- The mutual distance between each pair of alignment methods (i.e. MUSCLE vs. MAFFT, MUSCLE vs. PRANK, PRANK vs. MAFFT) was assessed using MetAl with the default *d_pos_* metric which accounts for positional information of gaps in sequence alignments (Blackburne and Whelan, 2012). A pair of alignments with a MetAl score of <0.15 were judged to be in mutual agreement, whereas a pair with a MetAl score of >0.15 were judged to be mutually discordant.
- If any pair of alignments were judged to be mutually discordant by MetAl, the column-based normalized mean distance of amino acid similarity across all sites in all three alignments was calculated using norMD, and the alignment with the highest similarity score was selected for orthogroup tree reconstruction (Thompson et al., 2001; Muller et al., 2010; Webb et al., 2017). Otherwise if all three alignment methods were in mutual agreement, the best-fit alignment was randomly selected.

Maxmimum-likelihood (ML) reconstruction was performed for each alignment using IQTREE with automated model selection *via* ModelFinderPlus and 100 nonparametric bootstrap replicates (Nguyen et al., 2015; Kalyaanamoorthy et al., 2017).

### Filtering animal phylogeny datasets using clan_check

Clan_check examines whether user-defined sets of taxa group together as a “clans” *sensu* Wilkinson within a set of unrooted orthogroup trees (Wilkinson et al., 2007; Siu-Ting et al., 2019) (**Fig. 2a**). Orthogroups may then be excluded from data matrices based on their corresponding tree’s inability to recover a set proportion of these clans subject to the user’s own criteria (Siu-Ting et al., 2019). We defined 6 “uncontroversial” clans to be tested across all orthogroups trees: the five major animal phyla (Bilateria, Cnidaria, Ctenophora, Porifera, Placozoa) and a 6^th^ clan containing all remaining outgroup taxa. Clan composition varied between datasets, except Placozoa which was always represented solely by *Trichoplax adhaerens* (**Table S1**). Datasets were filtered to retain orthogroups which could recover ≥3 clans (Siu-Ting et al., 2019) (**Table 2**). Filtered data matrices were constructed from these orthogroups using SCaFOs and TREE-PUZZLE with the default parameters (Schmidt et al., 2002; Roure et al., 2007). Orthogroups which failed our clan_check filter were retained for comparative analysis of phylogenetic information as detailed below.

### Comparison of original and filtered animal phylogeny datasets

Several small-scale analyses were performed to assess the effect of clan_check filtering on taxon sampling, intrinsic phylogenetic information and the predicted biological composition of each dataset. These analyses are described below.

#### Taxon sampling

The distribution of taxa per clan in orthogroups passing or failing our clan_check filter were visualized for each dataset as boxplots using the R package ggplot2, with significance assessed using Wilcoxson tests (p < 0.05) (Wickham, 2009) (**Fig. S2**).

#### Data heterogeneity, branch length and relative compositional frequency

Wilcoxson tests (p < 0.05) were performed in R (R Core Team 2021) to assess differences in branch length and compositional heterogeneity in orthogroups passing or failing our clan_check filter, using information taken from IQTREE runs (Siu-Ting et al., 2019) (**Table S2**). Relative compositional frequency values (RCFV), another measure of compositional bias, were calculated per orthogroup for each dataset using BaCoCa (Zhong et al., 2011; Kück and Struck, 2014). The distribution of RCFV values across orthogroups passing or failing our clan_check filter was visualized using ggplot2, with significance assessed using a Wilcoxson test (p < 0.05) (Wickham, 2009) (**Fig. S3**). Chi-squared tests for compositional homogeneity across all datasets and their component orthogroups were performed using p4 (Foster 2004).

#### Gene ontology categories

Predictive sequence annotation of all human sequences from each animal dataset was performed using InterProScan (Jones et al., 2014). Gene ontologies (GOs) were extracted for sequences from orthogroups passing or failing our clan_check filter in each dataset. GOs were grouped into three major categories - “Cellular Component”, “Molecular Function” and “Biological Process” - using the Python package GOATools (Klopfenstein et al., 2018). Pearson’s chi-square tests of independence (p < 0.05) was performed in R (R Core Team 2021) between GO categories in orthogroups passing or failing the clan_check filter for each dataset (**Table S3**) (Siu-Ting et al., 2019).

#### Evaluation of phylogenetic information

Assessment of information content and biases in each original dataset was performed using PhyKIT (Steenwyk et al., 2021). Several metrics associated with phylogenetic information quality, heterogeneity and other biases were tested for all orthogroups and/or orthogroup trees in each dataset: alignment length, mean bipartition support, mean branch length, parsimony-informative sites, sequence saturation, variable sites and treeness divided by relative compositional values (RCV) (Steenwyk et al., 2021). Boxplot graphs comparing orthogroups which passed our clan_check filter with those that failed the filter were generated for each metric using ggplot2, with significance assessed using a Wilcoxson test (p < 0.05) (Wickham, 2009) (**Fig. 3**).

### Gene and site concordance factor analysis

Gene and site concordance factor analysis was performed on original and filtered animal datasets (Minh et al. 2020). For a given species tree *T* and a set of gene trees *S*, the gene concordance factor (gCF) of each branch *x* in *T* can be calculated as the number of trees in *S* containing a branch concordant with *x*, divided by the number of trees in *S* decisive for *x* (i,e, trees that are capable of being either concordant or discordant with respect to *x*). Similarly, the site concordance factor (sCF) for each branch *x* in *T* is calculated as the number of sites in randomly-sampled subalignments of a larger data matrix in concordance with *x* divided by the number of decisive sites for *x*. Gene and site concordance factor analysis was performed using all original and filtered phylogenies (with manual resolution of some polytomies) using IQTREE (Nguyen et al. 2015; Minh et al. 2020). The results of concordance factor analysis for each set of original and filtered dataset were visualized along the branches of each original and filtered phylogeny using ggtree, and the distribution of gCF vs. sCF values per phylogeny were visualized as a scatterplot using ggplot2 (**Figs. S4-S8**) (Wickham 2009; Yu et al. 2017).

### Bayesian CAT-GTR+G4 reconstruction of filtered datasets

Bayesian CAT-GTR+G4 reconstruction was performed using PhyloBayes-MPI with two independent chains and removal of constant sites (Lartillot et al., 2013). PhyloBayes-MPI was run for at least 10,000 iterations on both chains for four filtered datasets, and 7,500 iterations for Simion2017_filtered due to dataset size and computational limitations. Quantitative assessment of Bayesian chain covergencce was performed using tracecomp with a burn-in of 5,000 iterations (**Table 3**). Visual assessment of chain convergence was performed using two approaches derived from https://github.com/wrf/graphphylo (Francis, 2018). First, trace plots of both chains were generated and visually assessed for patterns of parameter convergence using the plot_phylobayes_traces.R script from https://github.com/wrf/graphphylo**(Figs. S9-S13**).

Second, all-vs-all pairwise Robinson-Foulds distances (Robinson and Foulds, 1981) were calculated for all trees from each chain using RAxML and normalized RF distances were plotted using scripts from https://github.com/wrf/graphphylo and ggplot2 (Wickham, 2009; Stamatakis, 2014) (**Fig. S14**).

Posterior consensus trees were generated using bpcomp with a burn-in of 5,000 iterations and sampling every 10 iterations up to 10,000 iterations (or 7,500 iterations for Simion2017_filtered) (Lartillot et al., 2013). The same procedure was repeated for all filtered animal datasets with additional outgroup restriction: all datasets where outgroup sampling was restricted to Holozoa and Choanoflagellata (Whelan2015_D10_filteredHolo and Whelan2015_D20_filteredHolo) and all datasets where outgroup sampling was restricted to Choanoflagellata (Chang2015_filteredChoano, Whelan2015_D10_filteredChoano, Whelan2015_D20_filteredChoano and Simion2017_filteredChoano). For all filteredHolo / filteredChoano datasets except Simion2017_filteredChoano, PhyloBayes runs were performed up to 10,000 iterations with posterior consensus trees generated as previously described. For Simion2017_filteredChoano, computational and time limitations restricted PhyloBayes-MPI analysis to approximately 3,500 iterations on both chains, with a posterior consensus tree generated using bpcomp with a burn-in of 1,500 iterations and sampling every 10 iterations. All phylogenies were visualized and annotated using ggtree (Yu et al. 2017) (**Fig. 5, Figs. S15-25**).

### Assessment of model fit using posterior predictive analysis

Posterior predictive analysis (PPA) was performed to assess model fit for each PhyloBayes-MPI run (Bollback, 2002; Lartillot and Philippe, 2004; Feuda et al., 2017; Lartillot et al., 2013). Five PPA statistics were tested; three (PPA-DIV, PPA-CONV, PPA-VAR) assess modelling of site-specific heterogeneity (Lartillot et al., 2007; Feuda et al., 2017), and two (PPA-MAX, PPA-MEAN) assess modelling of lineage-specific heterogeneity (Blanquart and Lartillot, 2008). Model fit was quantified by computing the absolute Z-score for each statistic on observed and simulated data, where |Z| represents standard deviations of the simulated data from the observed mean. For each statistic, |Z| < 2 indicated adequate model fit whereas |Z| > 5 indicated that the model cannot adequately fit the data (Feuda et al., 2017; Lartillot, 2020). Posterior predictive analysis was performed for each run using the “--allppred” flag in readpb_mpi, with a burn-in of 5000 iterations and sampling every 10 iterations (Lartillot et al., 2013).

PPAs obtained from the filtered datasets were compared with those for the original datasets to assess possible improvements in model fit arising from our clan_check filtering approach. PPA results for the original Chang2015 and Whelan2015_D20 datasets were obtained from Feuda et al. (2017). For Whelan2015_D10 and Whelan2017_MCRS, PhyloBayes-MPI analysis was performed as above on a single chain run for at least 5,000 iterations to generate enough potential replicates representative of each dataset. PPA results were then obtained for these original datasets as previously described above, using a burn-in of 1,000 iterations and sampling every 10 iterations. For Simion2017, a single chain was run for at least 1,000 iterations due to computational limitations and PPA results were obtained using a burn-in of 500 iterations with sampling every 10 iterations. The observed mean, predicted mean and standard deviation of each PPA statistic was used to generate ridgeline density plots using ggridges and ggplot2 (**Fig. 4a, Fig. S26**) (Wickham, 2009).

### Assessment of model fit using estimated empirical substitution models

An alternative approach to assessing the effect of our clan_check filtering approach on model fit was conducted using the maximum-likelihood QMaker approach as implemented in IQTREE, which facilitates the generation of empirical amino acid substitution models directly from phylogenomic data matrices (Nguyen et al. 2015; Minh et al. 2020). Custom substitution models were estimated for each original and filtered animal dataset using the following procedure:

- For a given dataset, 50% of component orthogroups were grouped into a “training” set and the other 50% were grouped into a “test” set.
- Initial best-fit models were estimated for each orthogroup in the training set from a candidate set of LG, WAG and JTT with a single edge-linked tree across all orthogroups and rate heterogeneity limited to four categories (Duchêne et al. 2020).
- From these initial estimations, a joint time-reversible substitution matrix *Q* was estimated across all orthogroups in the training set.

Assessment of model fit was then ran for the “test” set of each original and filtered dataset in IQTREE, restricting model selection in ModelFinderPlus to *Q*, LG, WAG and JTT (Kalyaanamoorthy et al. 2017; Minh et al. 2020). The best-fit model, according to Bayesian information criterion, for each orthogroup in the test sets of all original and filtered datasets was tabulated and visualised as a stacked bar plot using ggplot2 (**Fig. 5b**) (Wickham 2009).

## Supplementary information

Data and scripts used in this study can be found at https://github.com/chmccarthy/ATOLRootStudy

## Acknowledgements

MJO’C would like to thank both the University of Leeds for her 250 Great Minds Fellowship career award and the University of Nottingham for funding for this project through CGPM and POM. CGPM, POM and MJO’C are grateful for access to the University of Nottingham’s Augusta HPC service. CJC wishes to acknowledge funding from the BBSRC (BB/R019185/1) and DAFM Ireland/DAERA Northern Ireland (R3192GFS) and both CJC and KS-T from the EC, Horizon 2020 (818368, MASTER). This work was supported by a fellowship from the Irish Research Council–Marie Sklodowska-Curie cofund program (ELEVATEPD/2014/69 to KS-T).

## Author contributions

MJO’C conceived the project and directed the work. CGPM and POM performed all data analyses with input from MJO’C. MJO’C, CGPM, POM, KST and CJC contributed to experimental design and interpretation of results. CGPM and MJO’C wrote the manuscript and all authors contributed to editing.

---

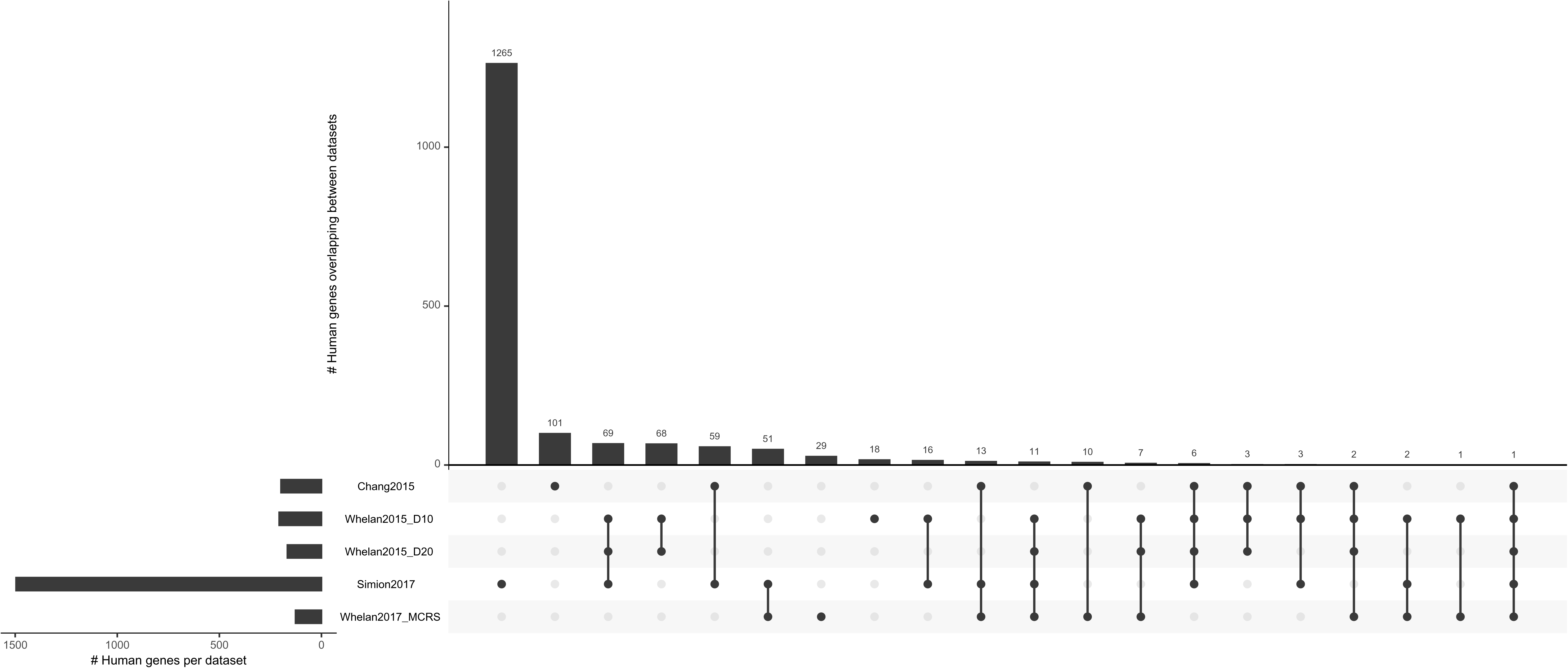
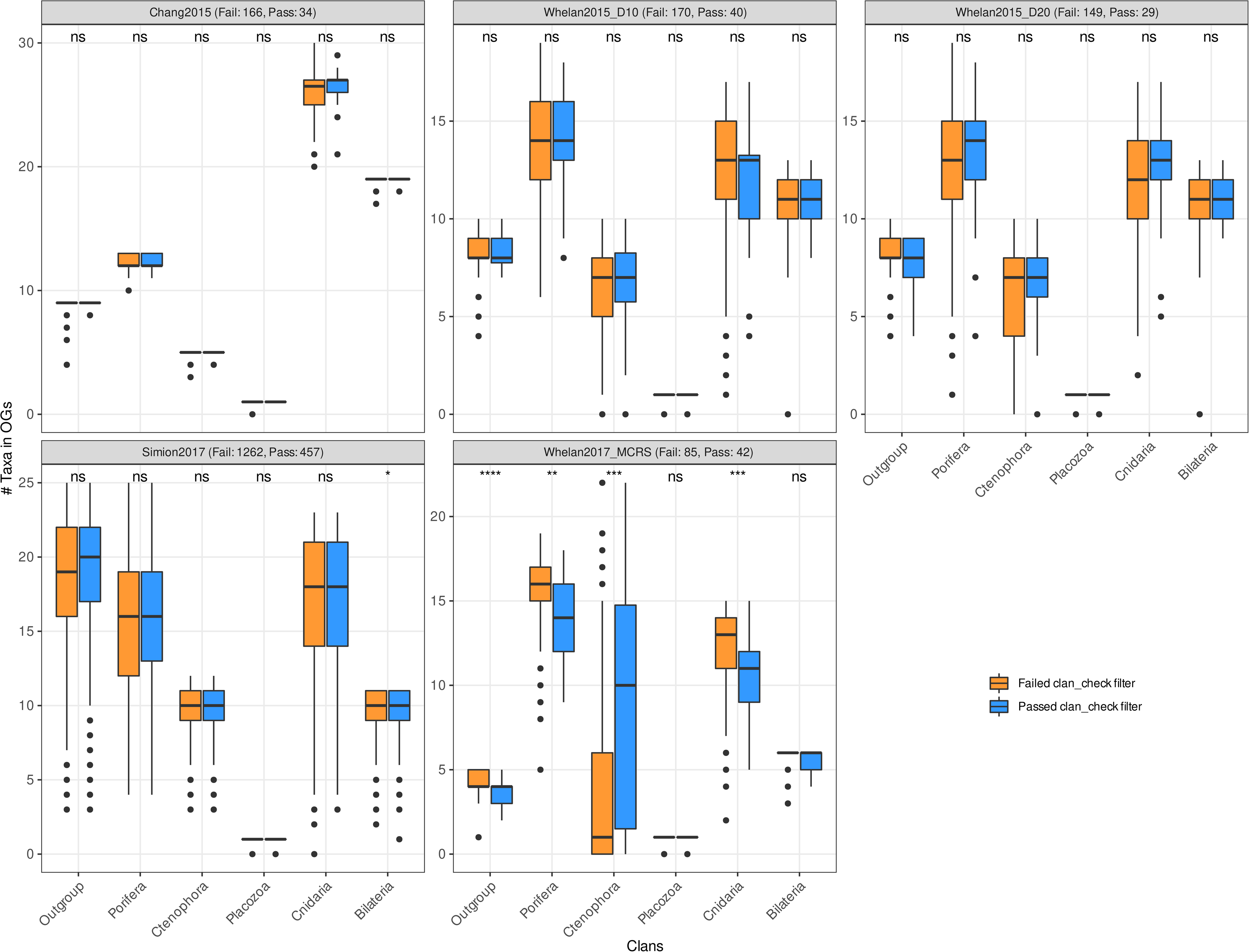
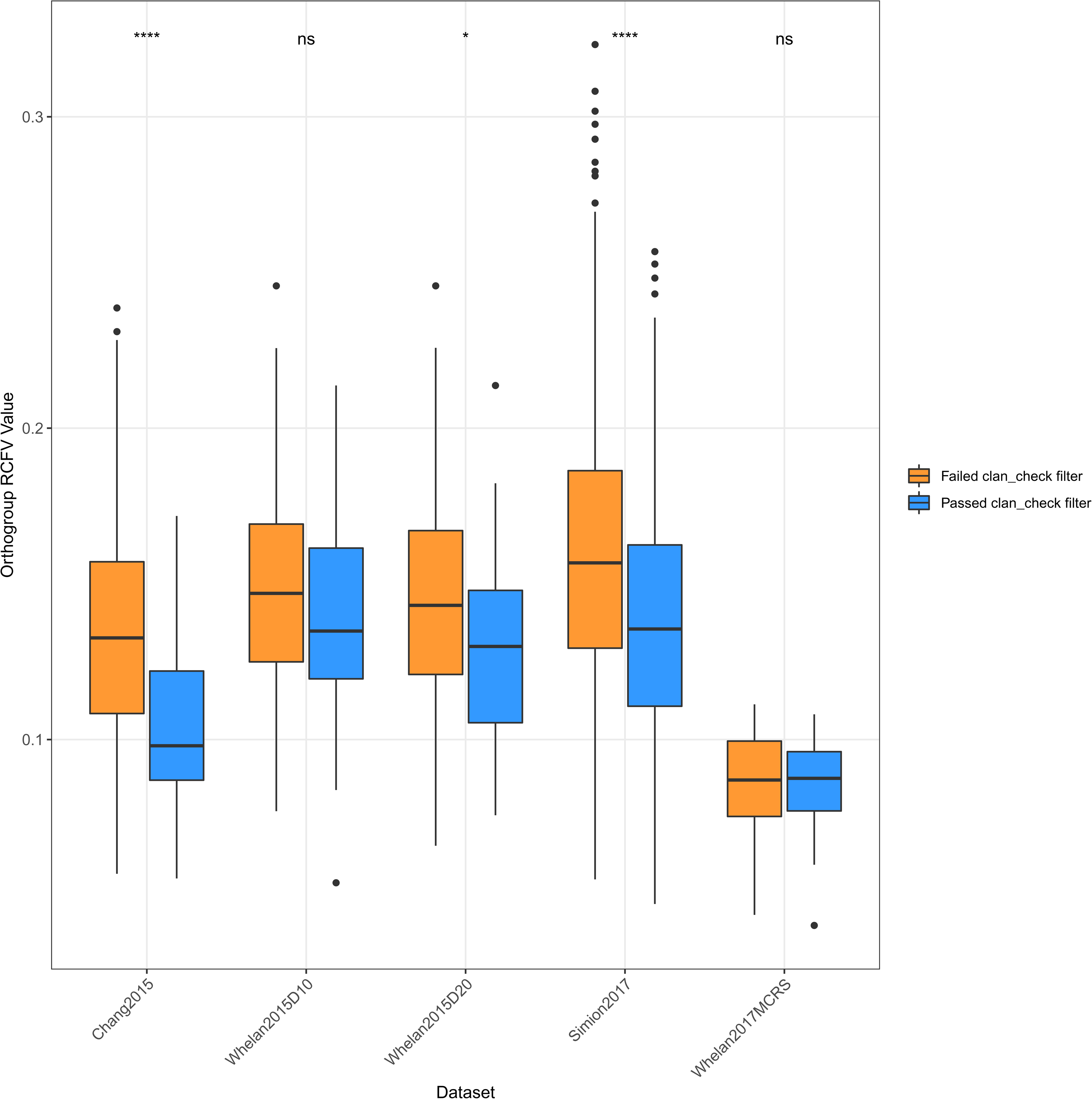
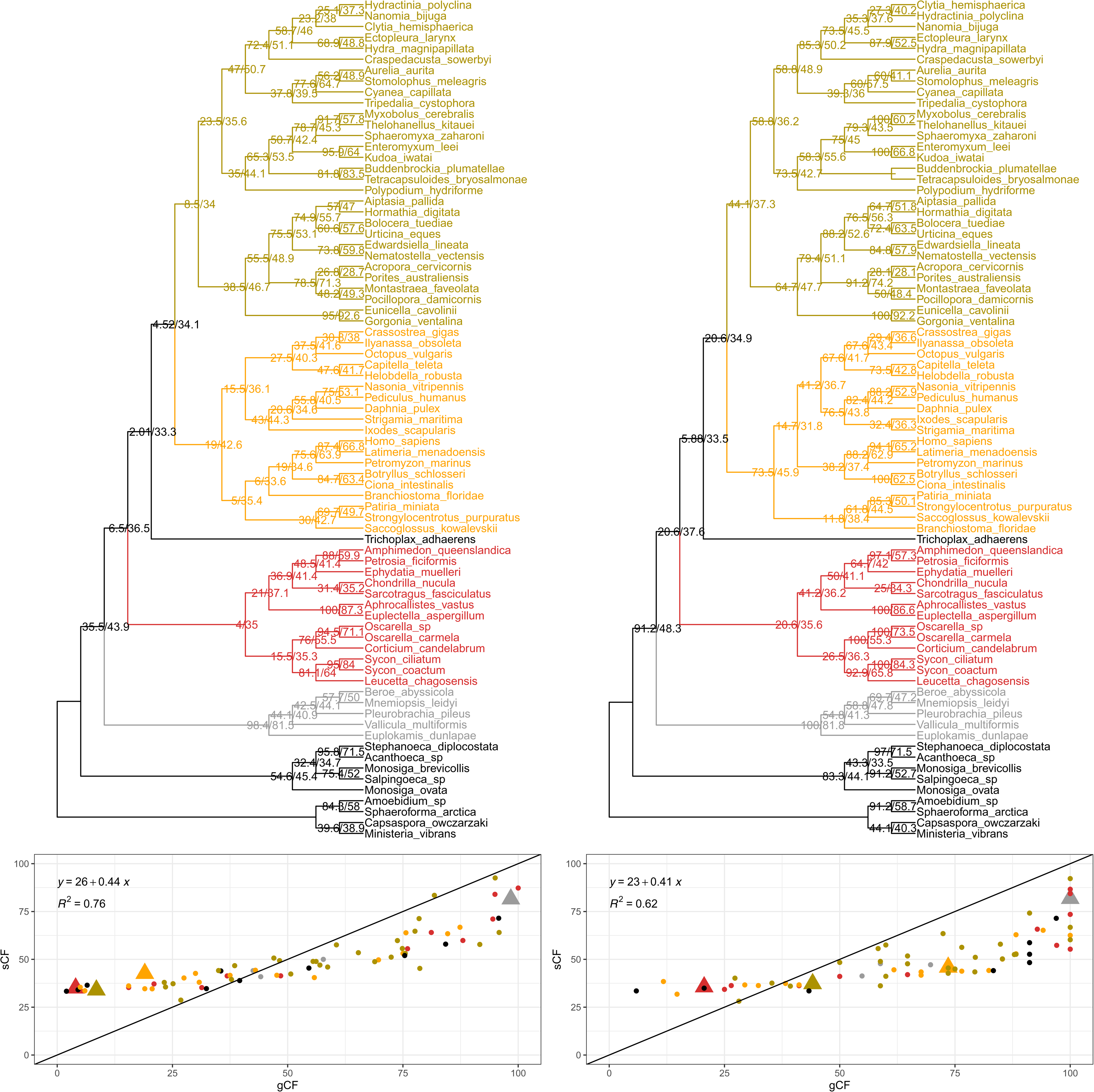
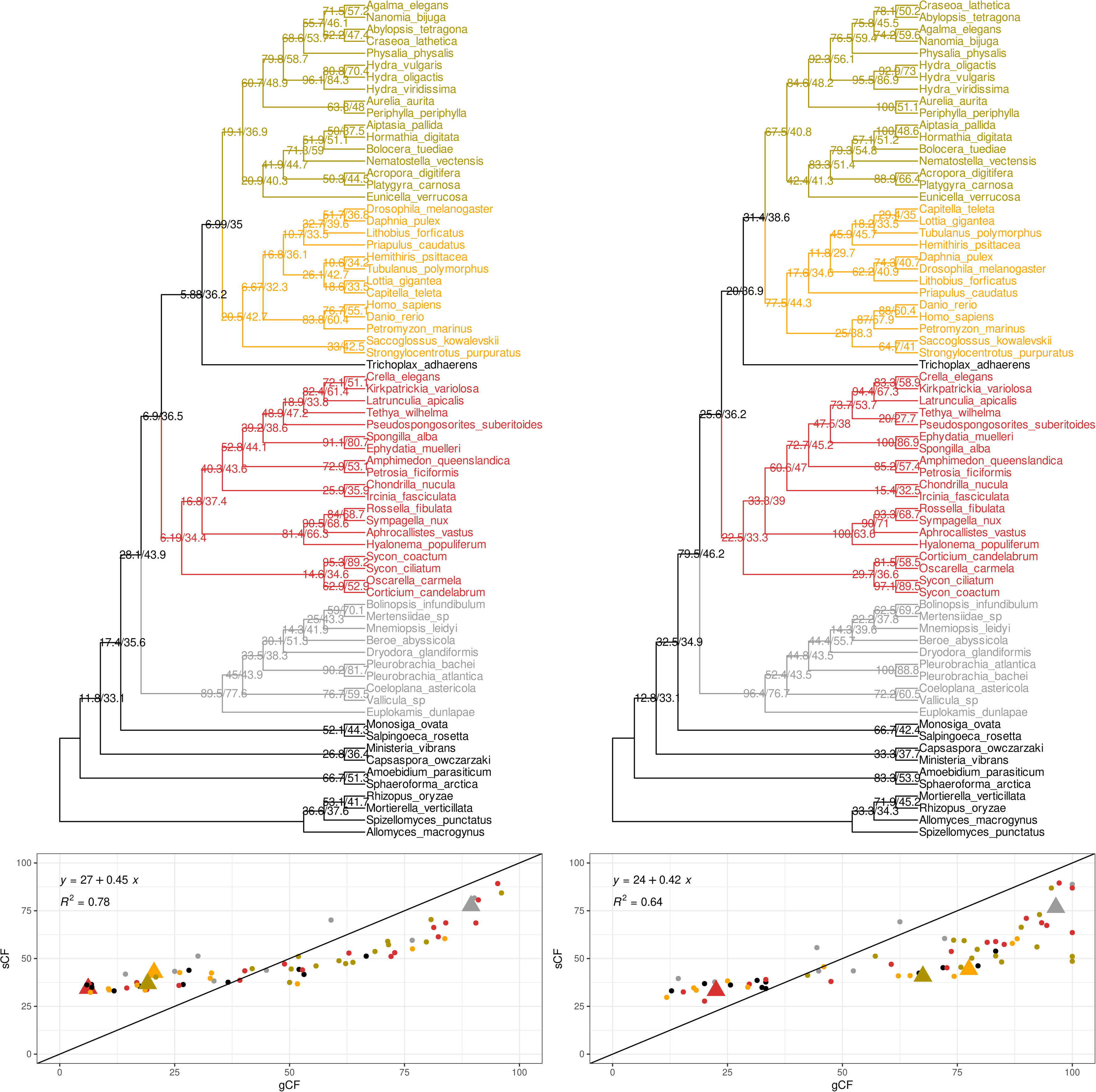
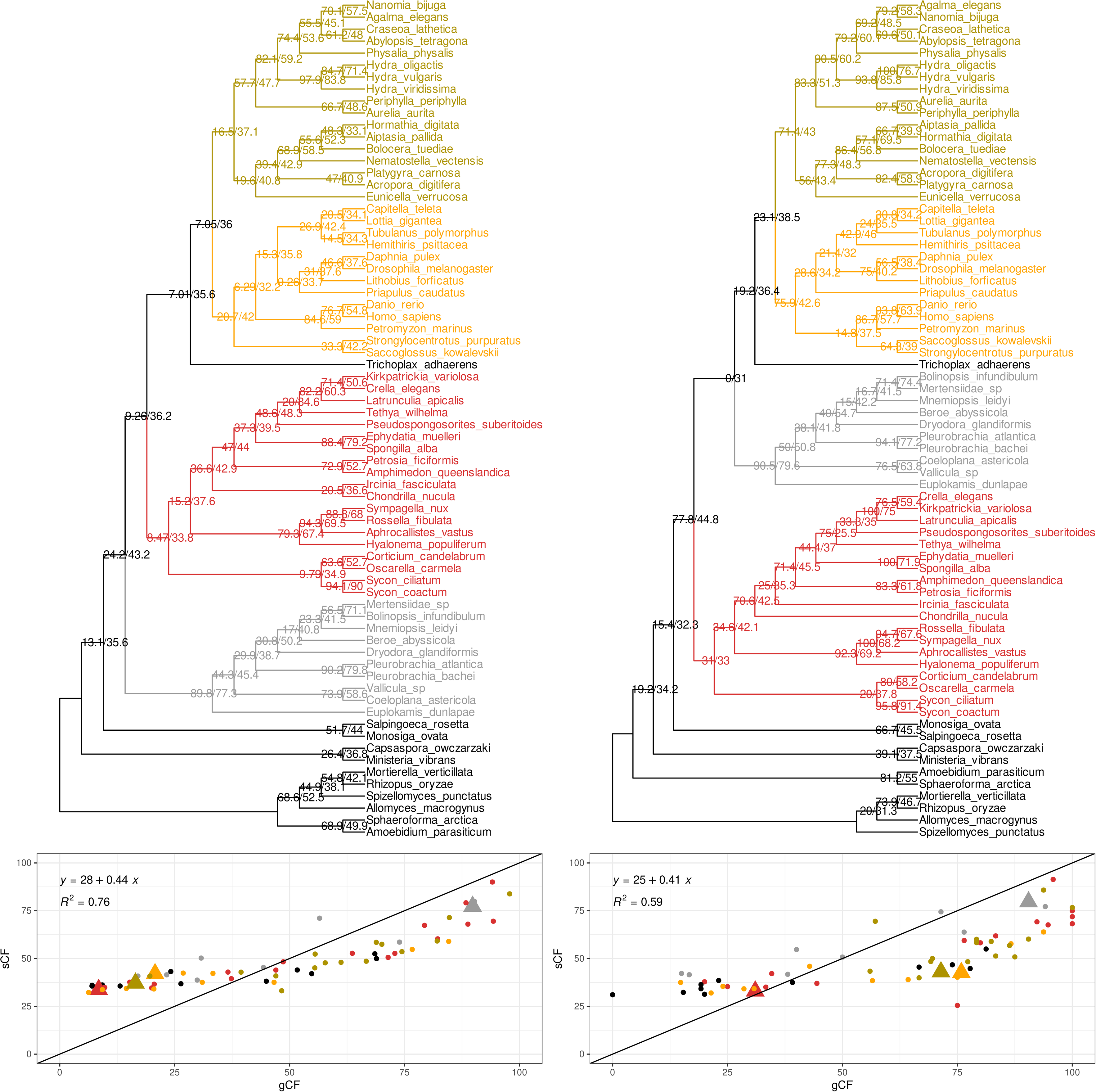
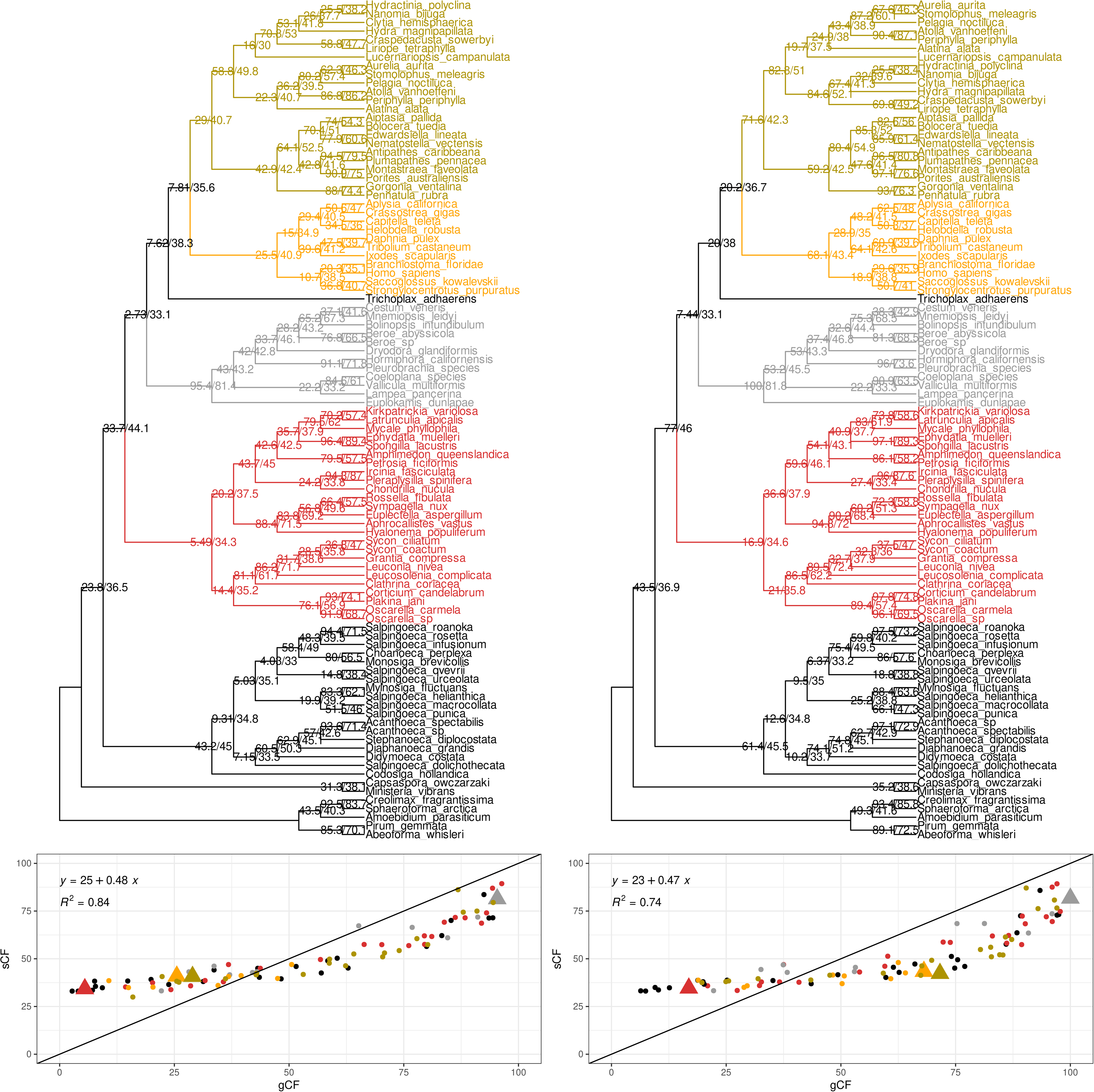
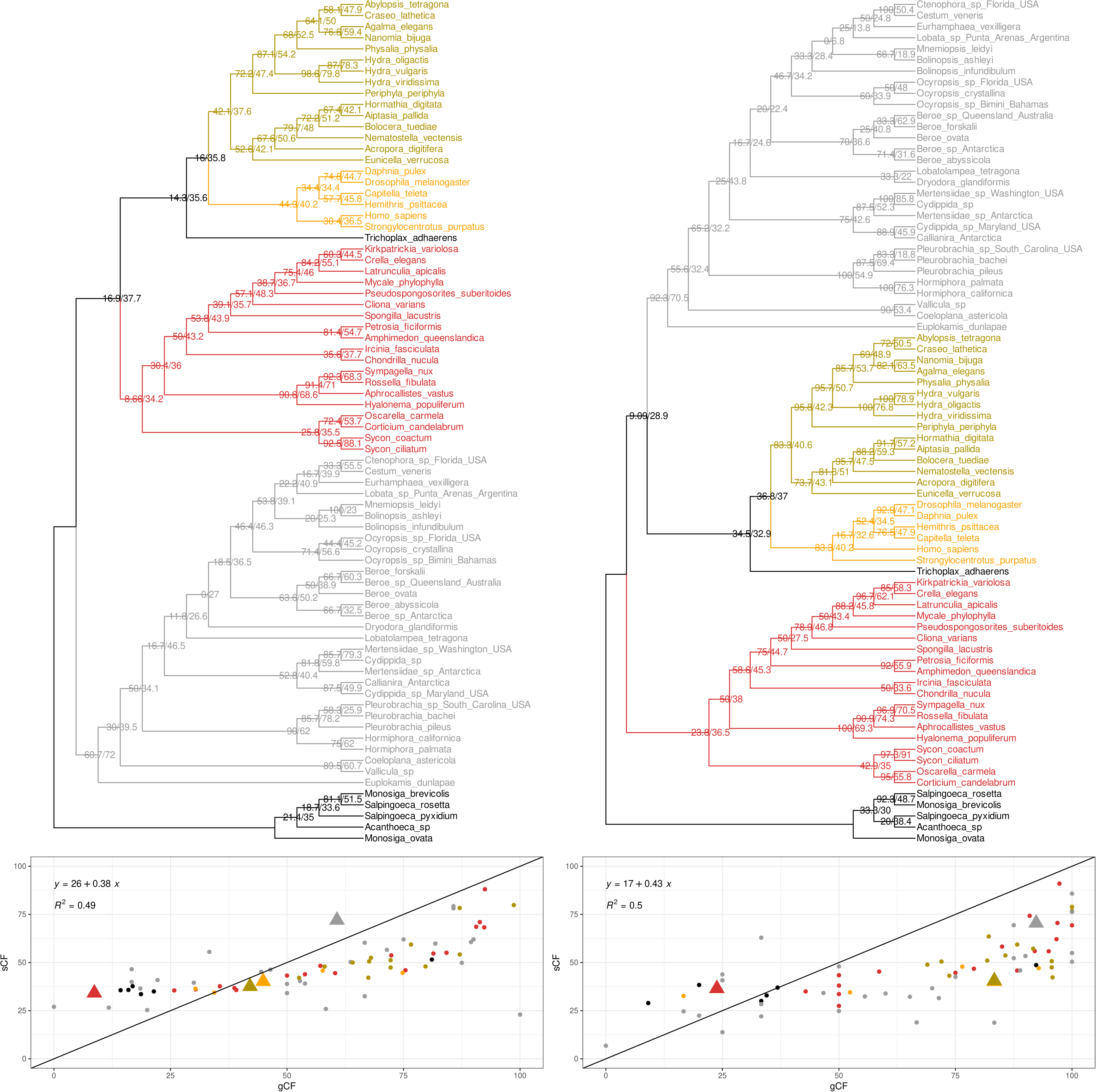
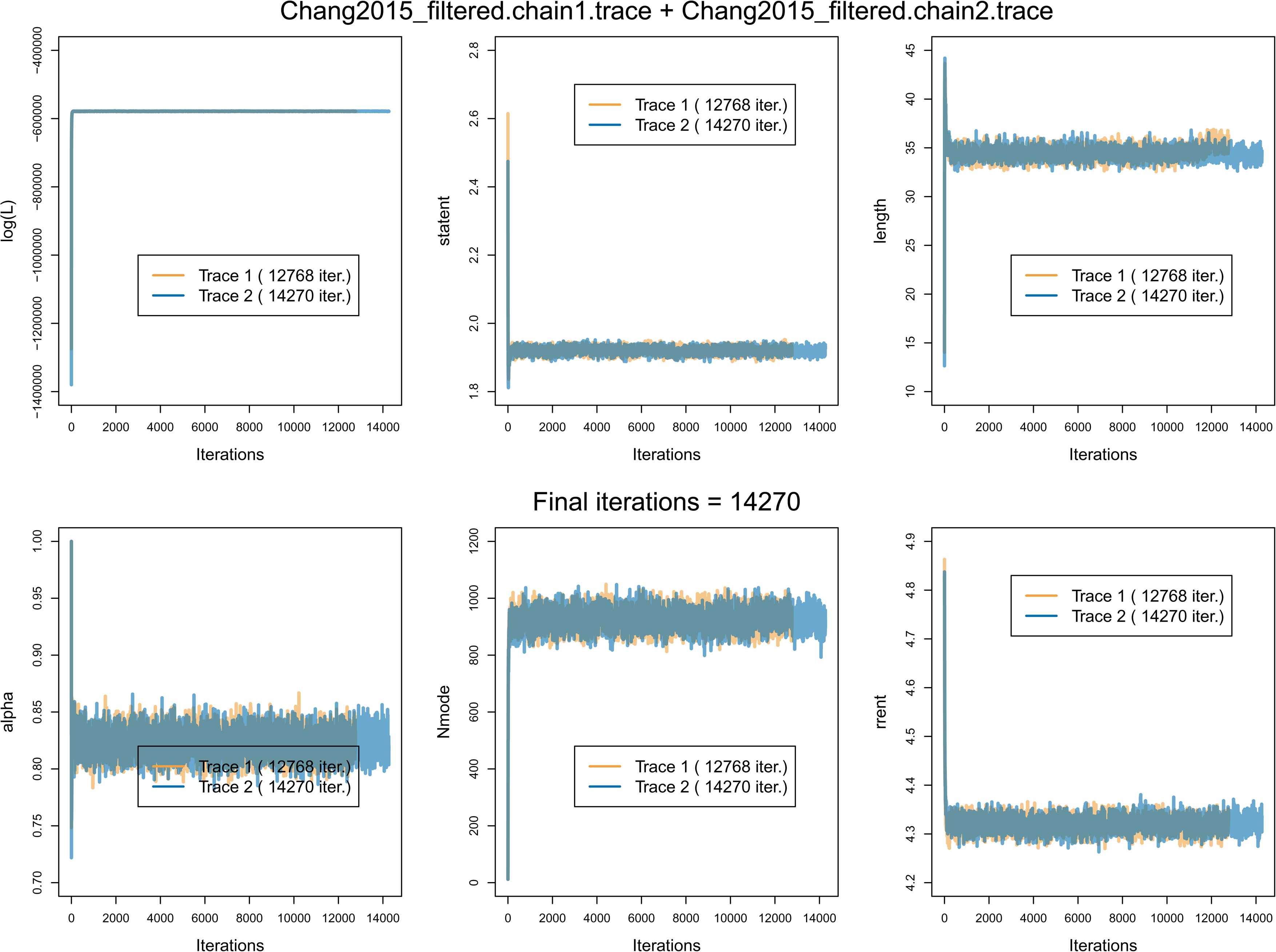
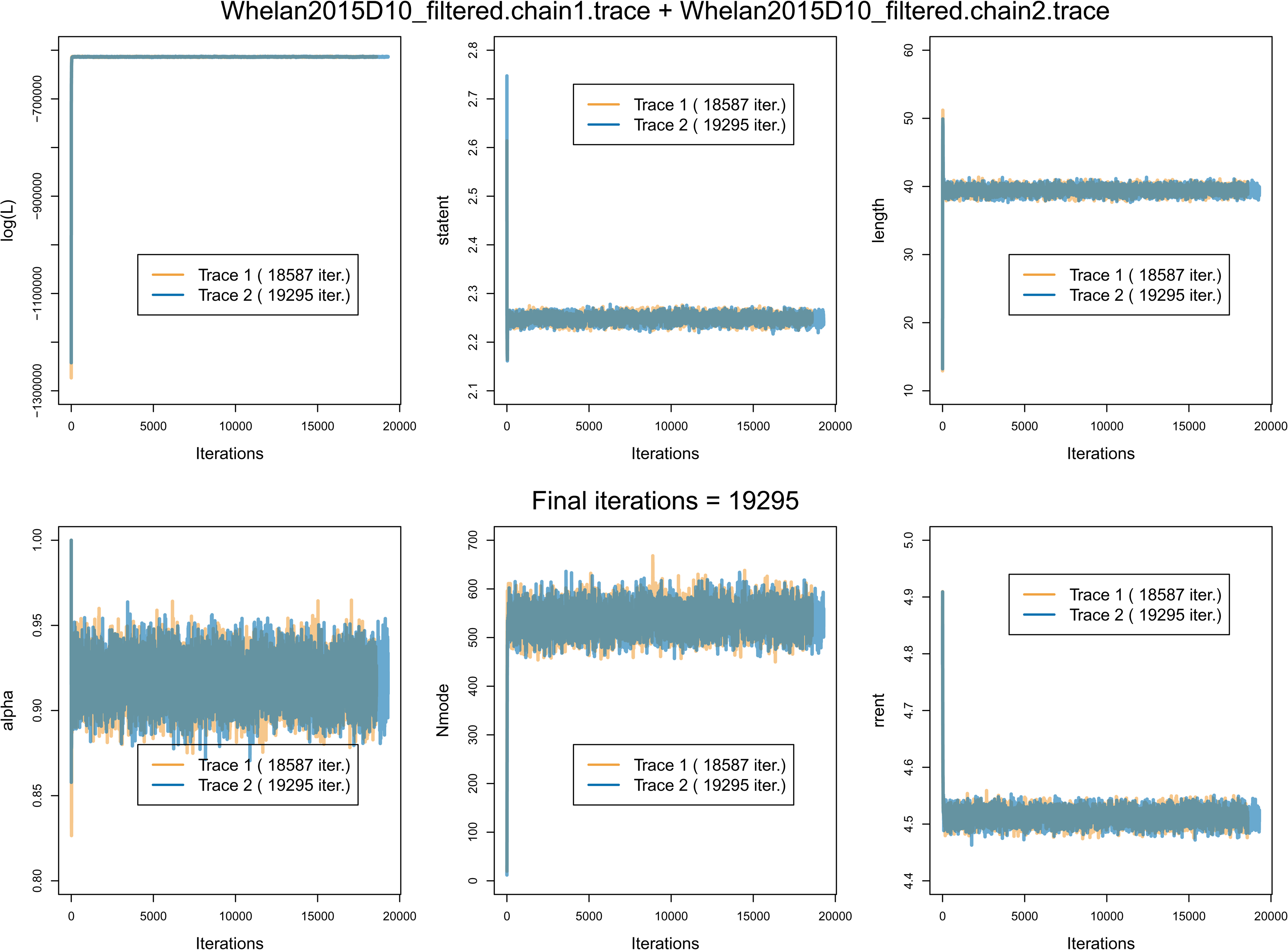
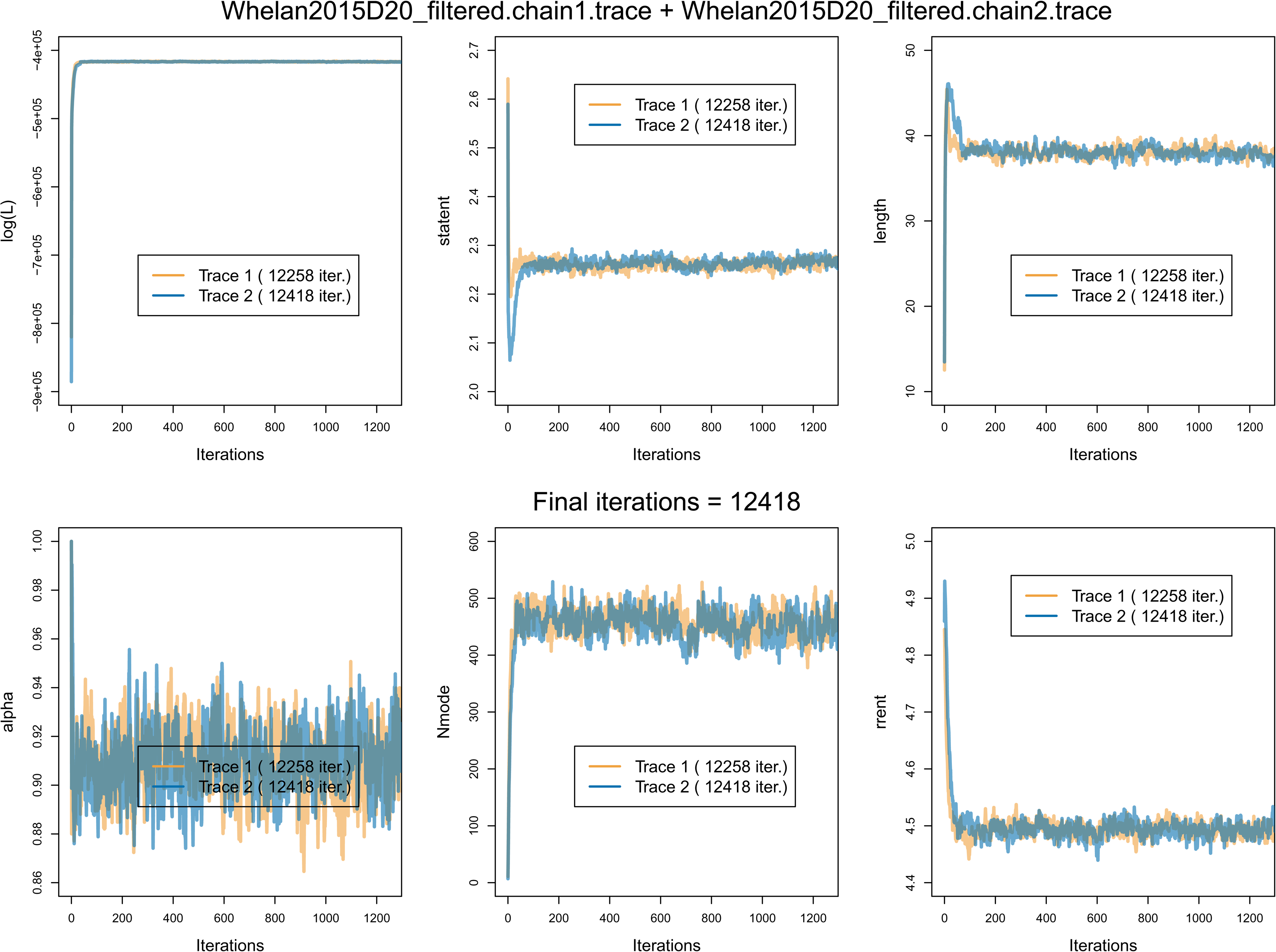
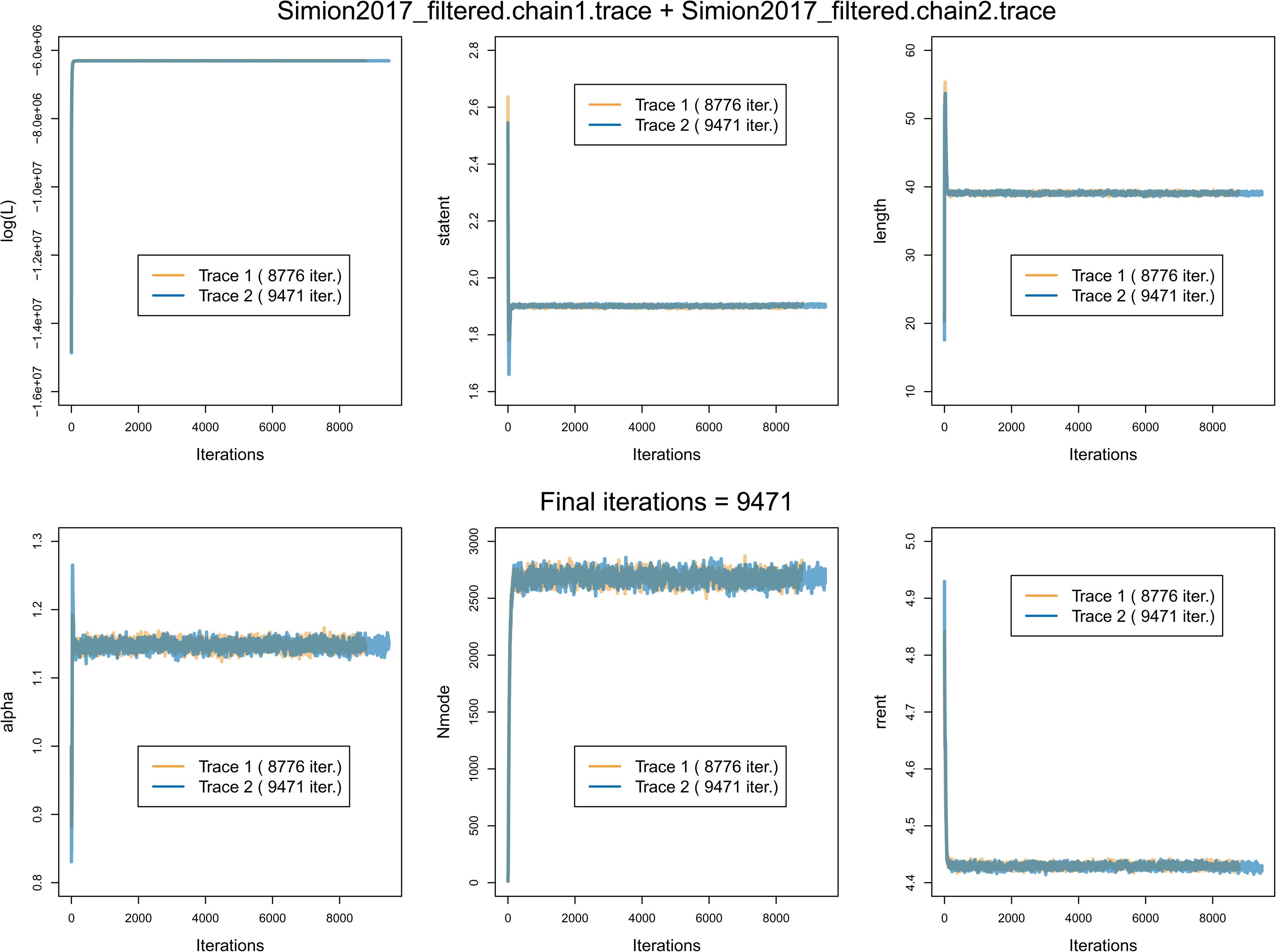
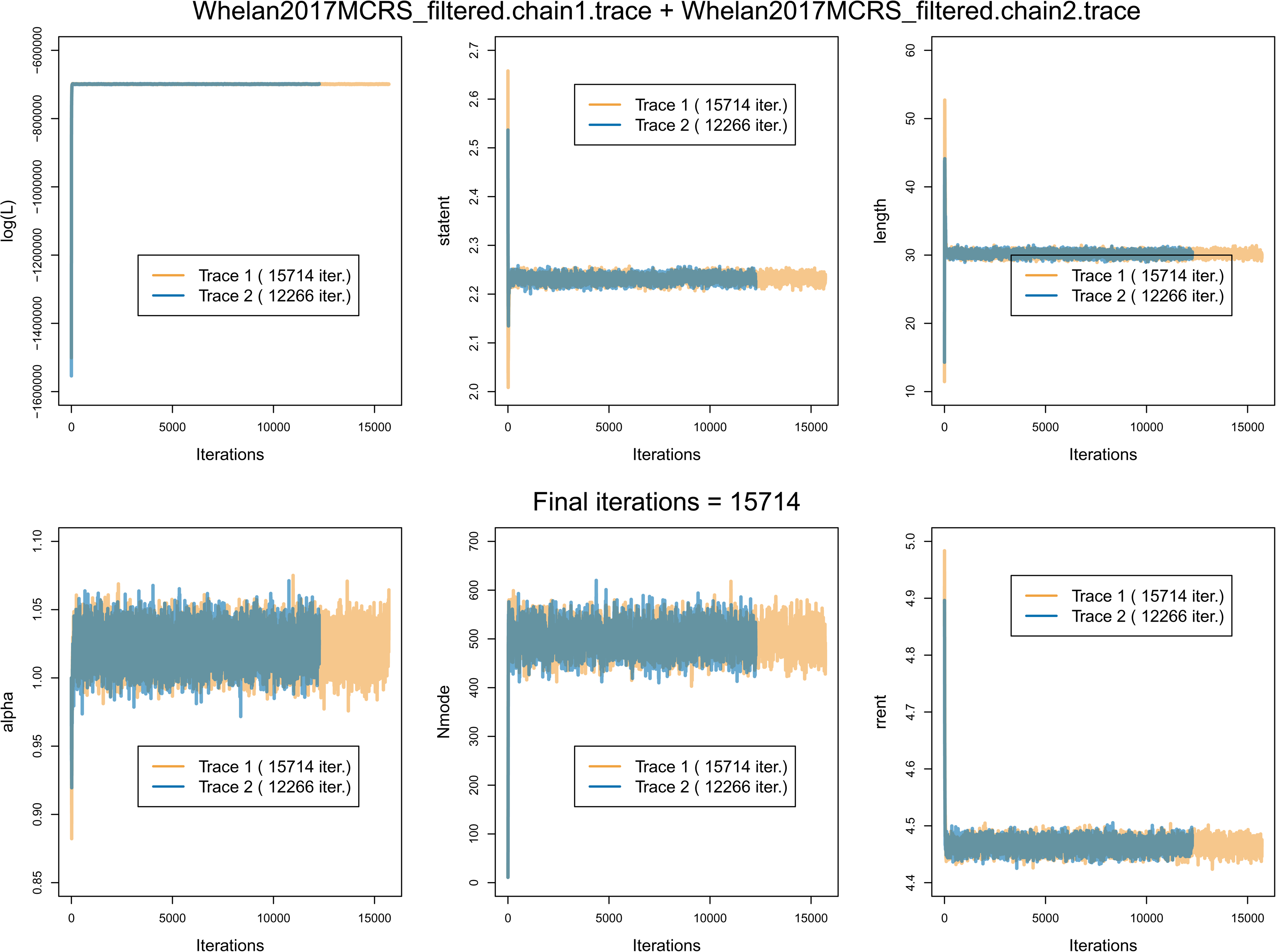
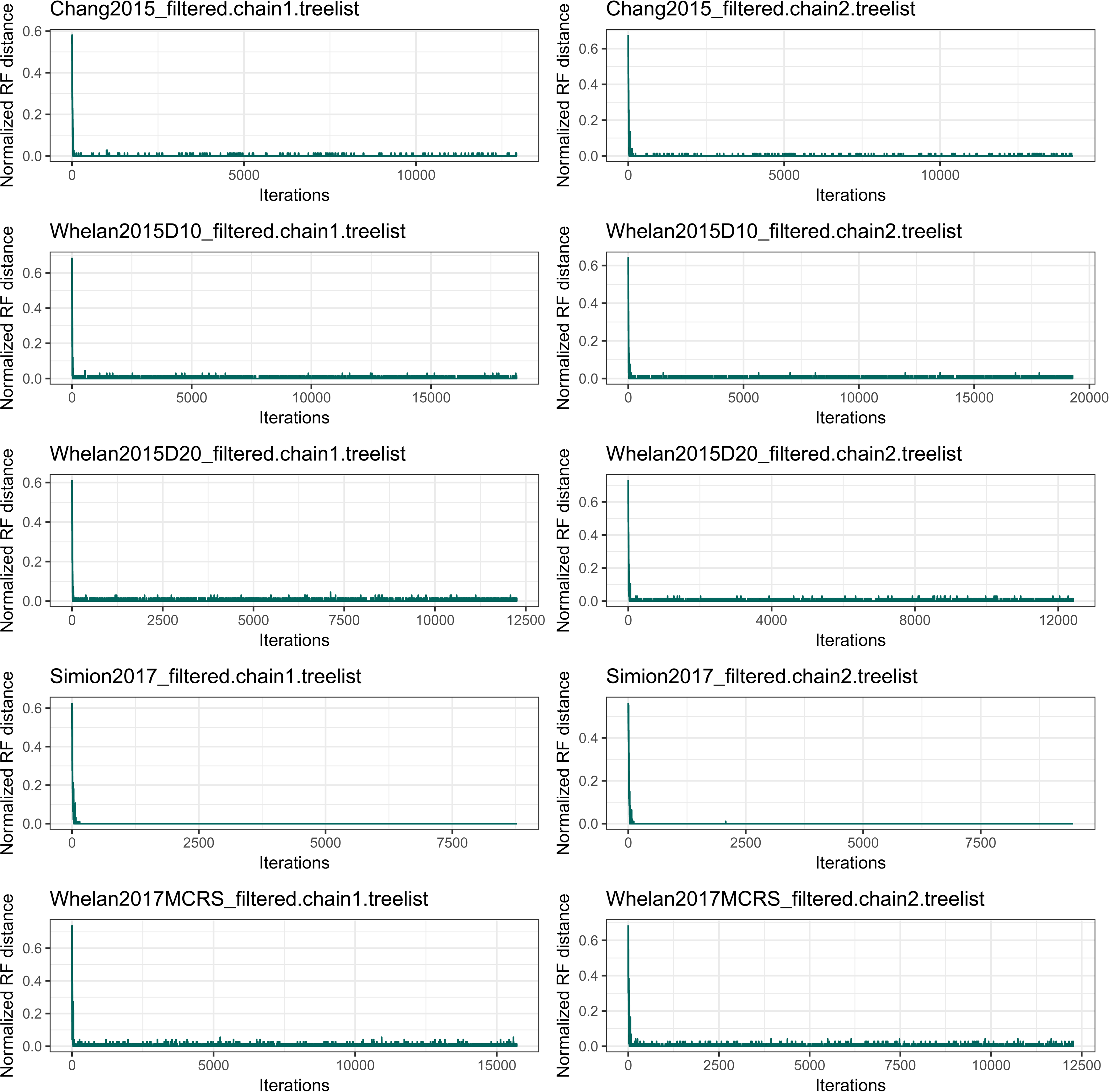
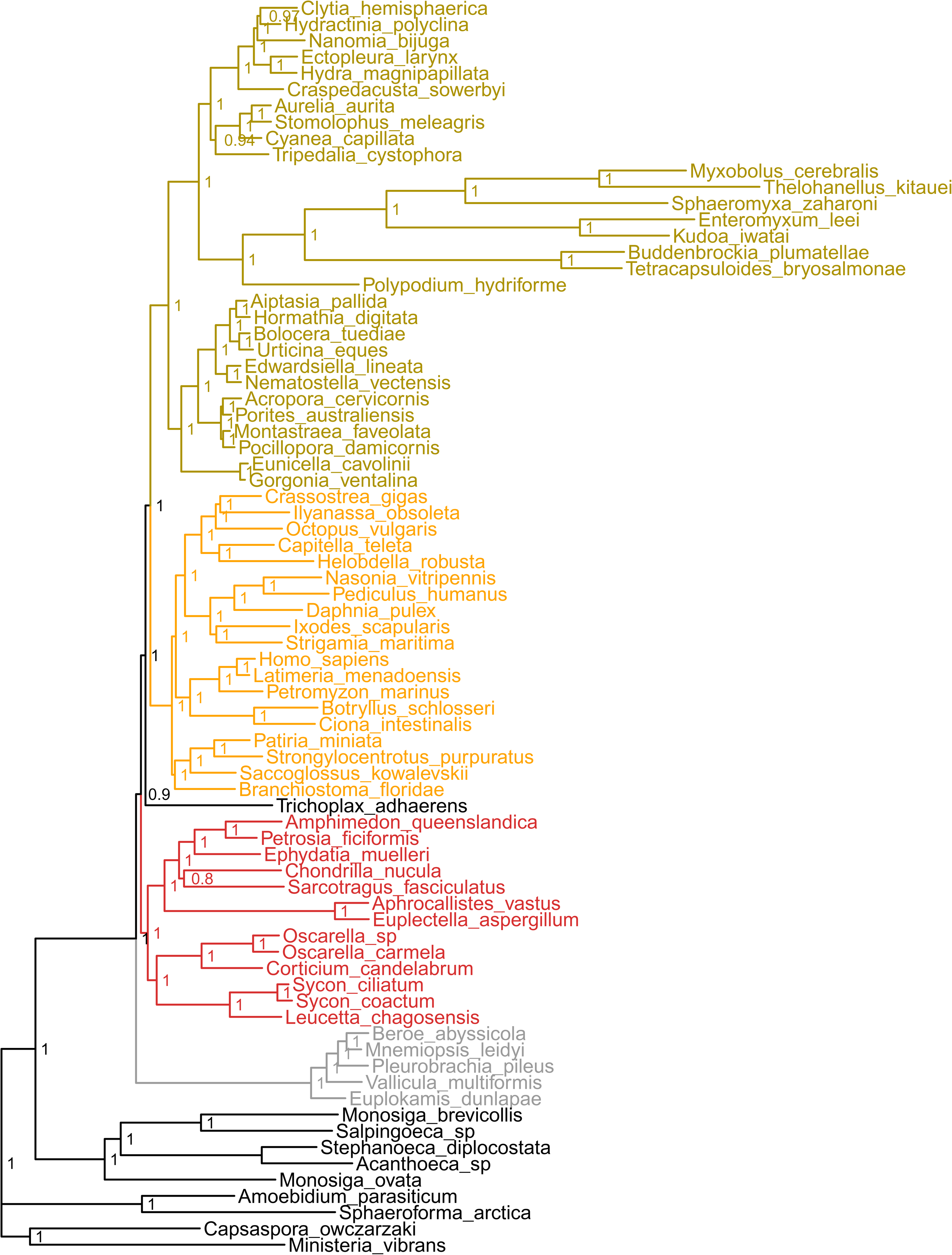
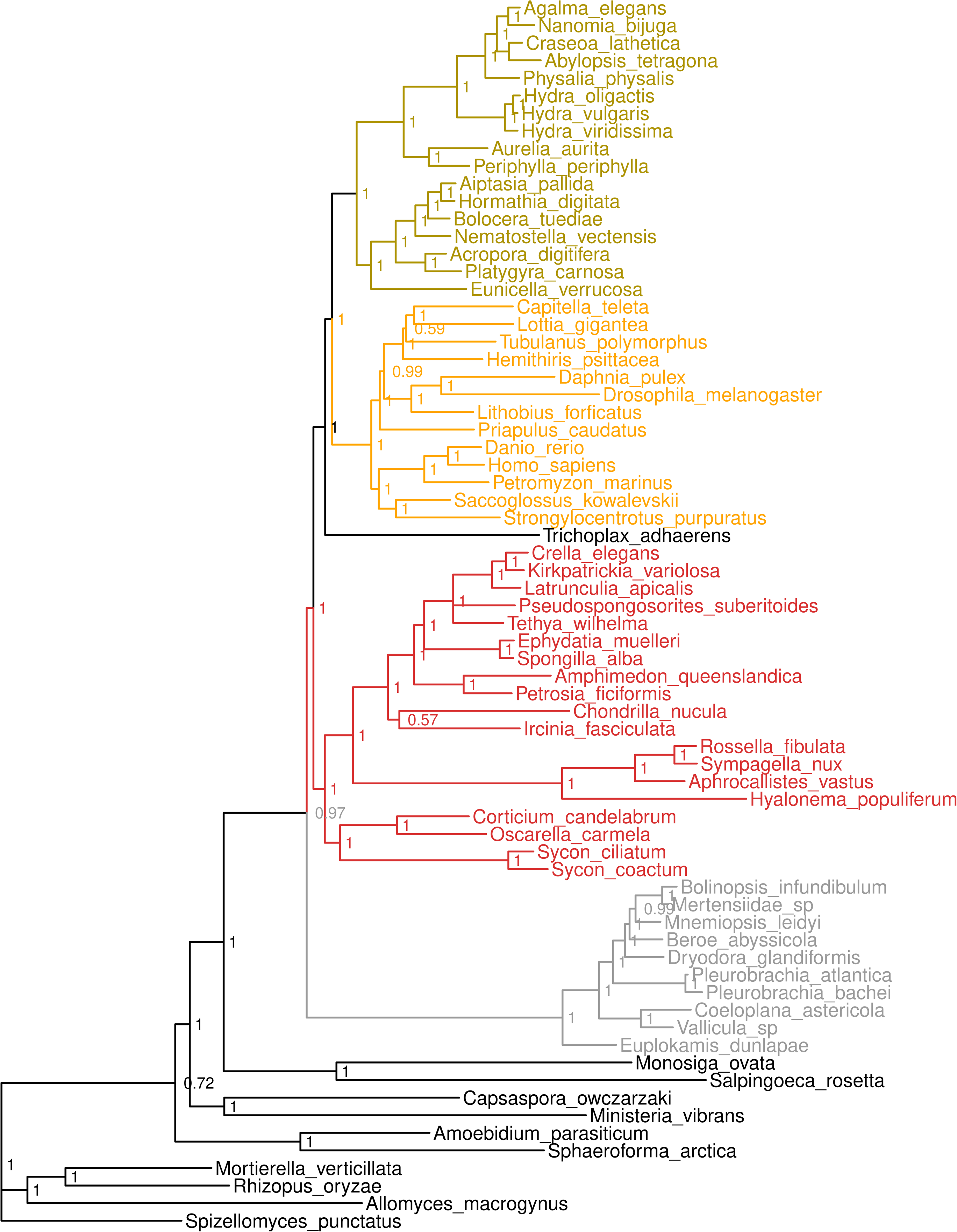
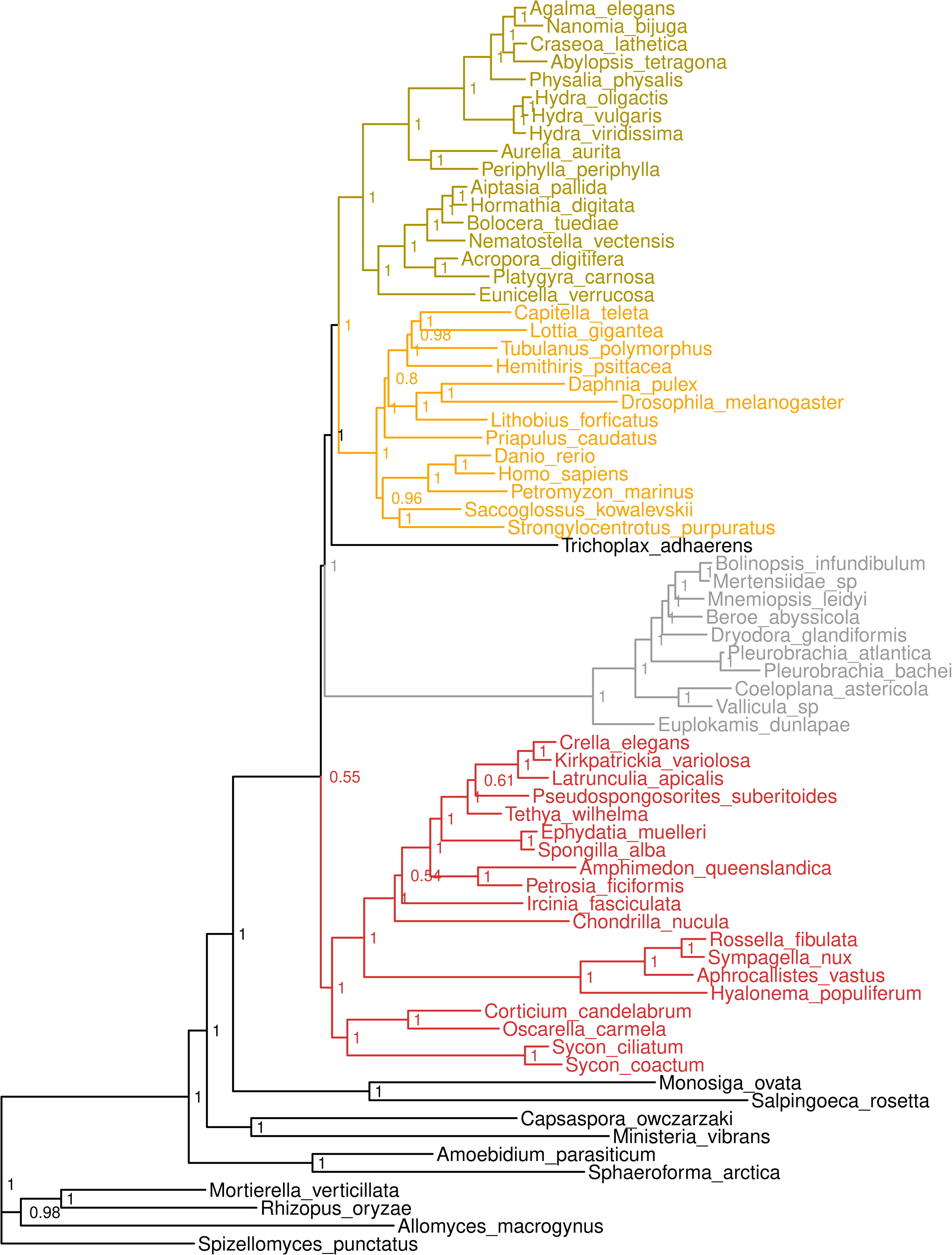
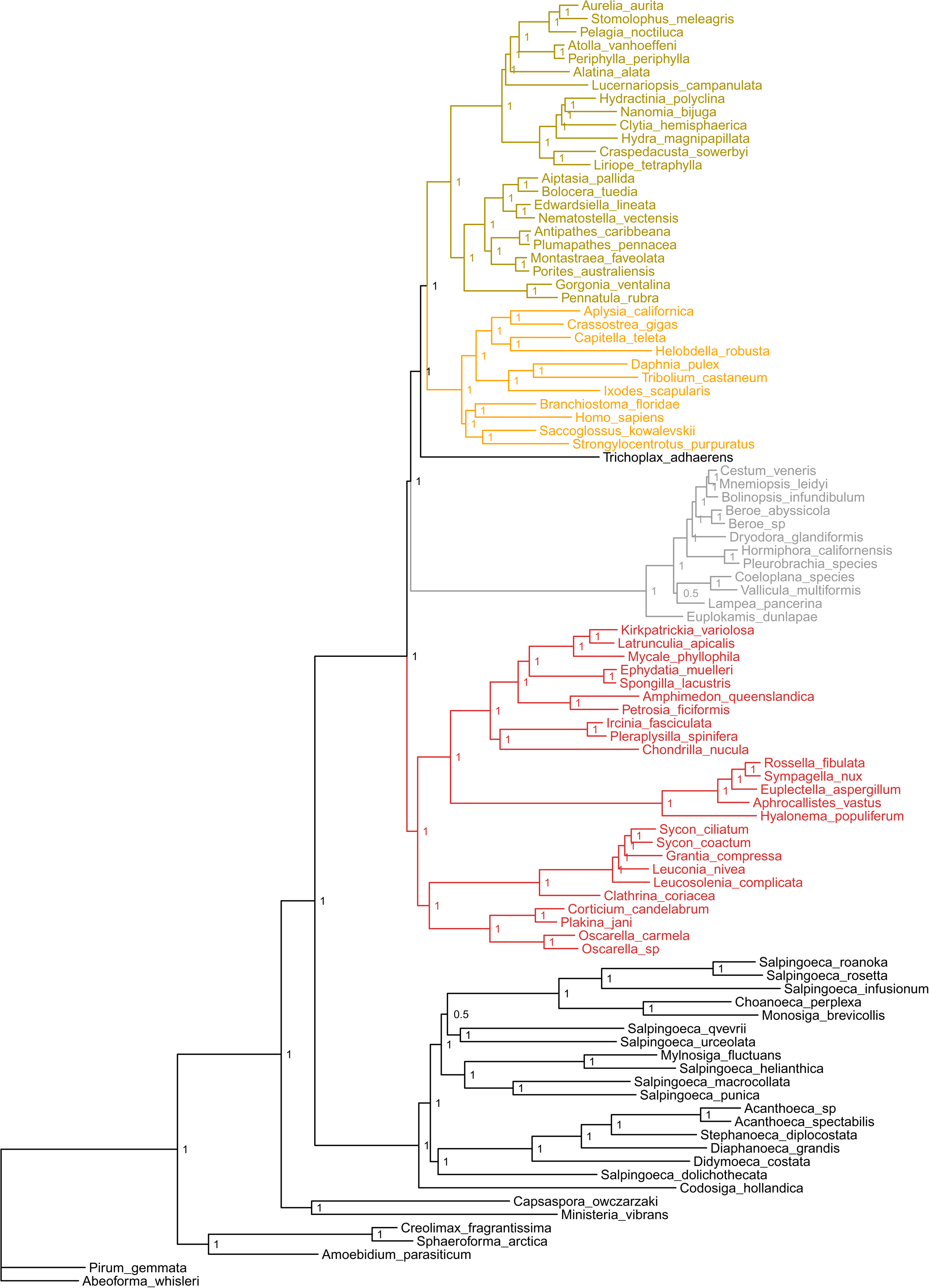
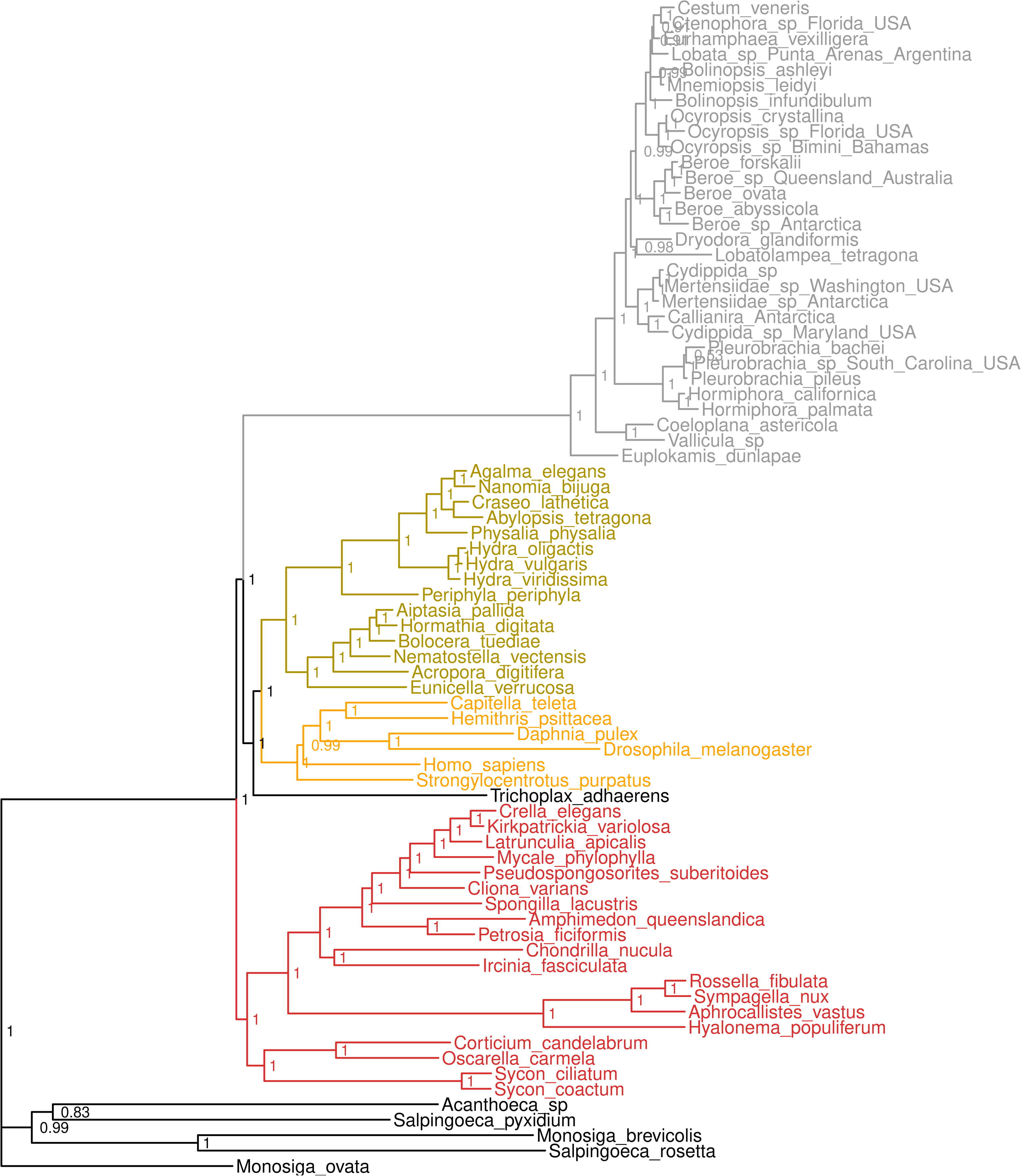
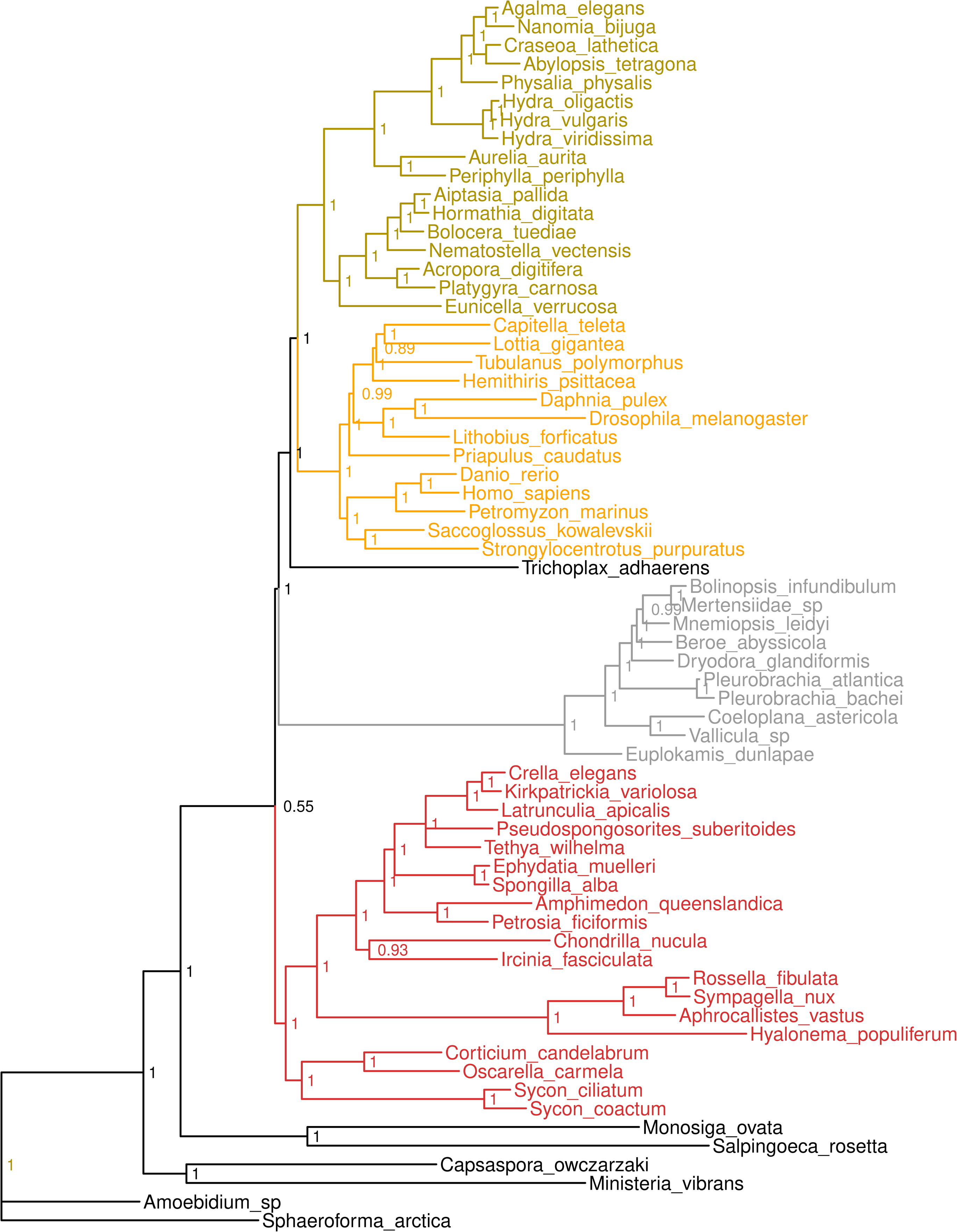
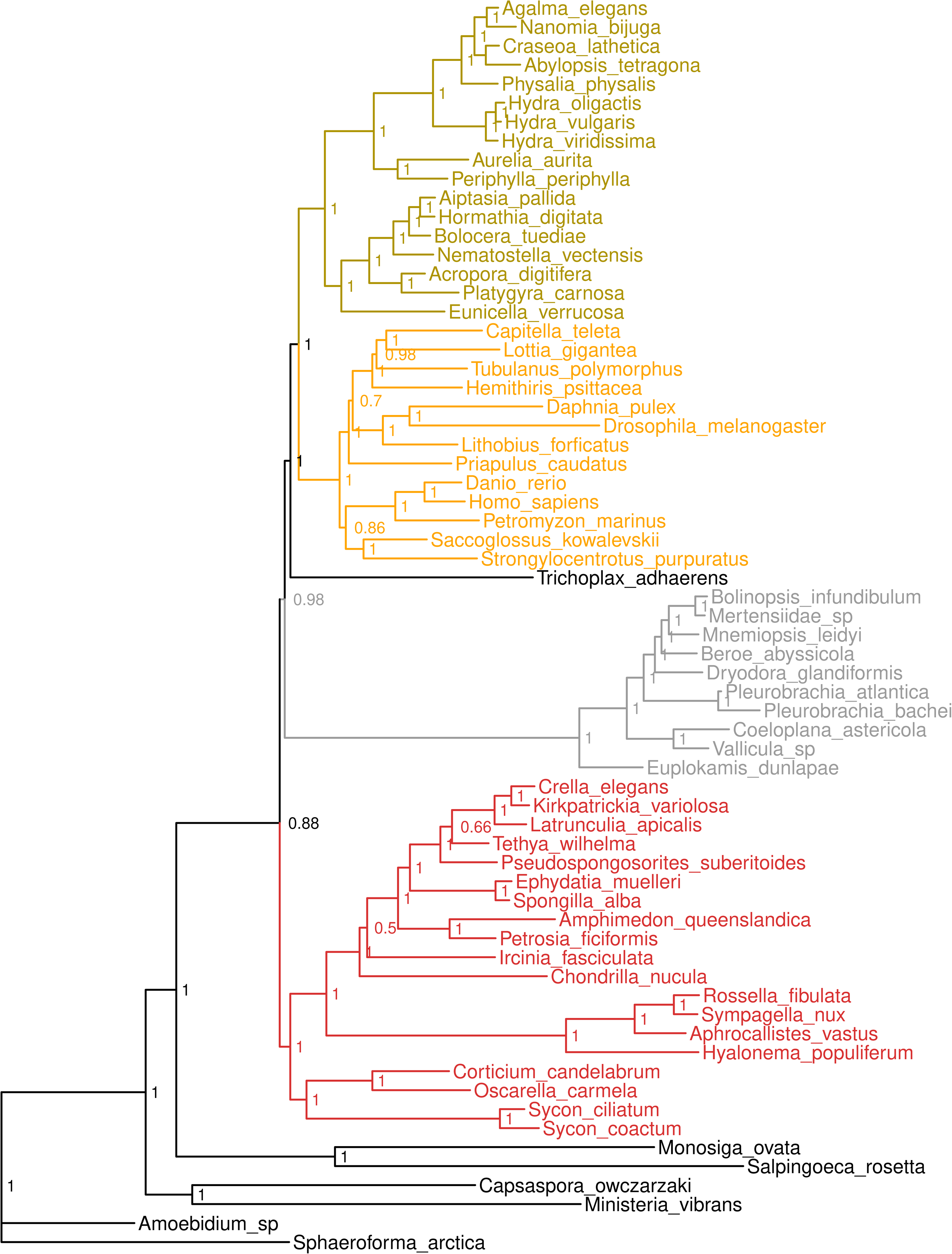
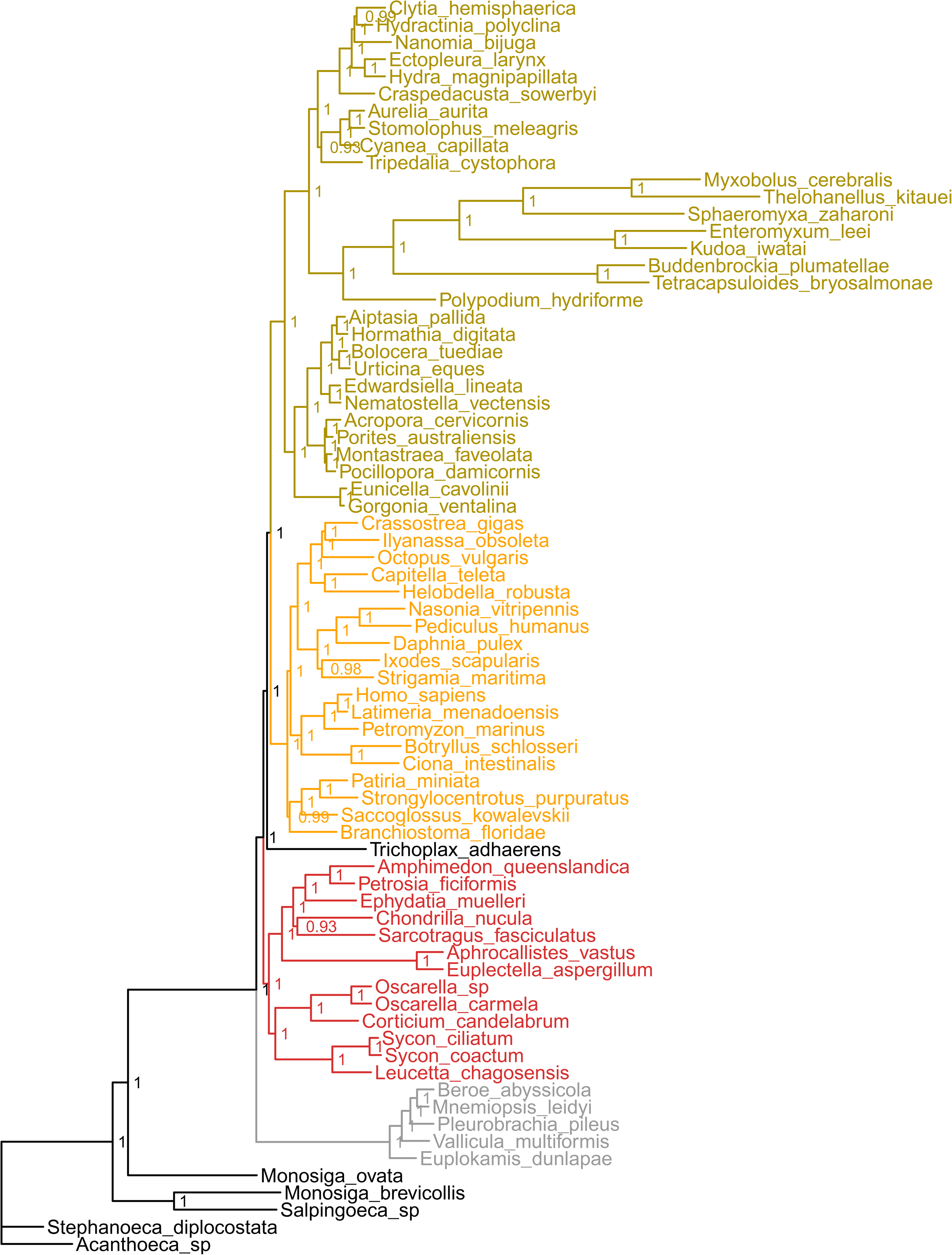
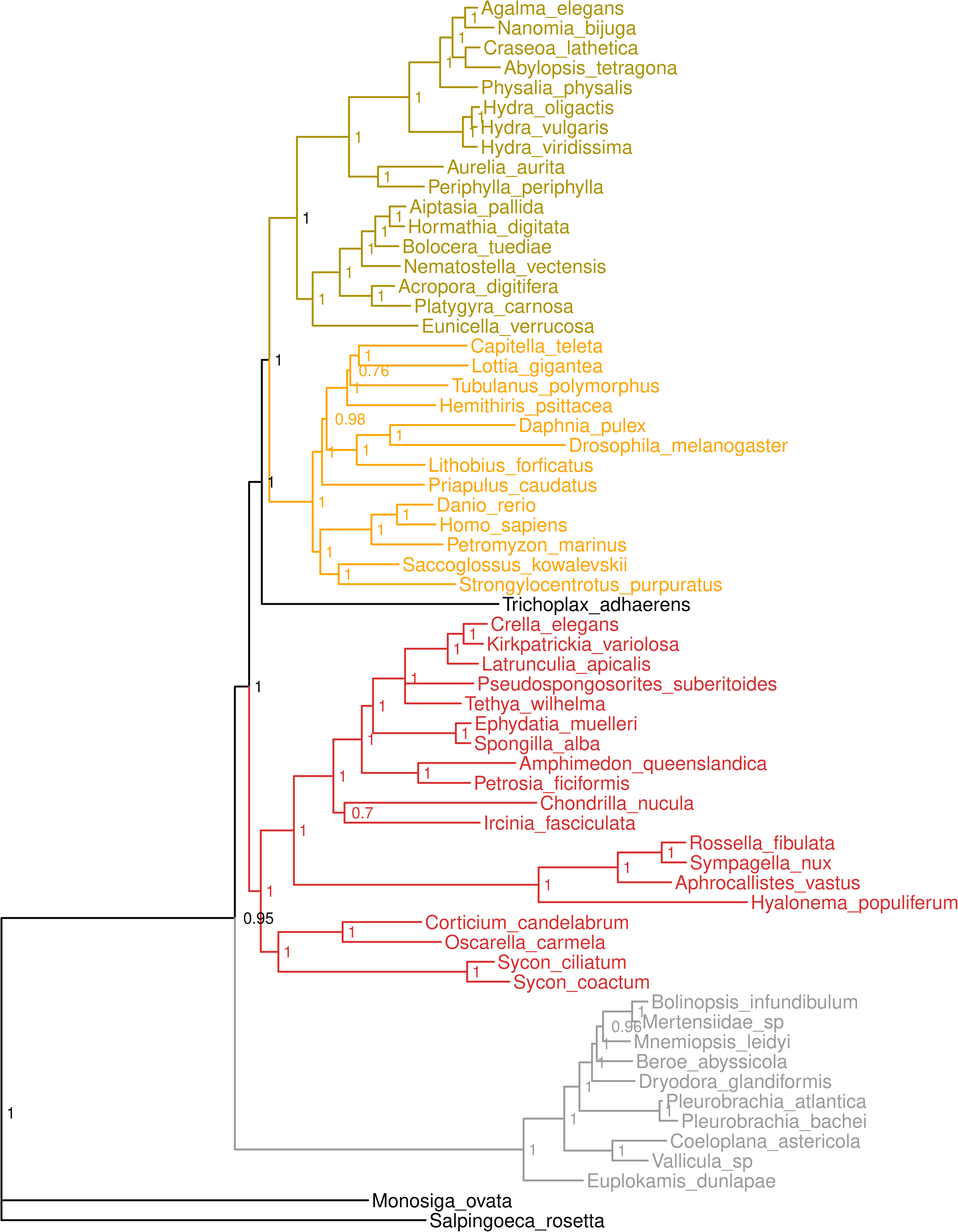
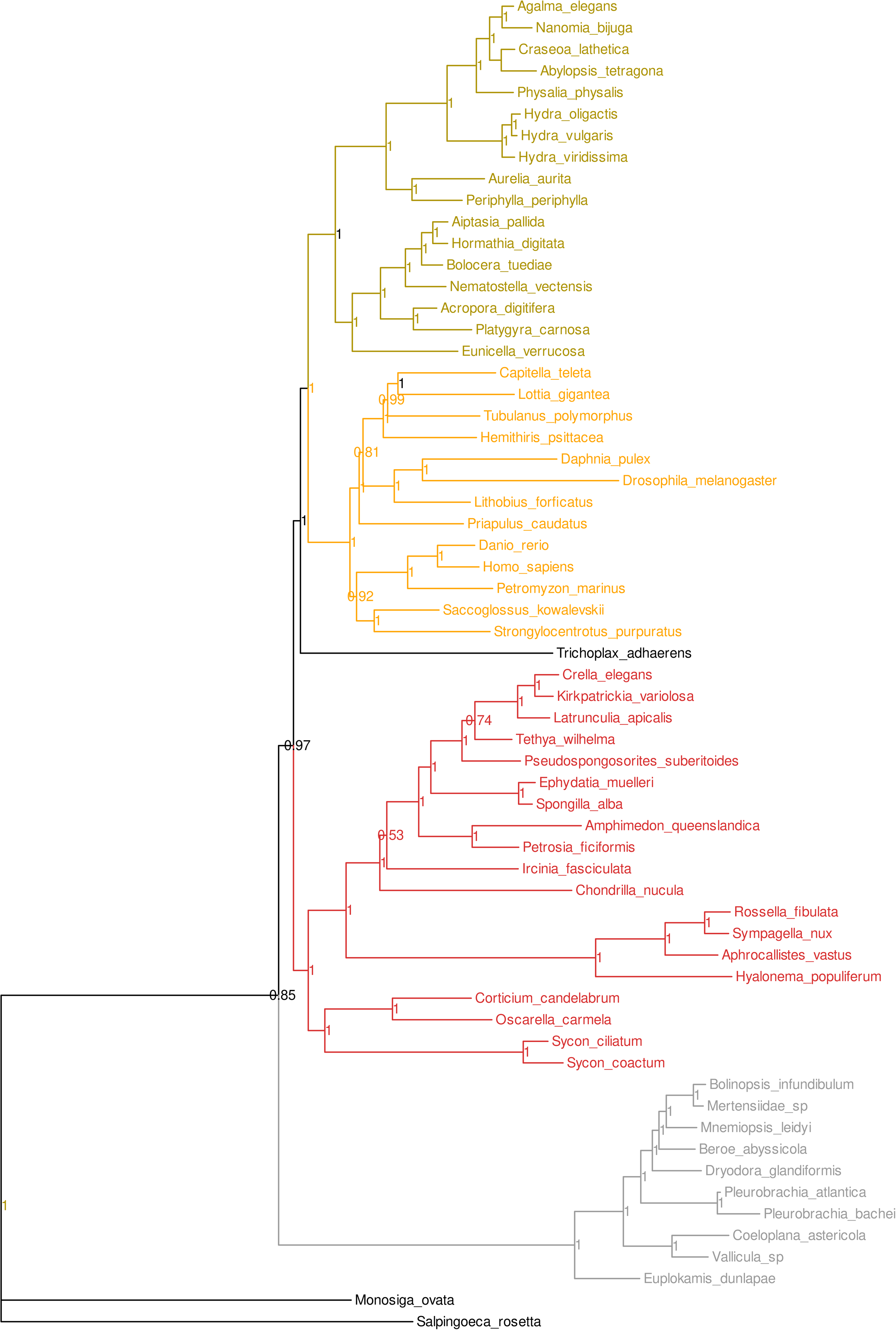
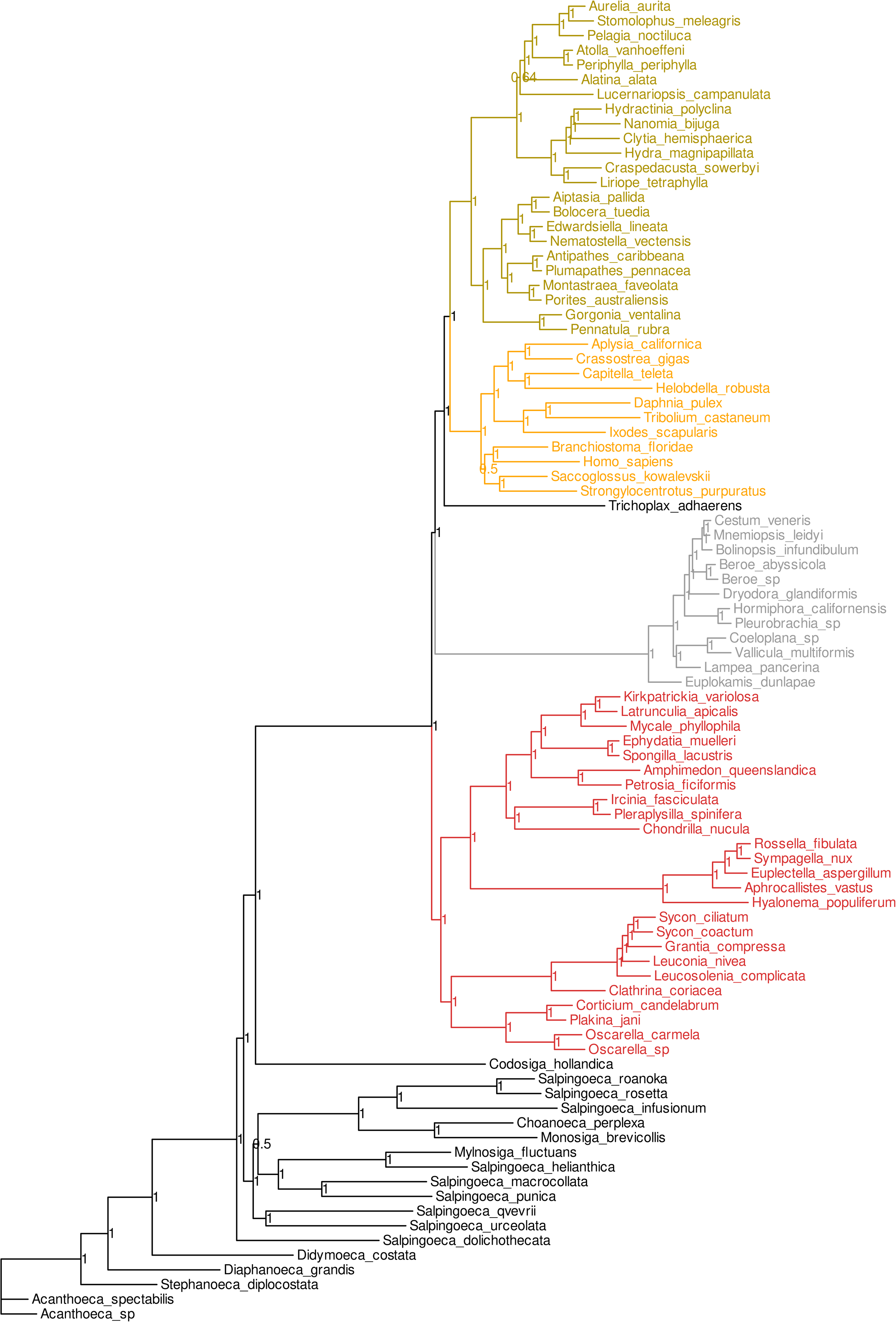
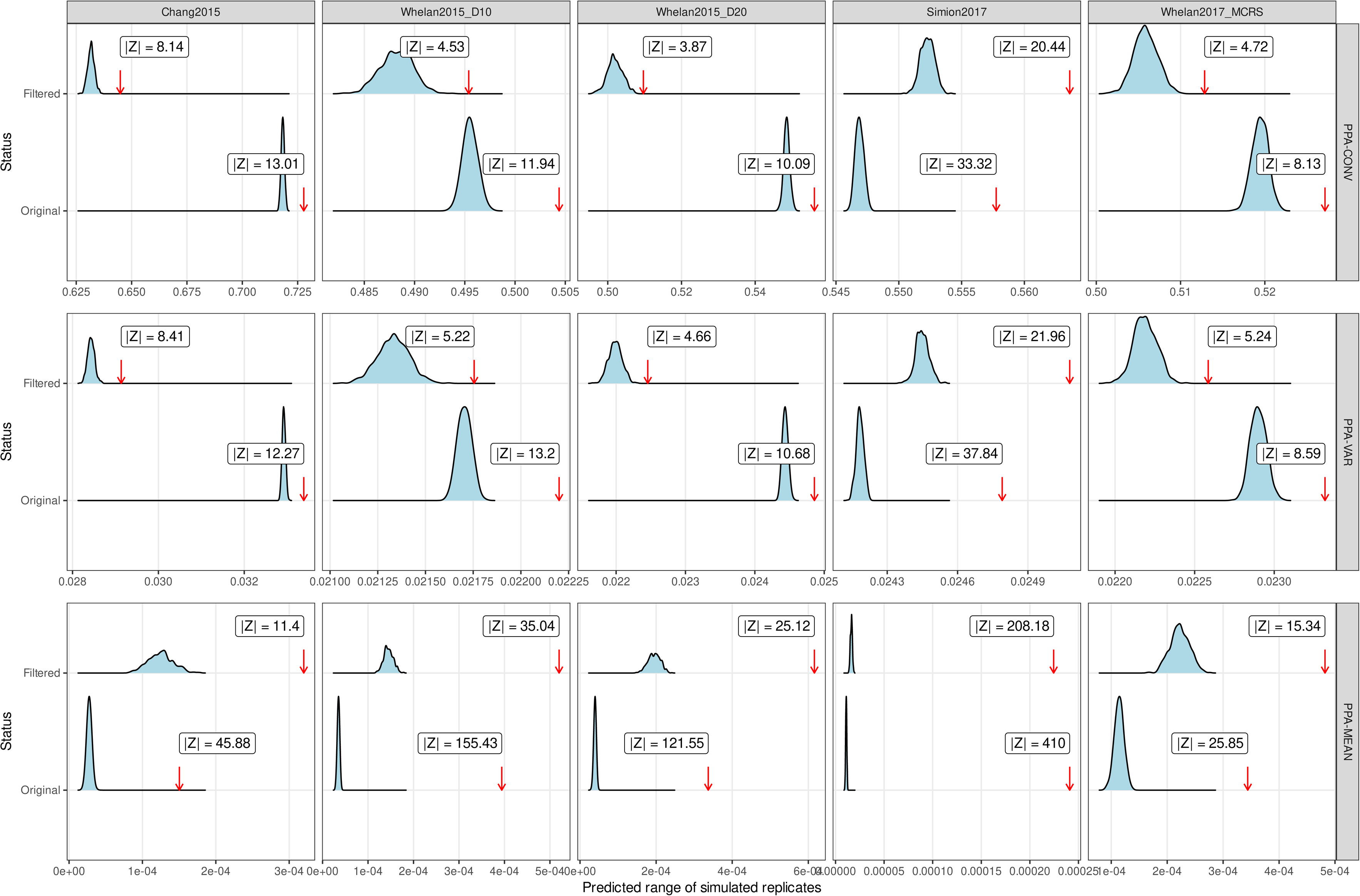

