## Supplementary Information Document for "Improving orthologous signal and model fit in datasets addressing the root of the animal phylogeny"

### **Supplementary information for “Improving orthologous signal and model fit in phylogenomic datasets addressing the root of the animal phylogeny”.**

Charley GP McCarthy^1+^ , Peter O Mulhair^1,2+^, Karen Siu-Ting^3^, Christopher J Creevey^3^ and Mary J O’Connell^1,2^*

^1^Computational and Molecular Evolutionary Biology Research Group, School of Life Sciences, Faculty of Medicine and Health Sciences, University of Nottingham, Nottingham, NG7 2RD, UK

^2^Computational and Molecular Evolutionary Biology Research Group, School of Biology, Faculty of Biological Sciences, University of Leeds, Leeds, LS2 9JT, UK

^3^Institute for Global Food Security, School of Biological Sciences, Queen’s University Belfast, Belfast, BT9 5DL, UK

^+^Both authors contributed equally to this work*.*

*Keywords:* Phylogenomic reconstruction, Animal phylogeny, Orthology, Model fitness, Systematic biases.

### **Supplementary Methodology**

#### **Dataset information**

The five datasets chosen for this analysis varied in the numbers of animal and outgroup taxa sampled (**Table S1**), and per-taxon sampling trends are provided in **Dataset S1.1**. Details on construction and analysis of these datasets in their original studies is provided below.

##### **Chang2015 dataset**

Chang et al. (2015) reconstructed the animal phylogeny to determine in part the phylogenetic position of Myxozoa. The authors assembled a 51,940-site curated dataset consisting of 200 ribosomal and nonribosomal protein sequences sampled from 77 species (Dunn et al., 2008; Philippe et al., 2009; Chang et al., 2015). Using this dataset, the authors recovered a Ctenophora-sister tree under both Bayesian CAT and maximum-likelihood GTR models, and also recovered a Ctenophora-sister tree in a separate analysis which excluded ribosomal sequences (Chang et al., 2015). Data from Chang et al. (2015) was obtained from TreeBASE (https://treebase.org/treebase-web/search/study/summary.html?id=17743) (Piel et al., 2009).

##### **Whelan2015_D10 and Whelan2015_D20 datasets**

Whelan et al. (2015) reconstructed the animal phylogeny to investigate the effect of potential sources of error on the placement of the animal root. Whelan et al. (2015) assembled an initial 81,008-site alignment of 251 sequences from 76 genomes and transcriptomes using HaMStR (Ebersberger et al., 2009; Whelan et al., 2015). A further 24 datasets were then generated using a hierarchical filtering strategy targeting common sources of error (e.g. long-branch attraction and data heterogeneity). All datasets generated by Whelan et al. (2015) recovered a Ctenophora-sister tree under partitioned maximum-likelihood reconstruction, and two of the strictest-filtered datasets also recovered a Ctenophora-sister tree under Bayesian CAT-GTR+G4 reconstruction (Whelan et al., 2015).

The 59,733-site Dataset10, which encompassed 210 sequences from 70 species, provided the main topology presented by the authors (Whelan et al., 2015). Dataset10 was specifically filtered to remove in turn: i) “certain” high-confidence paralogs identified by TreSpEx, ii) heterogeneous genes identified by BaCoCa, and iii) genes and taxa possessing hallmarks of long-branch attraction artefacts (Kück and Struck, 2014; Struck, 2014; Whelan et al., 2015). The 47,632-site Dataset20, which encompassed 178 sequences from the same 70 species, was chosen for its use in other phylogenomic reanalyses of animal tree rooting (Feuda et al., 2017, Li et al., 2021). Dataset20 was constructed in a similar manner to Dataset10, but with the additional removal of “uncertain” paralogs as identified by TreSpEx. Data from Whelan et al. (2015) was obtained from FigShare (http://dx.doi.org/10.6084/m9.figshare.1334306.v3).

##### **Simion2017 dataset**

Simion et al. (2017) reconstructed the animal phylogeny from a dataset built using a semi-automated pipeline intended to reduce known sources of error while improving phylogenetic signal and internal congruence. The authors assembled a 401,632-site dataset composed of 1,719 sequences sampled from 97 species, using a combination of core ortholog detection with OrthoMCL alongside bespoke integration of transcriptomic data and rigorous filtering protocols (Li et al., 2003; Simion et al., 2017). The authors recovered a Porifera-sister tree under maximum-likelihood partitioned reconstruction and Bayesian CAT+G4 reconstruction. The size of the dataset prevented the authors from performing a standard Bayesian reconstruction due to computational limitations - they instead took the approach of performing 100 separate jackknife analyses of approximately 25% of the dataset under CAT+G4 (~100,000 sites per replicate). Data from Simion et al. (2017) was obtained from GitHub (https://github.com/psimion/SuppData_Metazoa_2017).

##### **Whelan2017_MCRS dataset**

Whelan et al. (2017) reconstructed the animal phylogeny as part of an investigation of ctenophoran phylogeny and the ancestral ctenophoran body plan using increased transcriptomic data. The authors assembled two separate sets of hierarchically filtered datasets following the protocols of Whelan et al. (2015), with each set differing in outgroup taxa (Whelan et al., 2015, 2017). For datasets containing only choanoflagellate outgroups, the initial dataset was a 75,840-site alignment taken from 234 sequences across 76 species. As in their previous study, the authors recovered a Ctenophora-sister tree across all datasets using maximum-likelihood partitioned reconstruction and in the strictest-filtered datasets using Bayesian CAT-GTR+G4 reconstruction (Whelan et al., 2015, 2017).

The 49,388-site “Metazoa_Choano_RCFV_strict'' dataset, which encompassed 127 sequences from 76 species, provided the main topology presented by the authors and was chosen for this analysis. Metazoa_Choano_RCFV_strict was specifically filtered to remove i) paralogs, and ii), sequences and taxa which were outliers in terms of relative compositional heterogeneity (Kück and Struck, 2014; Struck, 2014). Five taxa ultimately removed from Metazoa_Choano_RCFV_strict by Whelan et al. (2017) to facilitate convergence in PhyloBayes-MPI were retained for our analysis. Data from Whelan et al. (2017) was obtained from FigShare (http://dx.doi.org/10.6084/m9.figshare.4484138.v1). Partition information for Metazoa_Choano_RCFV_strict was obtained separately (Nathan Whelan, personal communication).

### **Supplementary Results**

#### **Minimal overlap in gene content between animal datasets**

The overlap in gene content between the five animal datasets analyzed in this study was inferred by searching human orthologs where present from all orthogroups for each dataset against the SwissProt human protein set using BLASTp (e = 1e^-4^) (Camacho et al., 2009), and taking the overlap of top hits (Francis and Canfield, 2020; The UniProt Consortium, 2021). According to this approach, little overlap in ortholog content exists between ≥2 or more of these datasets when these results are visualized as an UpSet plot (**Fig. S1**). This is likely due to different approaches in ortholog detection and construction/filtering in each dataset and follows similar comparisons of other animal datasets (Francis and Canfield, 2020). Unsurprisingly, the larger Simion2017 dataset shows the greatest one-to-one overlap with each of the other three datasets and the two Whelan2015 datasets show greatest overlap between each other (**Fig. S1**).

#### **Assessing content differences in original and filtered animal datasets**

**Taxon sampling**

Comparing the distribution of taxa per group among orthogroups that pass our filter versus those that do not shows no impact on relative taxon sampling for Chang2015, Whelan2015_D10 and Whelan2015_D20 (**Fig. S2**). In Simion2017, there is a significant difference in Bilateria sampling (p < 0.05) and in Whelan2017_MCRS a significant increase in ctenophoran taxon sampling in passing OGs coincides with a significant decrease in taxon sampling in other non-bilaterian clans (although the range of taxa present in these clans is more uniform relative to failing orthogroups). The increase in mean ctenophoran taxon sampling among passing orthogroups is likely due to the exclusion of OGs lacking ≥2 ctenophores (**Fig. S2**).

**Data heterogeneity, branch length and relative compositional frequency**

We performed Wilcoxon tests for each dataset between the branch lengths and compositional heterogeneity of orthogroups/orthogroup trees passing or failing our clan_check filter (**Table S2**) (Siu-Ting et al., 2019). For per-sequence compositional heterogeneity, there was no significant difference between orthogroups that passed or failed clan_check filtering. For branch lengths, there was a significant difference between orthogroups from the Simion2017 dataset which passed or failed the clan_check filter (**Table S2**). No individual orthogroups displayed evidence of per-family compositional heterogeneity as assessed by the χ^2^ test for compositional homogeneity in p4 (p = 1), but each data matrix failed the test indicating dataset-wide heterogeneity (p = 0) (Foster, 2004). We also calculated RCFV values for each dataset using BaCoCa, to infer the compositional biases of orthogroups passing or failing our clan_check filter (Zhong et al., 2011; Kück and Struck, 2014) (**Fig. S3**). It appears that clan_check filtering may remove many RCFV outliers and produce datasets more uniform with respect to this kind of data heterogeneity. The RCFV values observed in Whelan2017_MCRS are much lower and tightly-distributed than every other dataset, due to that dataset being specifically constructed in part to exclude RCFV outliers (Whelan et al., 2017) (**Fig. S3**).

**Gene ontology**

For each dataset, human sequences were extracted (where available) from orthogroups passing or failing our clan_check filter. Predictive sequence annotation was carried out on all human sequences using InterProScan, and we performed Pearson’s χ^2^ tests for independence for all three gene ontology (GO) categories in passing or failing orthogroups (p < 0.05) (**Table S3**) (Jones et al., 2014; Siu-Ting et al., 2019). In four out of five datasets, no significant difference was observed in GOs between passing or failing orthogroups (**Table S3**). This implies that in most cases our clan_check filtering approach is not biasing datasets with regarding to their predicted functional composition. As for Chang2015, the significant difference in GOs between passing and failing orthogroups may reflect the composition of that dataset as a curated combination of ribosomal and non-ribosomal protein sequences (Chang et al., 2015).

#### **Assessment of PhyloBayes-MPI chain and tree convergence**

Trace plots for each PhyloBayes-MPI run suggest that the parameters of both chains converged relatively early in each run (i.e., within 1,000 iterations) (**Figs. S9-S13**). Quantitative assessment of convergence indicated that each run had generated at least 100 independent points (effsize) per parameter, apart from mean site entropy for Simion2017_filtered (71 points). Discrepancies between chain parameters (rel_diff) were under the acceptable cutoff of 0.3 for every parameter except tree length in Chang2015_filtered and three parameters in Simion2017_filtered (Lartillot, 2020). The presence of a long *Polypodium+*Myxozoa branch within Cnidaria is the likely cause for discrepancies in tree length in the Chang2015_filtered dataset (Chang et al., 2015). As for Simion2017_filtered the higher discrepancies observed for log-likelihood, mean site entropy and the alpha parameter are likely a reflection of dataset size. Assessing changes in tree topology per iteration in each chain using Robinson-Foulds distances indicates that only minor changes occur after 1,000 iterations for each dataset (**Fig. S14**).

Each dataset except Simion2017_filtered also achieved acceptable convergence (maxdiff < 0.3) in tree space when sampled using bpcomp (burn-in = 5,000). For Simion2017_filtered, maxdiff remained at 1 even after reaching 7,500 iterations. Inspecting the bipartitions observed in each chain via bpcomp indicates that this lack of convergence is being driven solely by large discrepancies between the chains in the placements of *Lampea pancerina* within Ctenophora and a pair of choanoflagellates (*Salpingoeca qvevrii* and *Salpingoeca urceolata*) within the outgroup. In each instance, both chains almost completely diverge in topology at these nodes (maxdiff = 1) but display no discrepancies in every other bipartition (maxdiff < 0.1). This is reflected in the lower posterior support (PP = 0.5) at these nodes compared to all other nodes in the Simion2017_filtered tree (PP = 1) (**Fig. S18**). The RF distance plots for each chain in this run show almost no change in tree topology after ~1,000 iterations, indicating that the failure of this dataset to converge in tree space may be down to discrepancies in intraphylum relationships which become fixed early in each chain and not necessarily a failure to resolve deeper relationships (**Fig. S14**).

#### **Comparison of original and clan_check filtered animal phylogenies**

##### **Chang2015_filtered dataset**

We recovered a Ctenophora-sister tree under CAT-GTR+G4, in line with the original findings of Chang et al. (2015) (**Fig. S15**). Our tree is almost identical to the original Chang2015 tree in both topology and statistical support, with the exception of the placement of the model Cephalochordata species *Branchiostoma floridae*. In the original tree *B. floridae* branches sister to the tunicates and craniates within Chordata, but in our reconstruction *B. floridae* is placed sister to a clade containing the echinoderms and hemichordates (Ambulacraria) and implies a paraphyletic Deuterostomia. Recent studies have suggested deuterostome monophyly may be artefactual and definitive resolution of major deuterostome relationships has proven difficult (Cannon et al., 2016; Philippe et al., 2019; Kapli and Telford, 2020). Given the monophyly of the chordates has been established in the literature, the placement of *B. floridae* outside Chordata has more likely arisen in this case from informational loss (Bourlat et al., 2006).

##### **Whelan2015_D10_filtered dataset**

We recover a Ctenophora-sister tree using a filtered version of Dataset 10 under Bayesian CAT-GTR+G4, as Whelan et al. (2015) did with the original dataset in their study (**Fig. S16**). Our filtered tree is almost identical in topology to the original tree, with the only major difference being an inability to resolve the relationship between *Pseudospongosorites suberitodies* and *Tethya wilhelma* within the Demospongaie. As the node separating these two sponges had relatively weak support in the original tree (61% BS), it is likely that there is a lack of sufficient signal to resolve this node within the filtered dataset (Whelan et al., 2015).

**Whelan2015_D20_filtered dataset**

We recovered a Porifera-sister tree using a filtered version of Dataset 20 under Bayesian CAT-GTR+G4 (**Fig. S17**), which contrasts with the Ctenophora-sister tree recovered by Whelan et al. (2015) in their original analysis. Notably this dataset includes quite distant outgroups leading back to Fungi, a sampling strategy which has previously been thought to produce predominantly Ctenophora-sister trees. The noticeably low support (P = 0.55) at the node separating Porifera and Ctenophora suggests that there is still conflicting signal regarding the relationship between these two groups within this dataset after filtering (**Fig. S17**). A number of sponge taxa differ slightly in the position within Porifera in the filtered tree relative to the original maximum-likelihood tree, such as the relationship of *Ircinia fasciculata* and *Chondrilla nucula* or relationships within the aforementioned Demospongaie (Whelan et al., 2015).

##### **Simion2017_filtered dataset**

We recovered a Porifera-sister tree using a filtered version of the Simion2017 dataset under Bayesian CAT-GTR+G4, similar to the original reconstruction of the full dataset jack-knifed CAT+G4 approach by Simion et al. (2017) (**Fig. S18**). Alongside supplementary Bayesian analysis in Simion et al. (2017), our analysis suggests that recovery of a Porifera-sister tree may not be solely limited to datasets with choanoflagellate-only outgroups (Simion et al., 2017b). In terms of topological differences between the original tree and our filtered tree, the only noticeable difference is the placement of *Lucernariopsis campanulata* within the Medusazoa. In the original tree *L. campanulata*, the sole taxon representing Staurozoa (stalked jellies), branched as sister to Hydrozoa with relatively weak support (71% JS) (Simion et al., 2017b). In contrast, our filtered dataset tree shows *L. campanulata* branching sister to a clade encompassing both Cubozoa (box jellies) and Scyphozoa (true jellies) (**Fig. S18**). As with the lack of resolution for deuterostomes in Chang2015_filtered it is inappropriate to assume an animal root dataset would provide definitive information on internal cnidarian relationships, but it is worth noting that this topology is similar to those recovered in recent phylogenomics studies of Cnidaria (Zapata et al., 2015; Kayal et al., 2018).

##### **Whelan2017_MCRS_filtered dataset**

Whelan et al. (2017) recovered a Ctenophore-sister tree using their “Metazoa_Choano_RCFV_strict” dataset in the original study. In our filtered 22,280-site version of the same dataset we instead recover a Porifera-sister tree under CAT-GTR+G4 (**Fig. S19**). This is in line with previous observations that limiting outgroup sampling to the choanoflagellates may produce a Porifera-sister tree under CAT-GTR+G4, which Whelan et al. (2017) contradicted (Pisani et al., 2015; Halanych et al., 2016; Whelan et al., 2017; Li et al., 2021). Rooting aside, comparing the topology of our filtered tree with that of the original “Metazoa_Choano_RCFV_strict” tree reveals two notable differences within Ctenophora (**Fig. S19**) In the original tree, the Lobata-like species *Lobatolampea tetragona* is placed as sister to a Lobata clade with very weak support (46% BS), whereas in our reconstruction *L. tetragona* branches sister to the Cydippia-like species *Dryodora glandiformis* (Whelan et al., 2017). Similarly, within the Lobata clade the placement of *Bolinopsis infundibulum* relative to *Bolinopsis ashleyi* cannot be resolved in our reconstruction but is resolved in the original tree albeit with weak support (52% BS) (Whelan et al., 2017). There is likely some informational loss amongst internal relationships after dataset filtering which may explain these variations within this group. Similarly *Strongylocentrotus purpuratus* and *Homo sapiens* do not form a monophyletic Deuterostomia clade in our filtered tree, which is likely also a consequence of informational loss (**Fig. S19**).

#### **Effect of outgroup exclusion on rooting in animal datasets**

**Holozoan and Choanoflagellate outgroup trees**

PhyloBayes-MPI analysis was repeated on both Whelan2015_D10_filtered and Whelan2015_D20_filtered with fungal outgroups removed (indicated as _FilteredHolo).These new phylogenies – and specifically which animal root hypothesis they supported - were compared with those of the Chang2015_filtered and Simion2017_filtered dataset which both contained Holozoan and Choanoflagellate outgroups. As is discussed in the main text, three of the four Holozoa+Choanoflagellata outgroup trees recovered a Porifera-sister tree whereas Chang2015_filtered recovers a Ctenophora-sister tree. Outside of the change in root hypothesis support when fungal outgroups are removed, there are minor differences between the two sets of trees from Whelan et al. (2015) when fungal outgroups are present or removed - largely in slight increases or decreases in support values along internal branches (**Figs. S20-21**).

**Choanoflagellate-only outgroup trees**

PhyloBayes-MPI analysis was repeated on all filtered datasets except Whelan2017_MCRS_filtered with all non-Choanoflagellate ougroups removed (indicated as FilteredChoano). The effect of differing outgroup inclusion on the animal root hypothesis supported by each dataset was discussed in the main text. In terms of impact elsewhere in the trees, there appeared to be only minor differences in support values along internal branches in trees with non-Choanoflagellate outgroups excluded compared to all other trees generated in this study (**Figs. S22-25**).

#### **Additional posterior predictive analysis of model fit**

For the remaining two site-heterogeneity statistics (PPA-CONV and PPA-VAR) and the remaining lineage-heterogeneity statistic (PPA-MEAN) not discussed in the main text, |Z| > 5 across each statistic for each filtered dataset except PPA-CONV in Whelan2015_D10_filtered, Whelan2015_D20_filtered and Whelan2017_MCRS_filtered as well as PPA-VAR in Whelan2015_D20_filtered (**Fig. S26**). These results indicate that substantial amounts of site-specific and particularly lineage-specific heterogeneity exist in each filtered dataset. As before, the model fit of these filtered datasets represent substantial improvements over the original datasets (**Fig. S26**).

### **Supplementary Tables**

| **Dataset** | **Outgroup** | **Porifera** | **Ctenophora** | **Placozoa** | **Cnidaria** | **Bilateria** |
| --- | --- | --- | --- | --- | --- | --- |
| **Chang2015** | 9 | 13 | 5 | 1 | 30 | 19 |
| **Whelan2015D10 and Whelan2015D20** | 10 | 19 | 10 | 1 | 17 | 13 |
| **Simion2017** | 25 | 25 | 12 | 1 | 23 | 11 |
| **Whelan2017MCRS** | 5 | 19 | 30 | 1 | 15 | 6 |

**Table S1**. **Taxon sampling per clan across the four animal datasets chosen for this study.** Outgroup clan composition (i.e. presence of Fungi, Holozoans and Choanoflagellates) varied between datasets.

| **Dataset** | **Branch length (Pass / Fail)** | **CH (Pass / fail)** |
| --- | --- | --- |
| **Chang2015** | **0.4** | **0.74** |
| **Whelan2015D10** | **1** | **0.459** |
| **Whelan2015D20** | **0.28** | **0.16** |
| **Simion2017** | **3.084527e-09** | **0.6** |
| **Whelan2017MCRS** | **0.83** | **0.137** |

**Table S2.** Wilcoxson test results for differences in branch lengths and compositional heterogeneity in animal datasets (p < 0.05). Comparison made between orthogroups passing or failing the ≥3 clans filter.

| **Dataset** | **Genes passing clan_check filter** | | | **Genes failing clan_check filter** | | | **Χ^2^ test results** |
| --- | --- | --- | --- | --- | --- | --- | --- |
|  | **Cellular component** | **Molecular function** | **Biological process** | **Cellular component** | **Molecular function** | **Biological process** |  |
| **Chang2015** | **12** | **101** | **45** | **83** | **176** | **113** | **Χ^2^ = 19.389,**  **df = 2, p = 6.161e-05** |
| **Whelan2015D10** | **5** | **35** | **22** | **34** | **150** | **96** | **Χ^2^ = 0.83934, df = 2, p = 0.6573** |
| **Whelan2015D20** | **8** | **14** | **16** | **26** | **114** | **75** | **Χ^2^ = 4.0684, df = 2, p = 0.13078** |
| **Simion2017** | **91** | **351** | **174** | **177** | **736** | **377** | **Χ^2^ = 0.46637, df = 2, p = 0.792** |
| **Whelan2017MCRS** | **9** | **54** | **34** | **31** | **84** | **48** | **Χ^2^ = 4.5514, df = 2, p = 0.1027** |

**Table S3**. Pearson’s χ^2^ tests for independence of gene ontology categories in animal datasets (p < 0.05). Gene ontologies assigned using InterProScan, and binned into namespaces using GOATools. χ^2^ = chi-squared statistic, df = degrees of freedom.

### **Supplementary Figures**

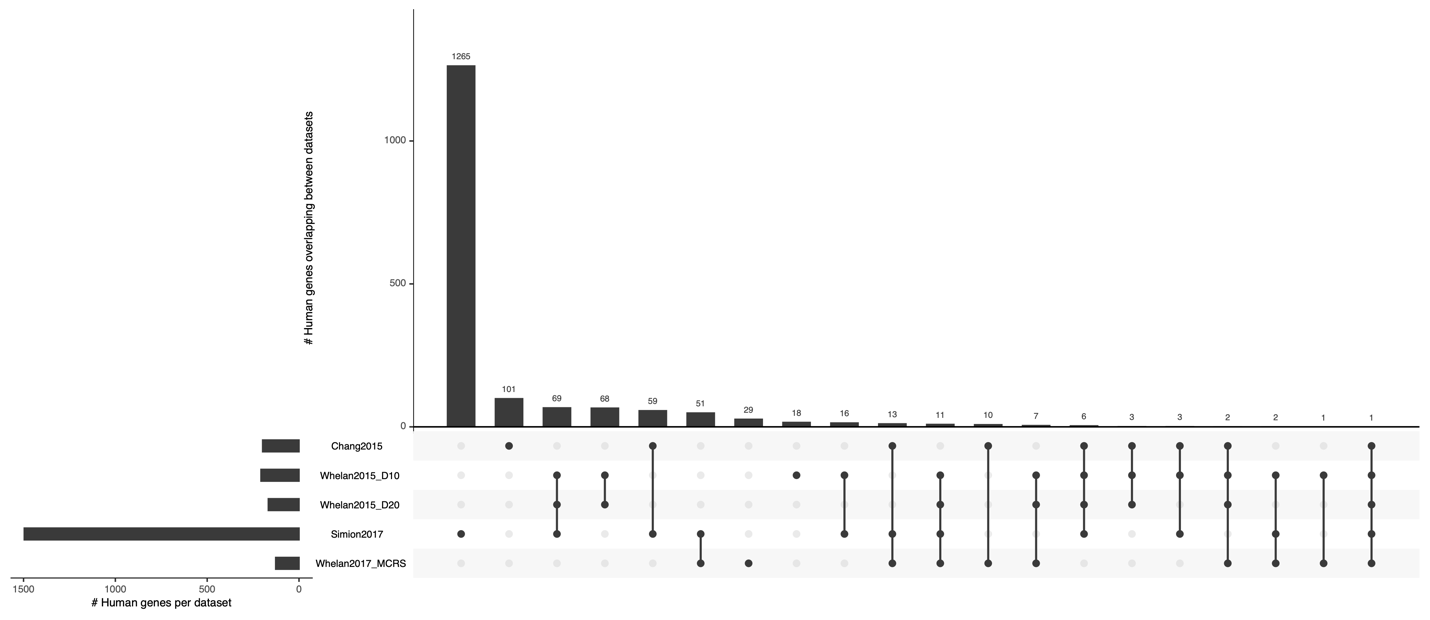

**Figure S1**. **Overlap in human sequence content inferred between animal datasets.** Overlap determined following Francis & Canfield (2020): human sequences were extracted from orthogroups where present in each dataset and searched against the Swiss-Prot human protein dataset using BLASTp (e = 1e-4). The intersections between datasets in this plot represents the overlap in top hits from BLASTp output which are visualized as an UpSet plot: the rows at the bottom represent possible singleton or intersecting sets of overlapping top hits and the bar chart above displays the number of top hits in each set.

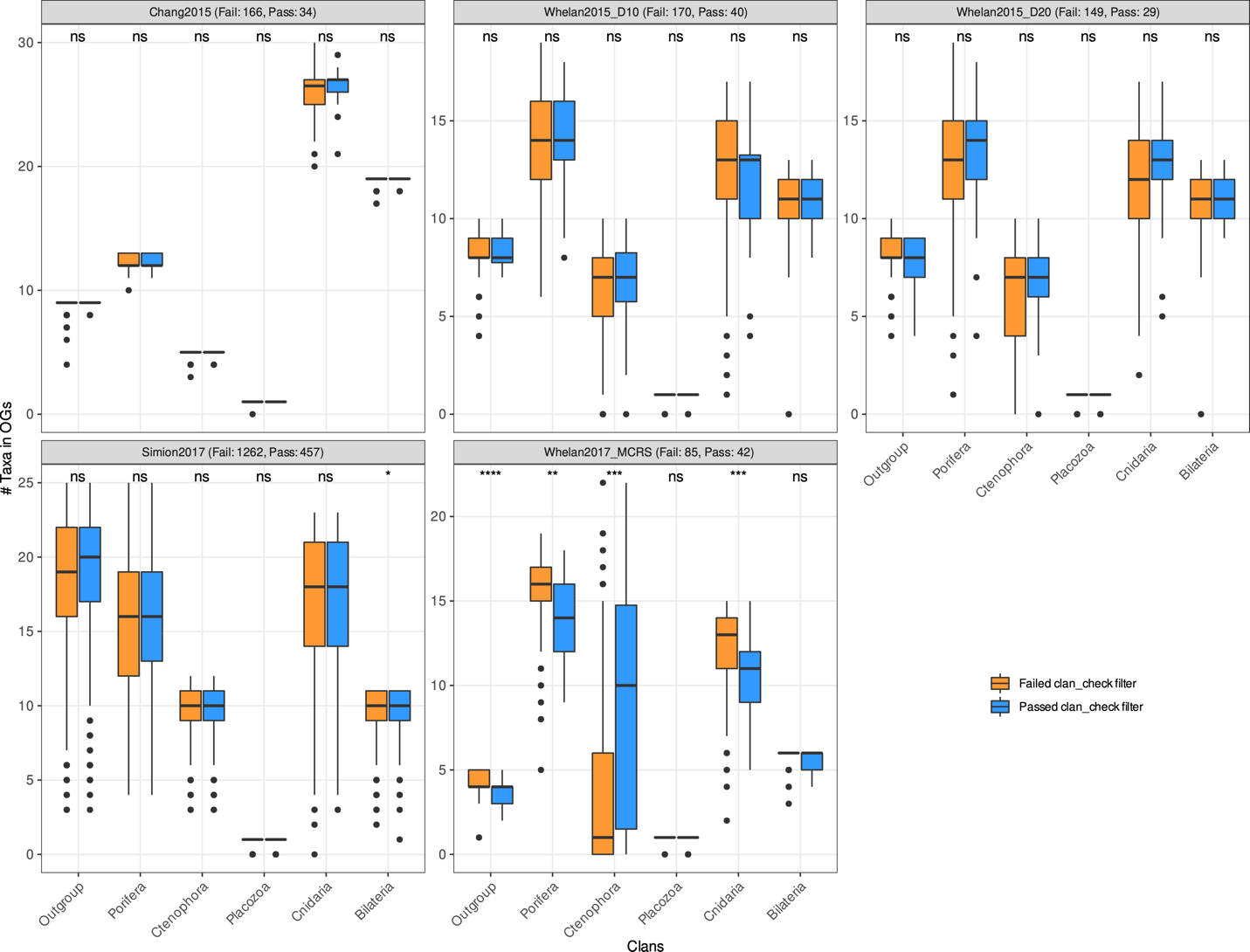

**Figure S2**. **Taxon sampling per clan among orthogroups which failed or passed our clan_check filter.** Boxplot represents the distribution per clan of a member taxa which passed (blue) or failed (orange) the ≥3 clans filter, for each animal dataset. Numbers of passing and failing orthogroups per dataset indicated in the text above each panel. Wilcoxson test results: **** = p ≤ 0.0001, *** = p ≤ 0.001, ** = p ≤ 0.01, * = p ≤ 0.05, ns = not significant (p > 0.05).

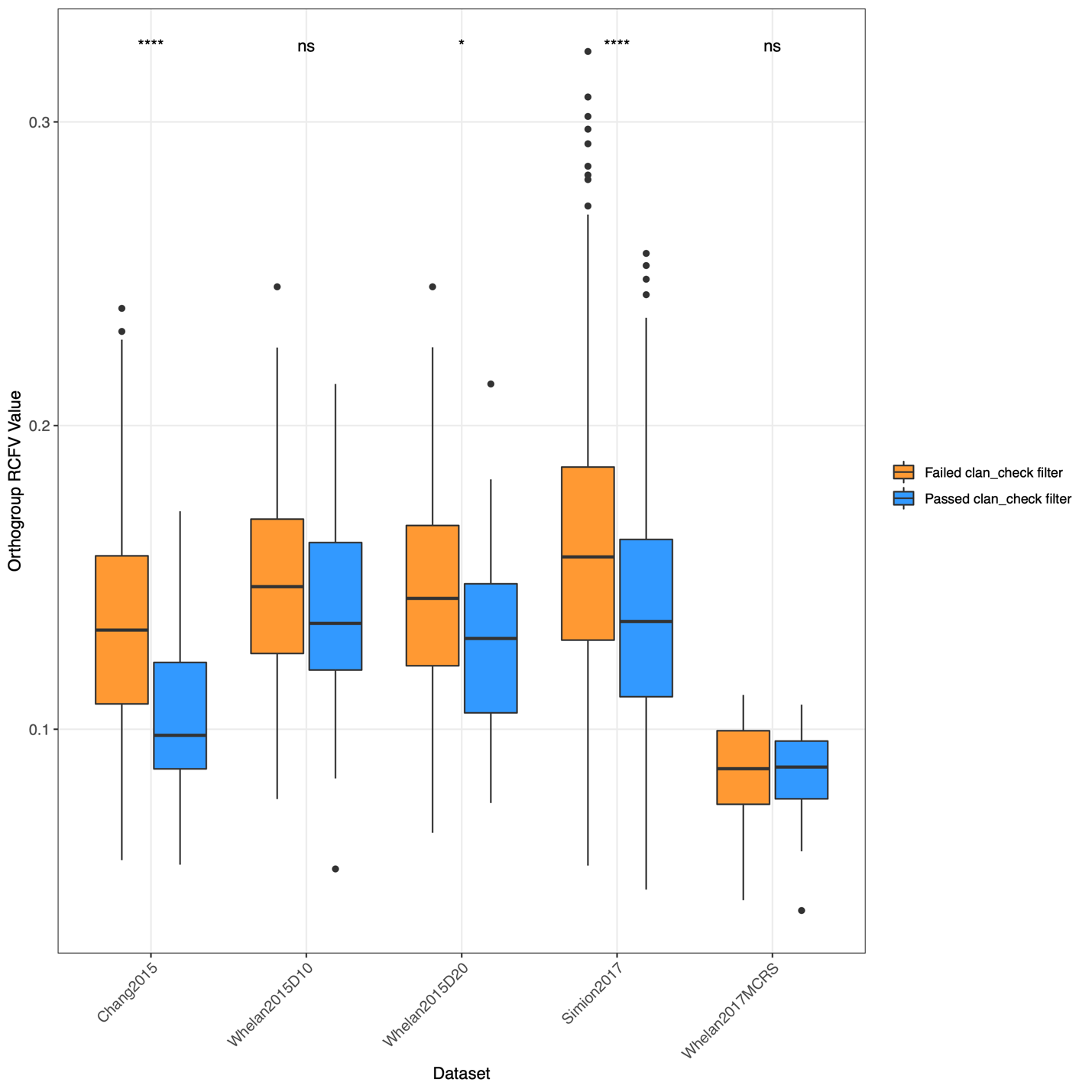

**Figure S3.** **Comparison of RCFV values between orthogroups passing or failing our clan_check filter in five animal datasets.** RCFV values for all orthogroups in each dataset calculated using BaCoCa. Boxplots represent the distribution of RCFV values across orthogroups which passed (blue) or failed (orange) the ≥3 clans filter. Wilcoxson test results: **** = p ≤ 0.0001, *** = p ≤ 0.001, ** = p ≤ 0.01, * = p ≤ 0.05, ns = not significant (p > 0.05).

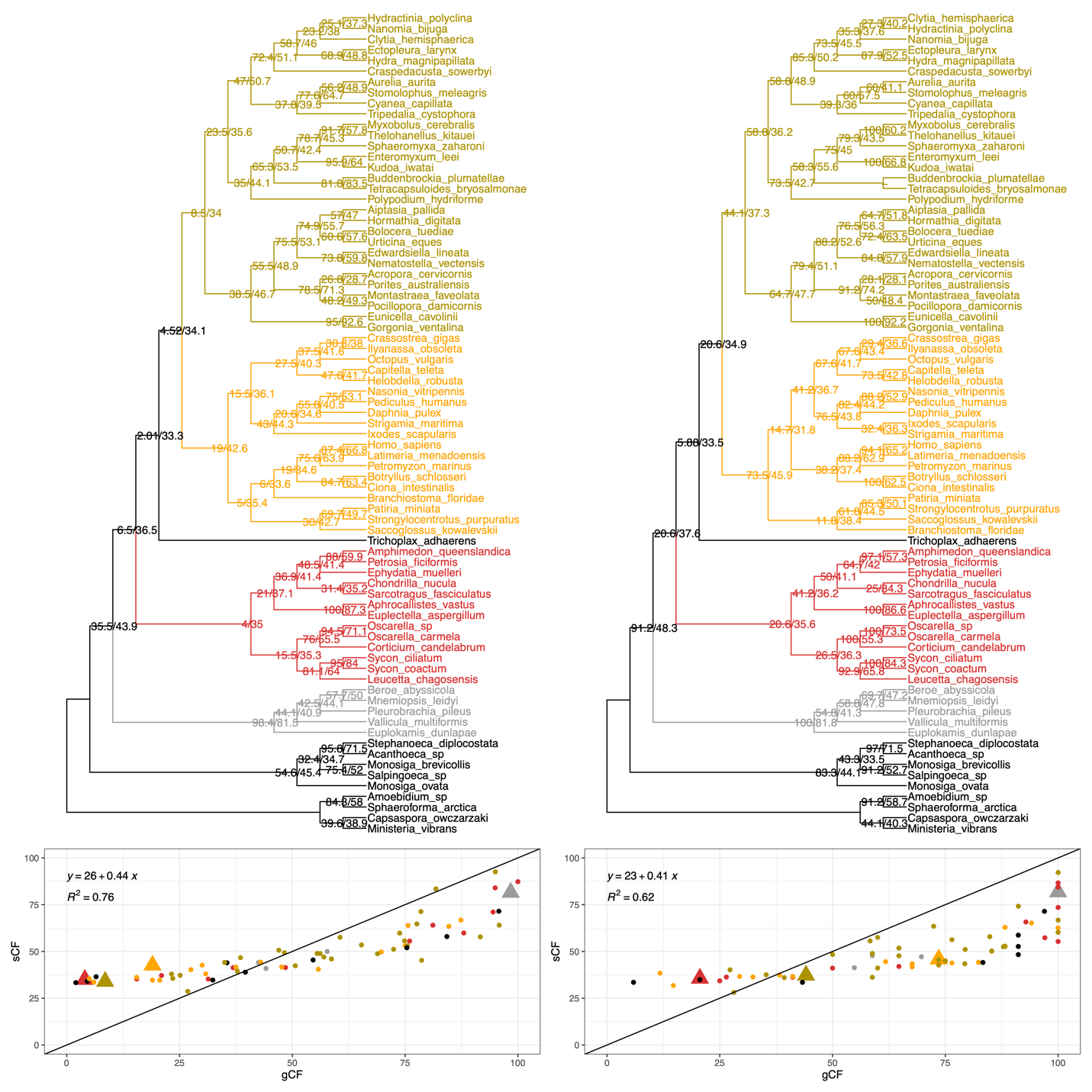

#### **Figure S4**. **Gene and site concordance factor analysis of Chang2015 datasets.** Concordance factors for original and filtered animal datasets calculated using IQTREE. Polytomies in filtered animal phylogenies were manually resolved to facilitate concordance analysis. **Top:** where present, gene concordance (gCF) and site concordance (sCF) factors are plotted onto phylogenies as node values. **Bottom**: scatterplot representing gCF vs. sCF values for each node in a phylogeny, coloured by taxonomy. Triangular points correspond to nodes representing major branches in animal, e.g. the node leading to Bilateria.

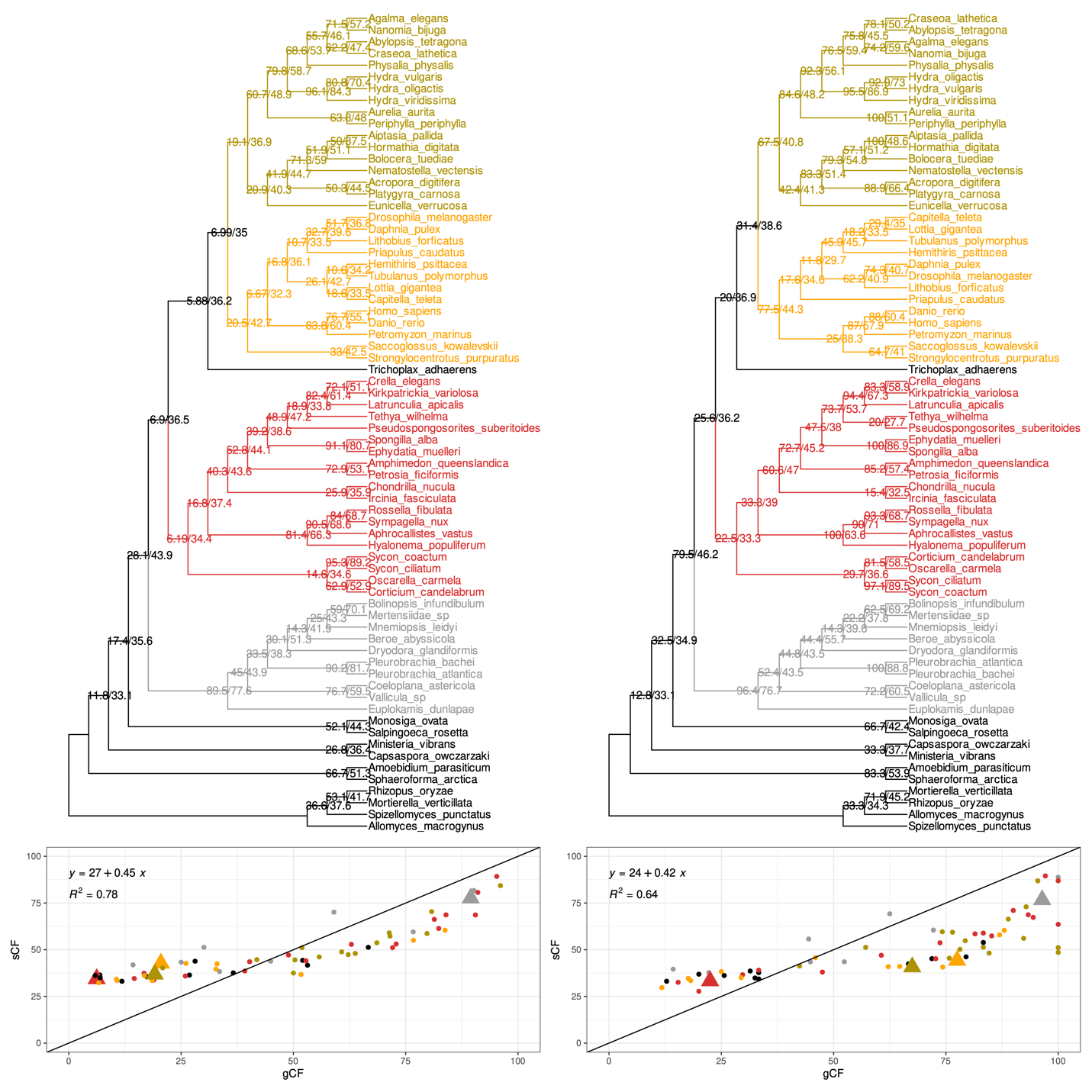

#### **Figure S5**. **Gene and site concordance factor analysis of Whelan2015_D10 datasets.** Concordance factors for original and filtered animal datasets calculated using IQTREE. Polytomies in filtered animal phylogenies were manually resolved to facilitate concordance analysis. **Top:** where present, gene concordance (gCF) and site concordance (sCF) factors are plotted onto phylogenies as node values. **Bottom**: scatterplot representing gCF vs. sCF values for each node in a phylogeny, coloured by taxonomy. Triangular points correspond to nodes representing major branches in animal, e.g. the node leading to Bilateria.

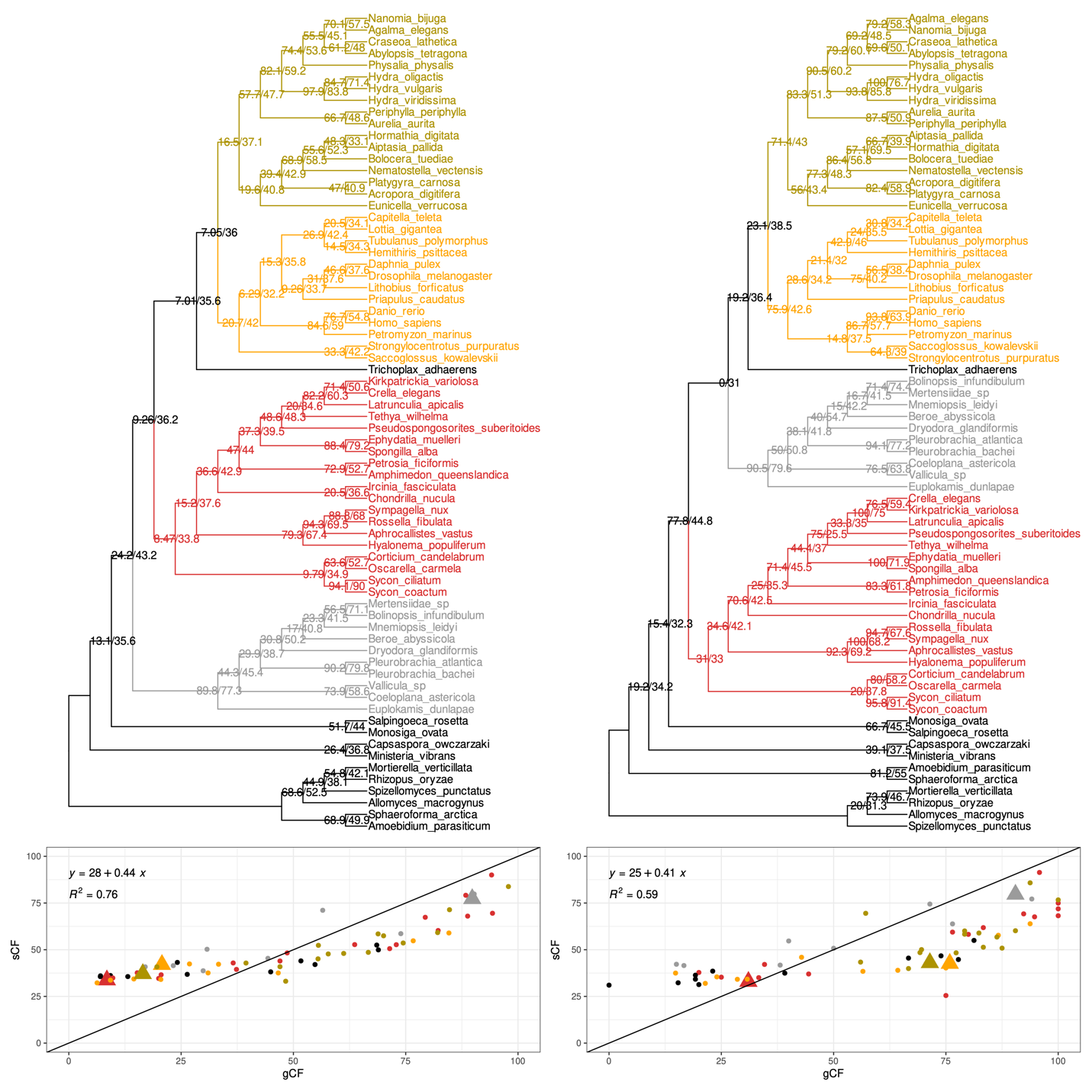

#### **Figure S6**. **Gene and site concordance factor analysis of Whelan2015_D20 datasets.** Concordance factors for original and filtered animal datasets calculated using IQTREE. Polytomies in filtered animal phylogenies were manually resolved to facilitate concordance analysis. **Top:** where present, gene concordance (gCF) and site concordance (sCF) factors are plotted onto phylogenies as node values. **Bottom**: scatterplot representing gCF vs. sCF values for each node in a phylogeny, coloured by taxonomy. Triangular points correspond to nodes representing major branches in animal, e.g. the node leading to Bilateria.

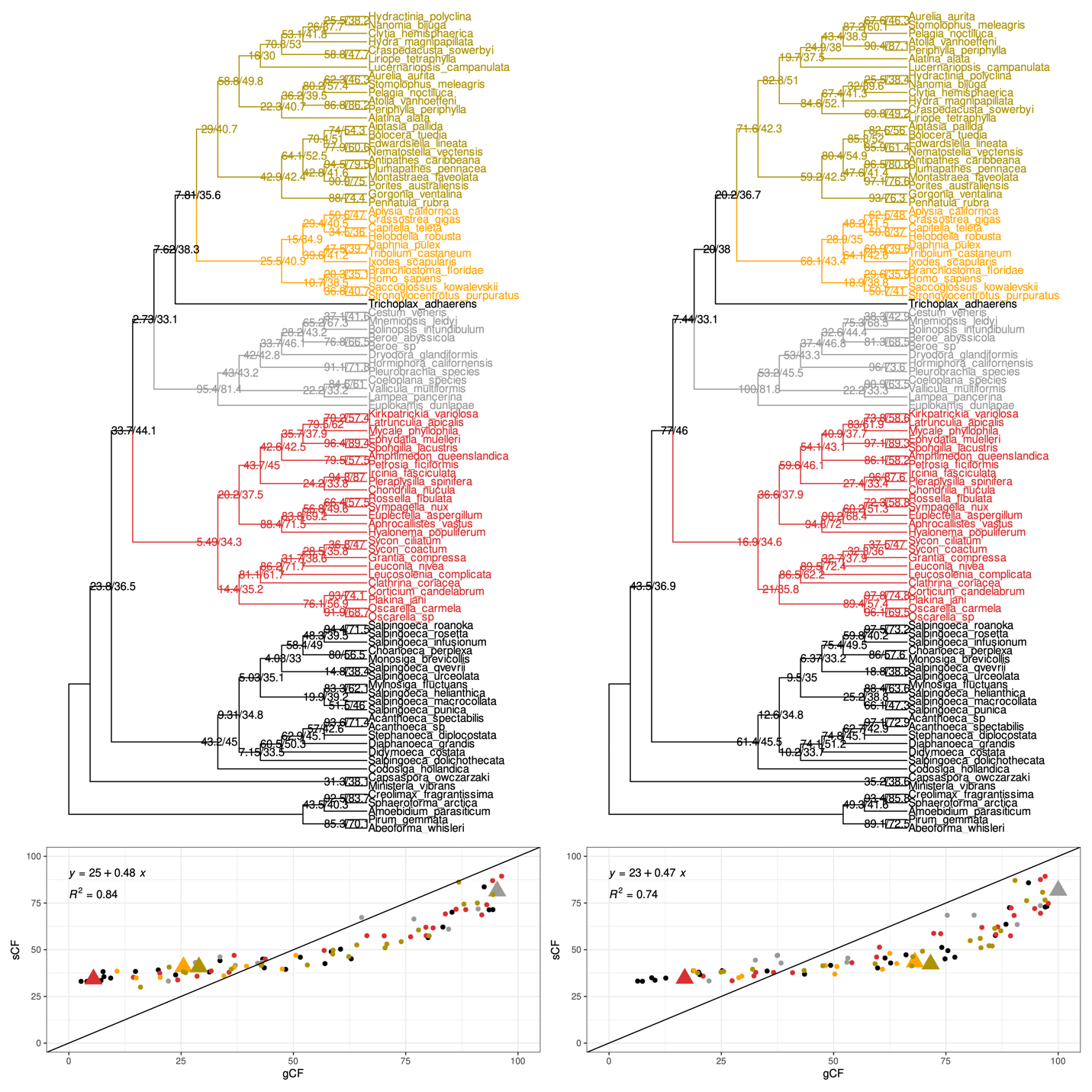

#### **Figure S7**. **Gene and site concordance factor analysis of Simion2017 datasets.** Concordance factors for original and filtered animal datasets calculated using IQTREE. Polytomies in filtered animal phylogenies were manually resolved to facilitate concordance analysis. **Top:** where present, gene concordance (gCF) and site concordance (sCF) factors are plotted onto phylogenies as node values. **Bottom**: scatterplot representing gCF vs. sCF values for each node in a phylogeny, coloured by taxonomy. Triangular points correspond to nodes representing major branches in animal, e.g. the node leading to Bilateria.

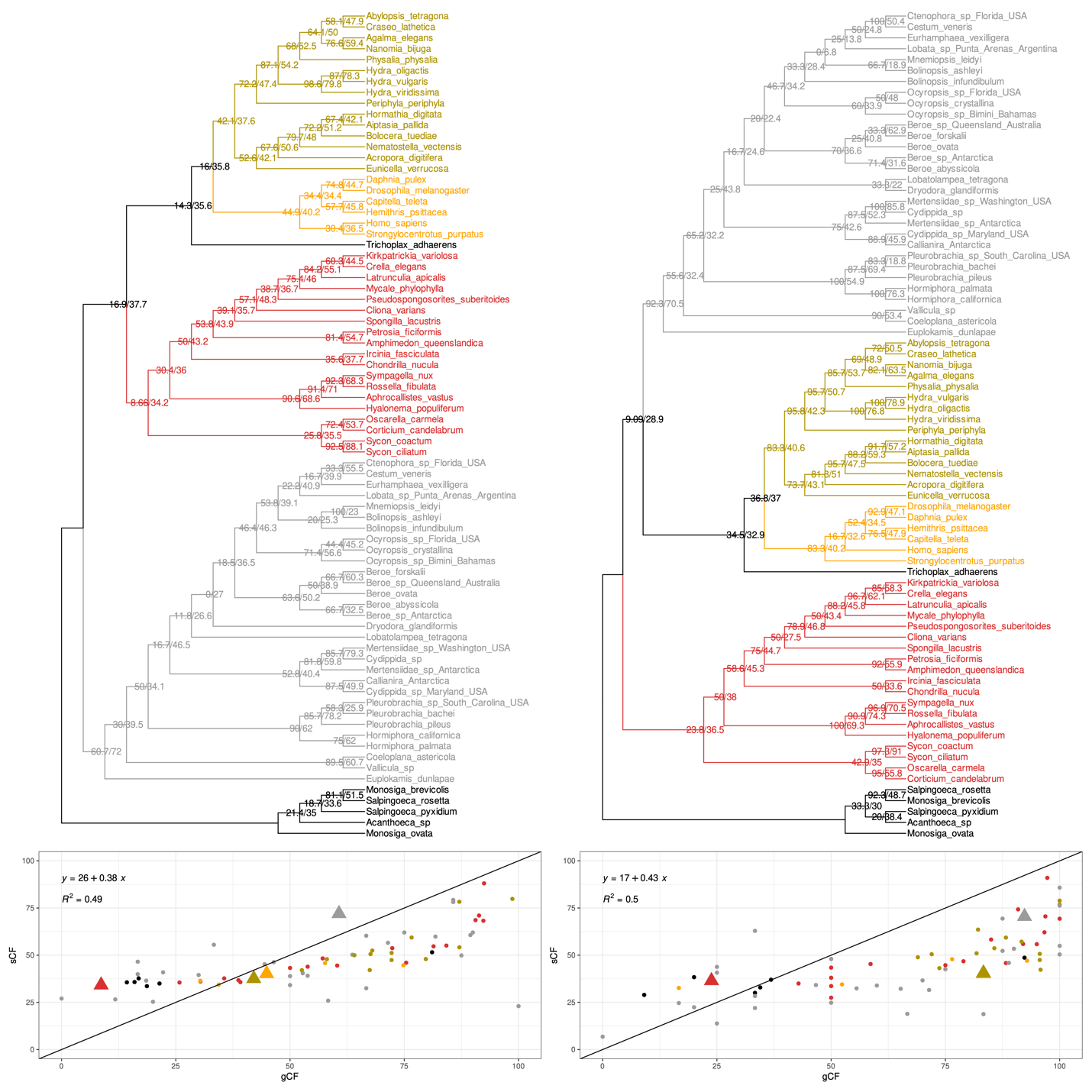

#### **Figure S8**. **Gene and site concordance factor analysis of Whelan2017_MCRS datasets.** Concordance factors for original and filtered animal datasets calculated using IQTREE. Polytomies in filtered animal phylogenies were manually resolved to facilitate concordance analysis. **Top:** where present, gene concordance (gCF) and site concordance (sCF) factors are plotted onto phylogenies as node values. **Bottom**: scatterplot representing gCF vs. sCF values for each node in a phylogeny, coloured by taxonomy. Triangular points correspond to nodes representing major branches in animal, e.g. the node leading to Bilateria.

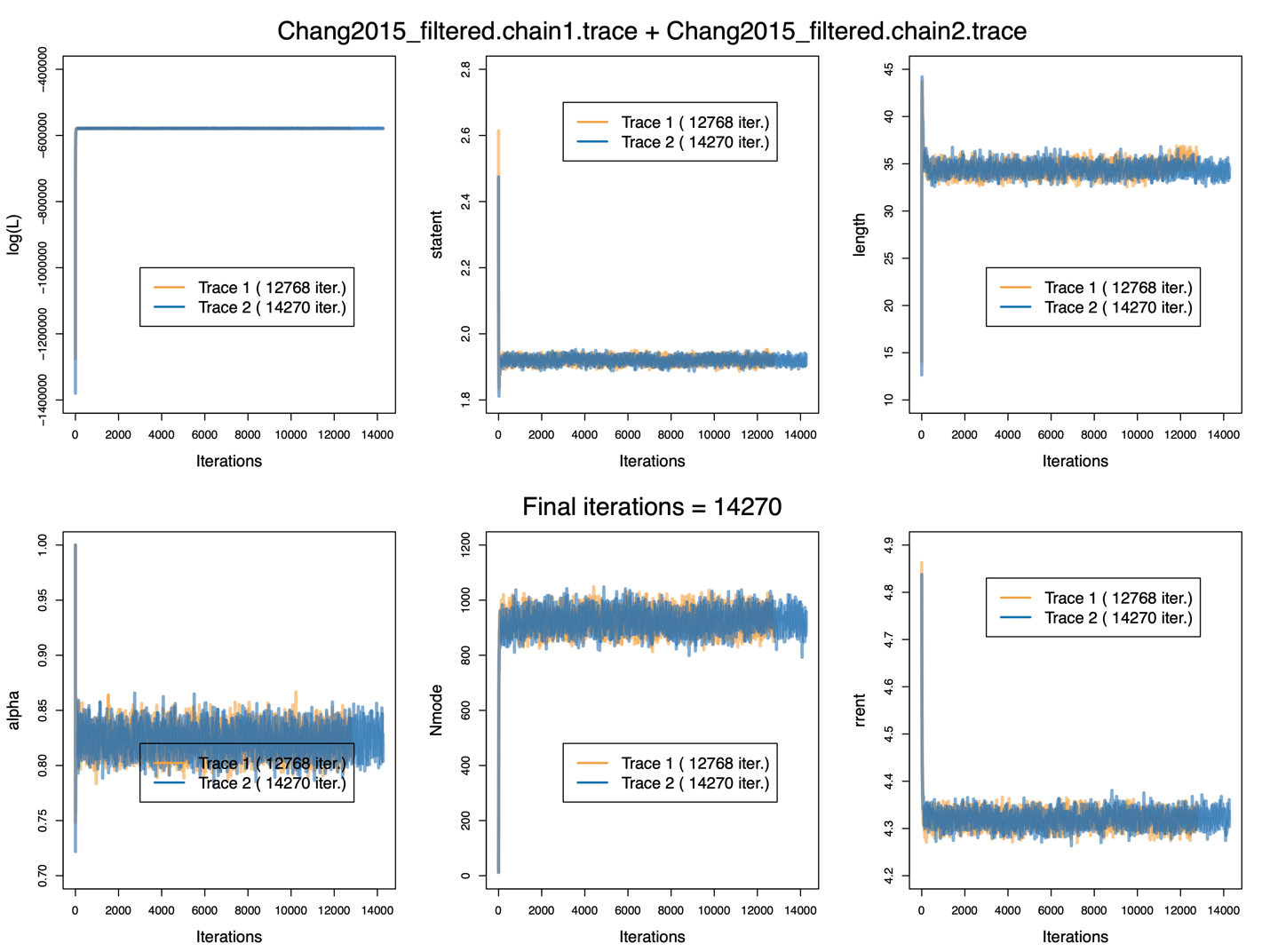

**Figure S9.** **Trace plots for Chang2015_filtered PhyloBayes-MPI run under CAT-GTR+G4 model**. Plots represent major components of mixture model. Log(L): log-likelihood, statent: mean site entropy, length: tree length, alpha: α parameter of gamma distribution of rates across sites, Nmode: occupied components of mixture, rrent: entropy of exchangeabilities.

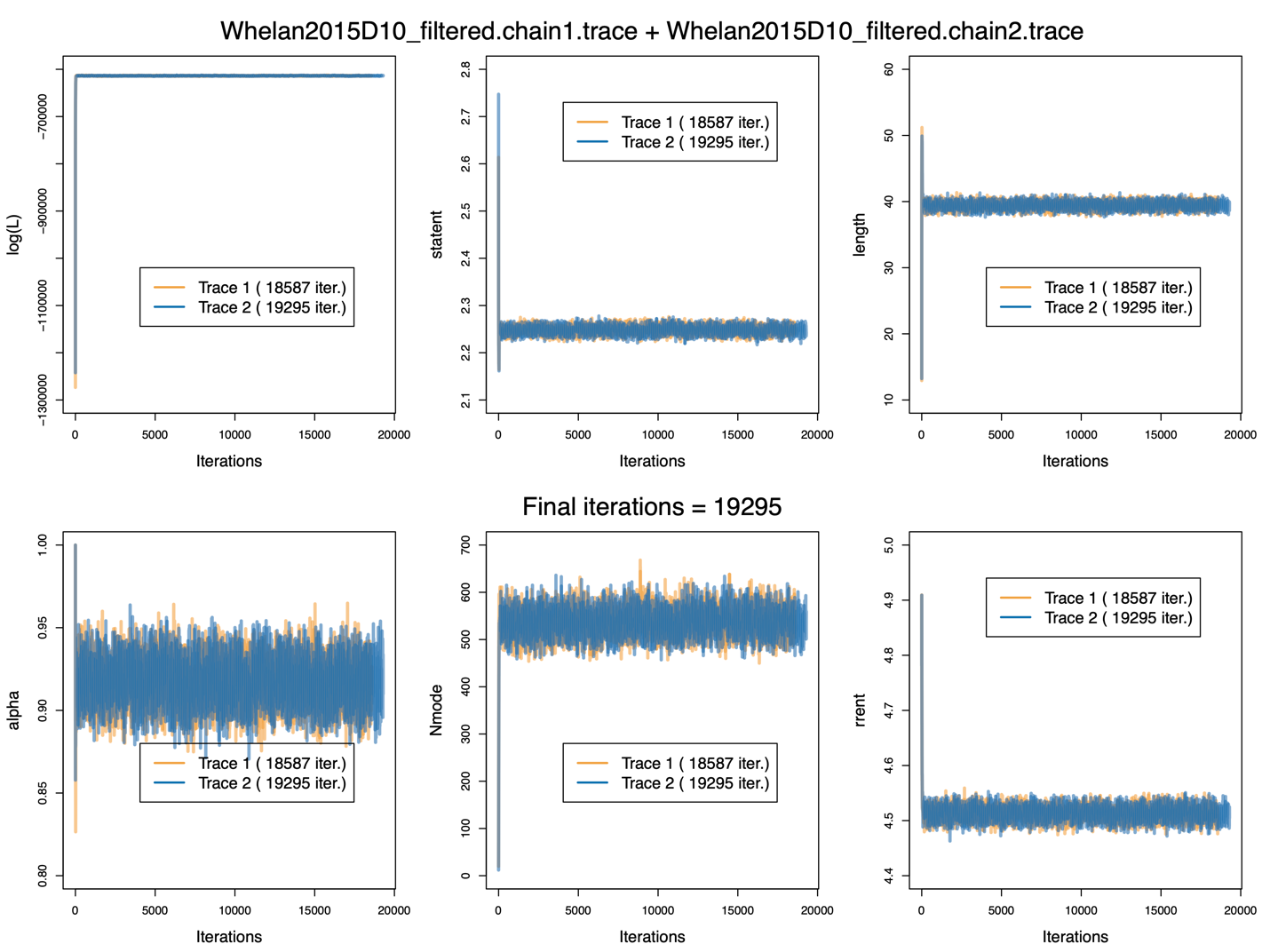

**Figure S10.** **Trace plots for Whelan2015D10_filtered PhyloBayes-MPI run under CAT-GTR+G4 model.** Plots represent major components of mixture model. Log(L): log-likelihood, statent: mean site entropy, length: tree length, alpha: α parameter of gamma distribution of rates across sites, Nmode: occupied components of mixture, rrent: entropy of exchangeabilities.

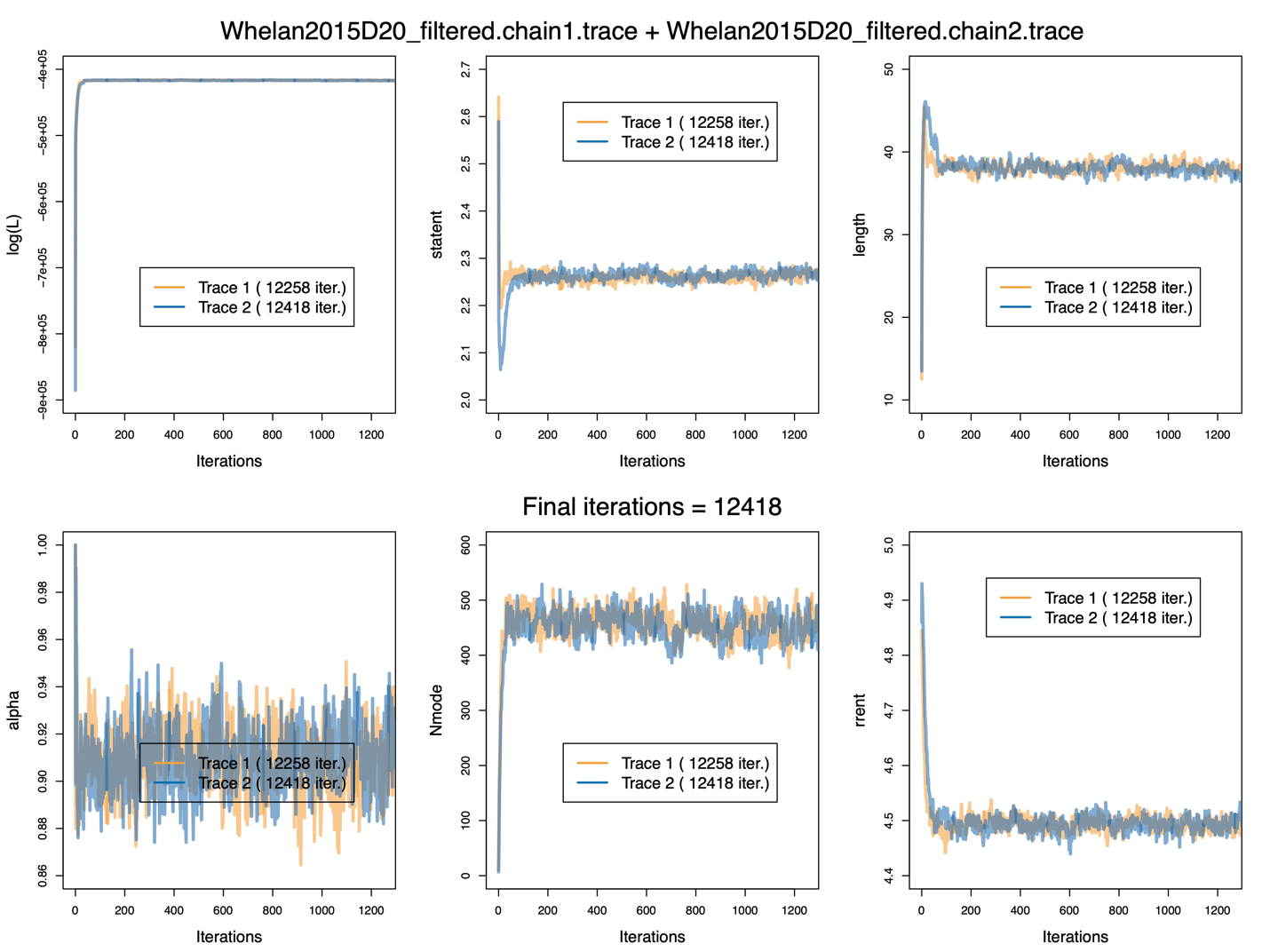

**Figure S11.** **Trace plots for Whelan2015D20_filtered PhyloBayes-MPI run under CAT-GTR+G4 model.** Plots represent major components of mixture model. Log(L): log-likelihood, statent: mean site entropy, length: tree length, alpha: α parameter of gamma distribution of rates across sites, Nmode: occupied components of mixture, rrent: entropy of exchangeabilities.

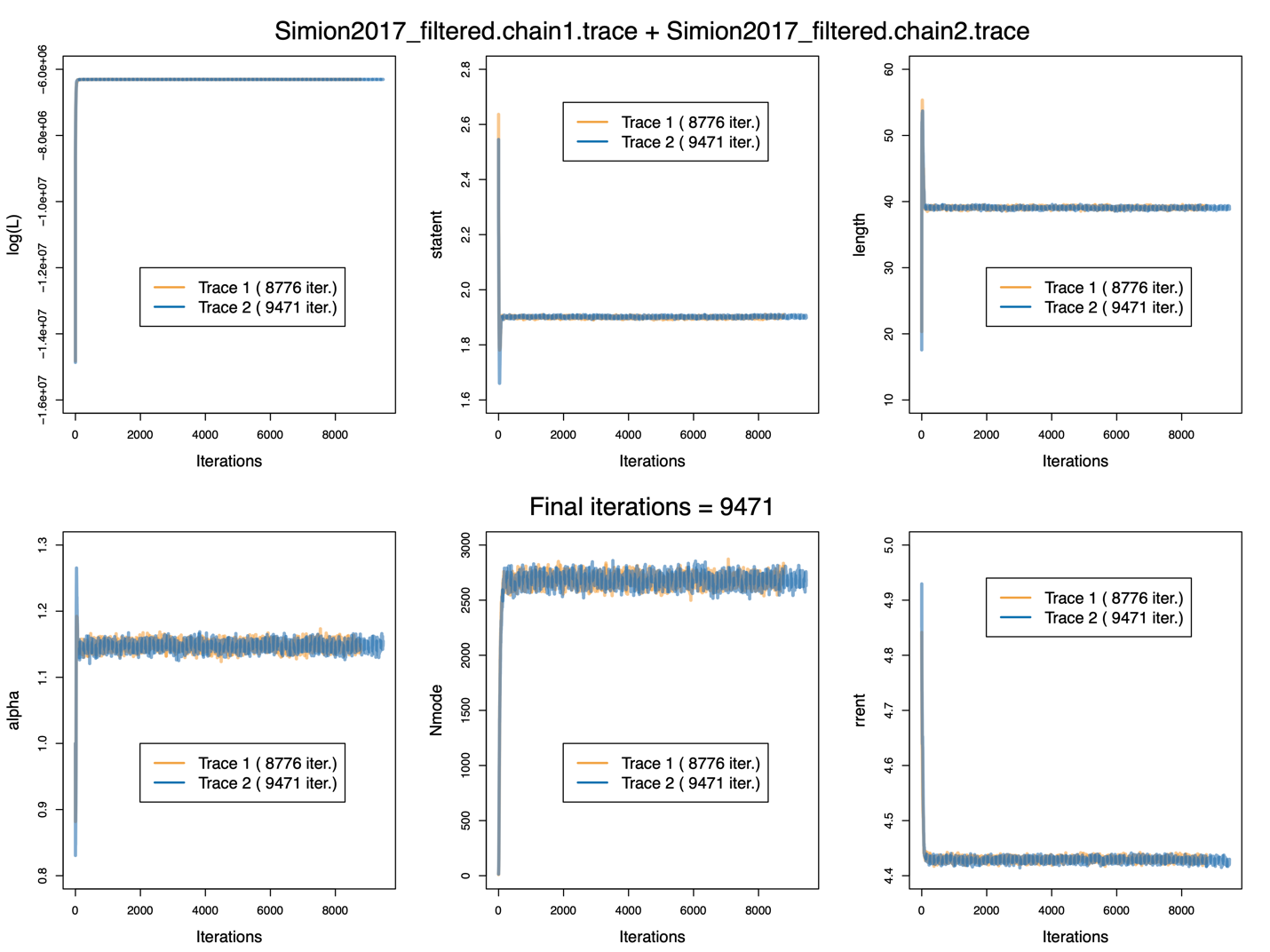

**Figure S12.** **Trace plots for Simion2017_filtered PhyloBayes-MPI run under CAT-GTR+G4 model.** Plots represent major components of mixture model. Log(L): log-likelihood, statent: mean site entropy, length: tree length, alpha: α parameter of gamma distribution of rates across sites, Nmode: occupied components of mixture, rrent: entropy of exchangeabilities.

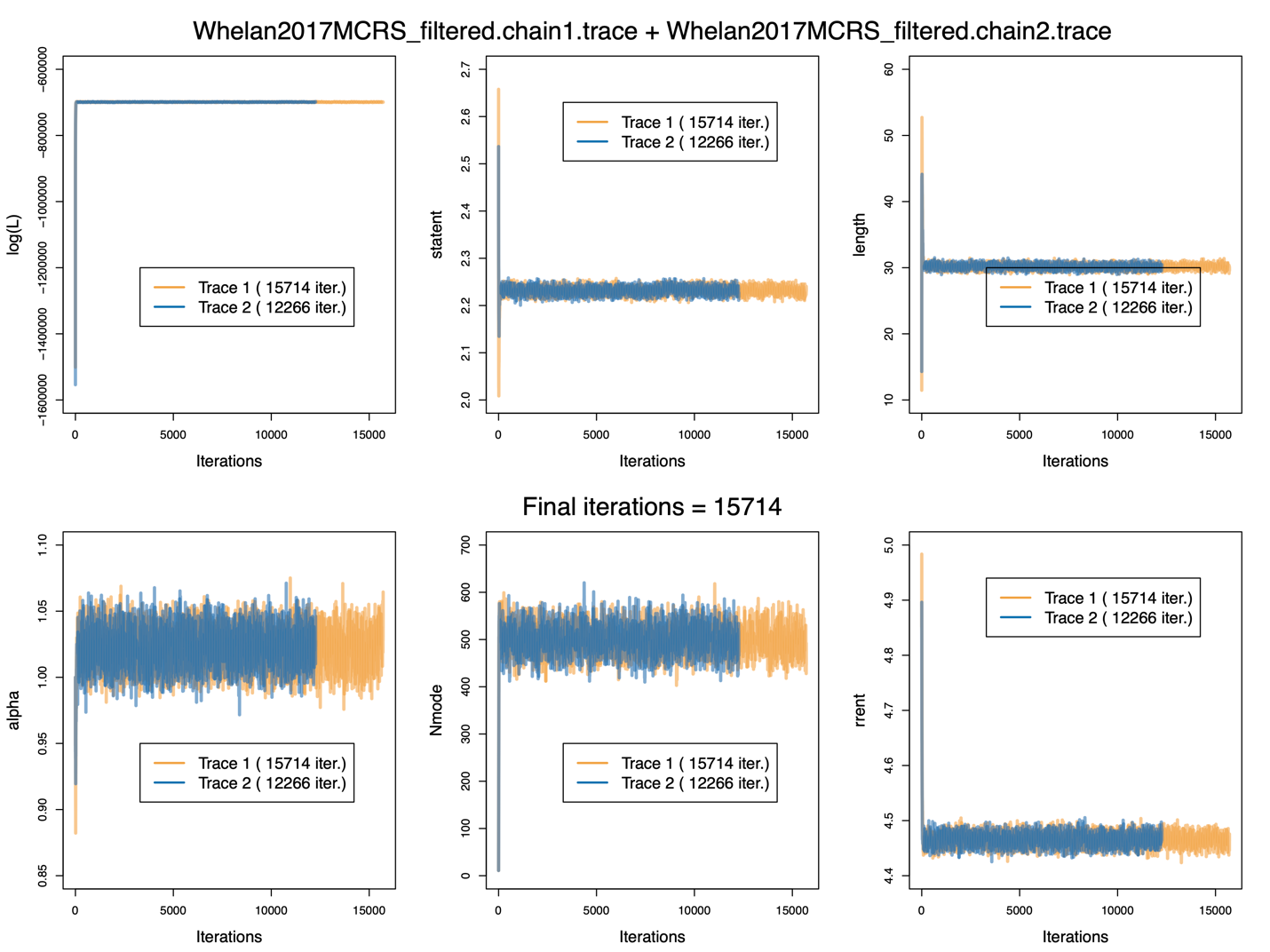

**Figure S13.** **Trace plots for Whelan2017MCRS_filtered PhyloBayes-MPI run under CAT-GTR+G4 model.** Plots represent major components of mixture model. Log(L): log-likelihood, statent: mean site entropy, length: tree length, alpha: α parameter of gamma distribution of rates across sites, Nmode: occupied components of mixture, rrent: entropy of exchangeabilities.

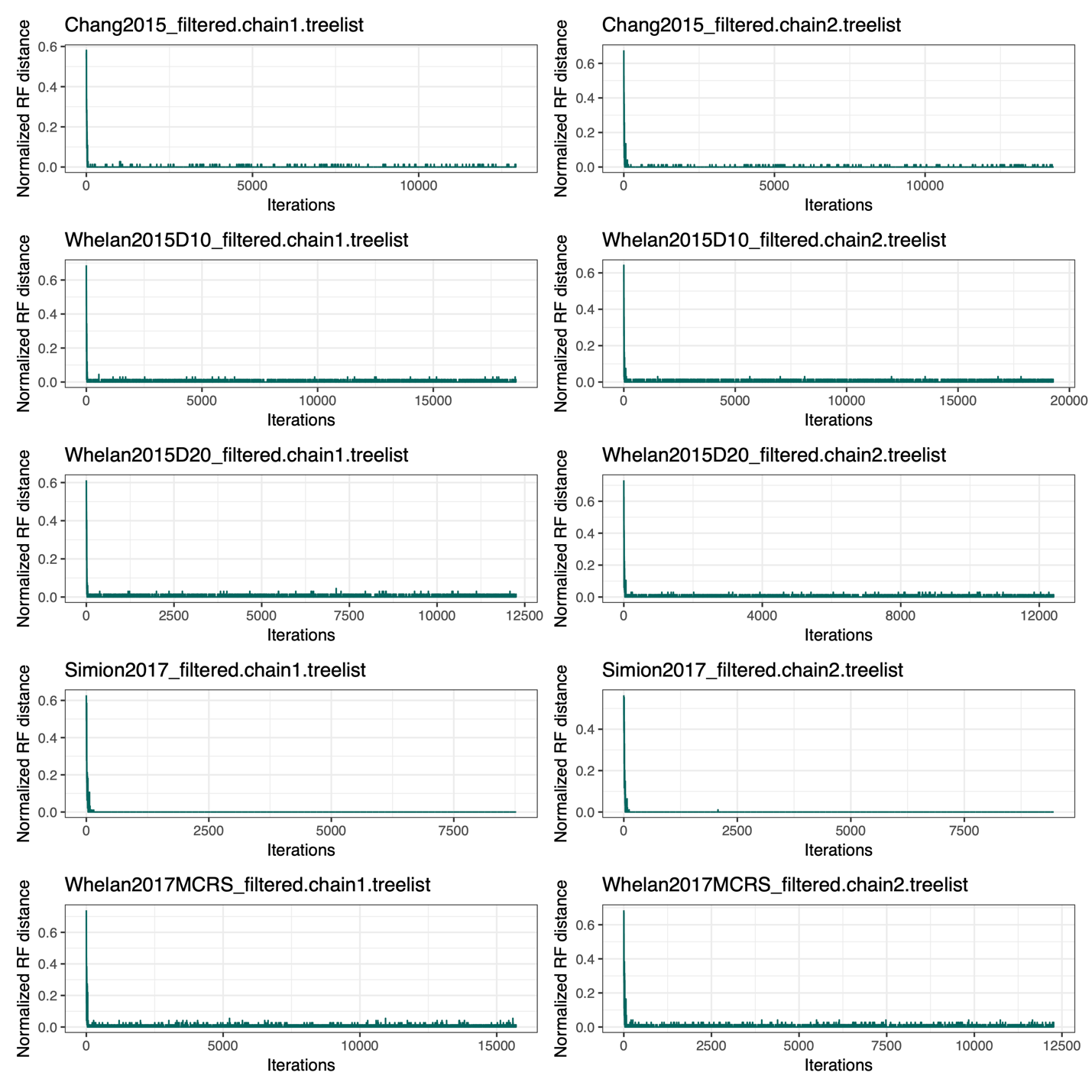

**Figure S14.** **Normalized pairwise Robinson-Foulds (RF) distance plots for each chain of each PhyloBayes-MPI run under the CAT-GTR+G4 model.** RF distances calculated between trees at each iteration of a given chain.

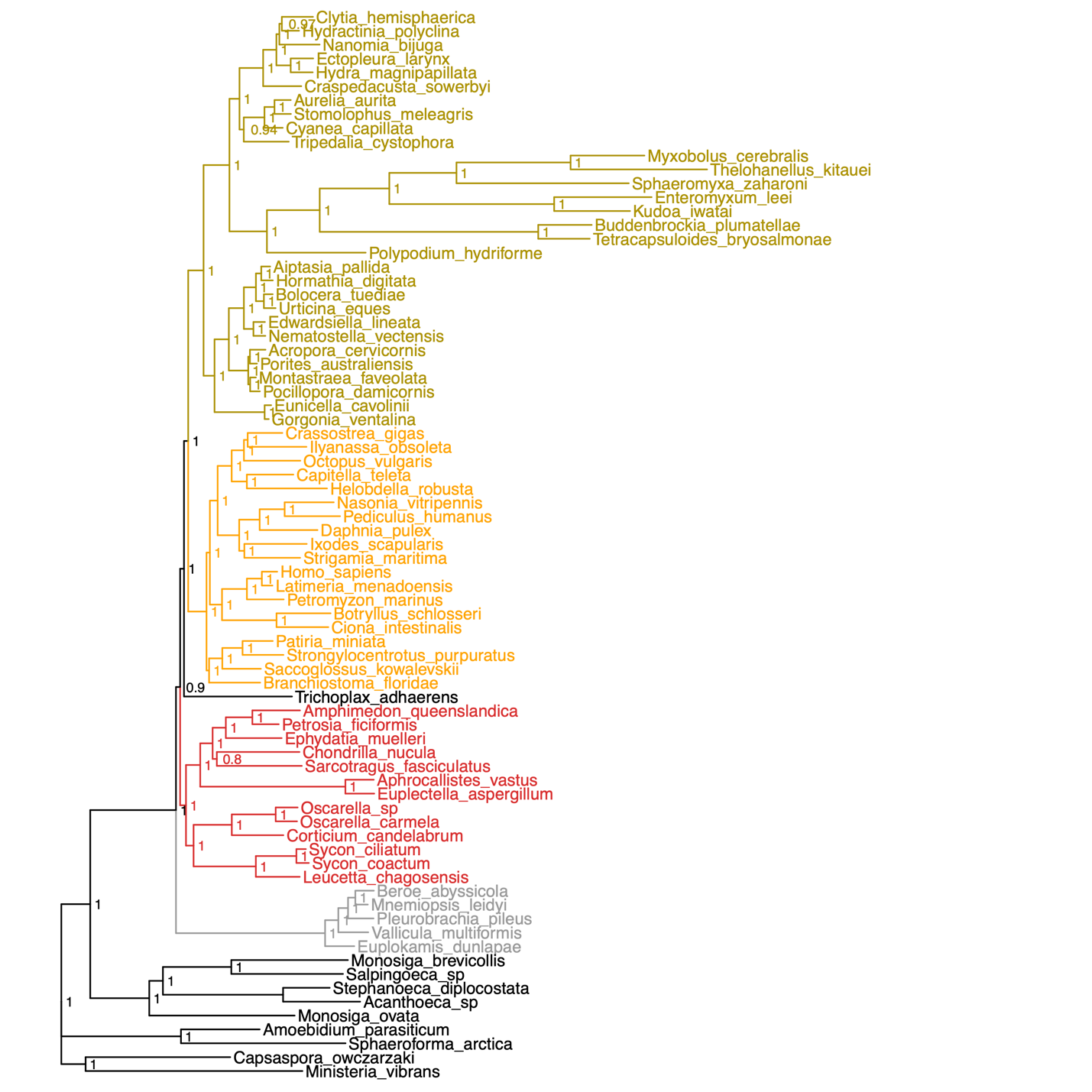

**Figure S15.** **Chang2015_filtered CAT-GTR+G4 phylogeny.** Animal subgroups coloured by taxonomy. Posterior probabilties given at each node.

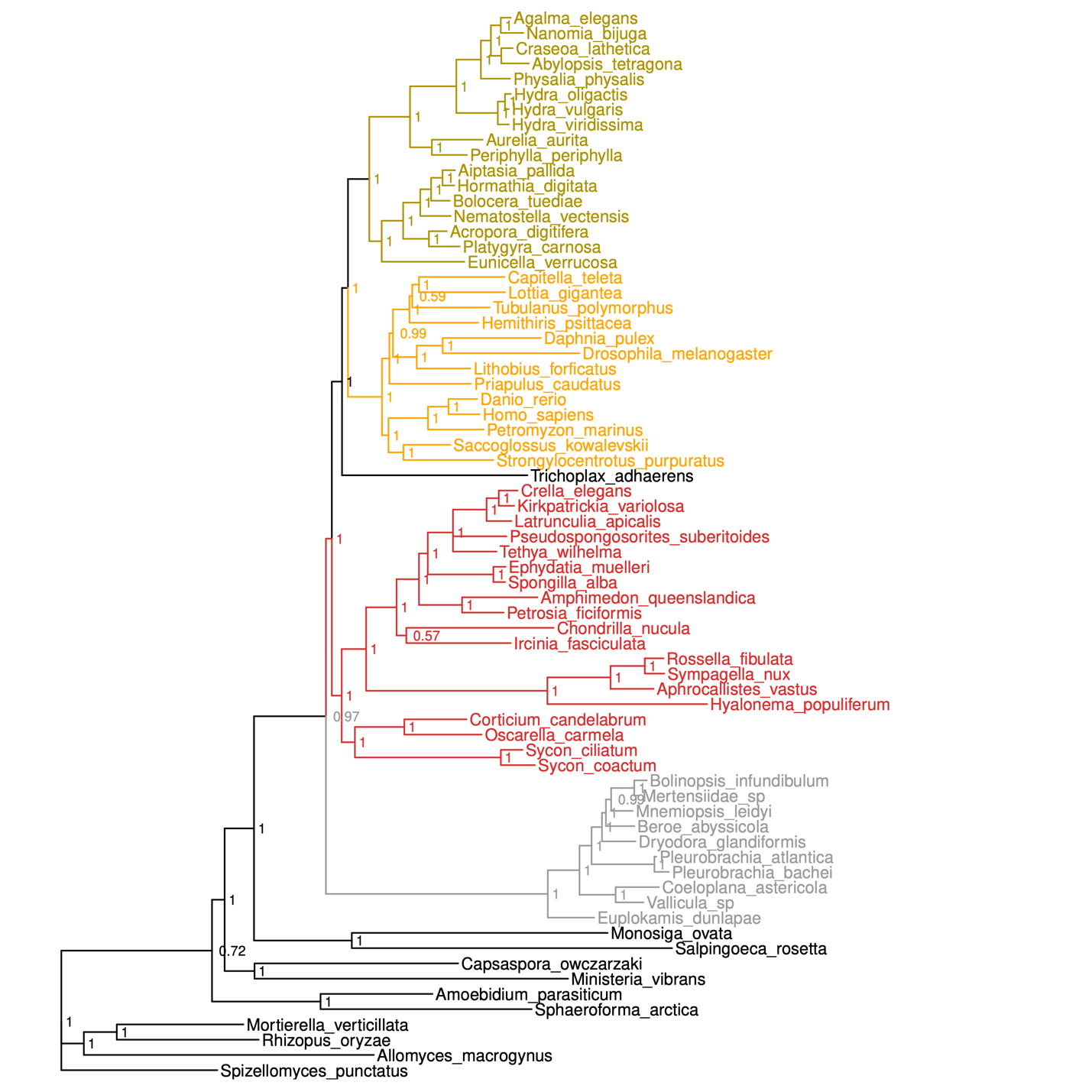

**Figure S16.** **Whelan2015D10_filtered CAT-GTR+G4 phylogeny.** Animal subgroups coloured by taxonomy. Posterior robabilities given at each node.

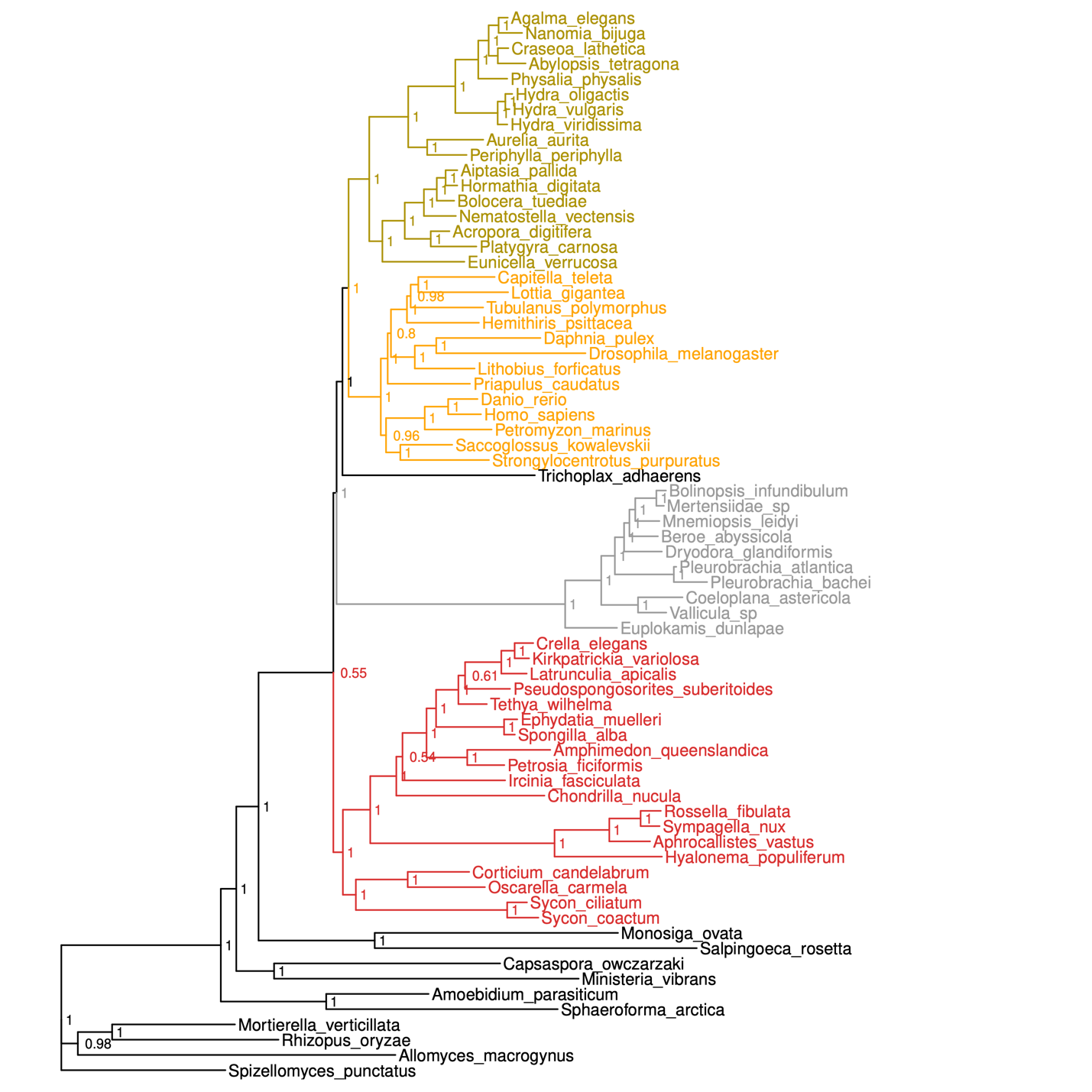

**Figure S17.** **Whelan2015D20_filtered CAT-GTR+G4 phylogeny.** Animal subgroups coloured by taxonomy. Posterior probabilties given at each node.

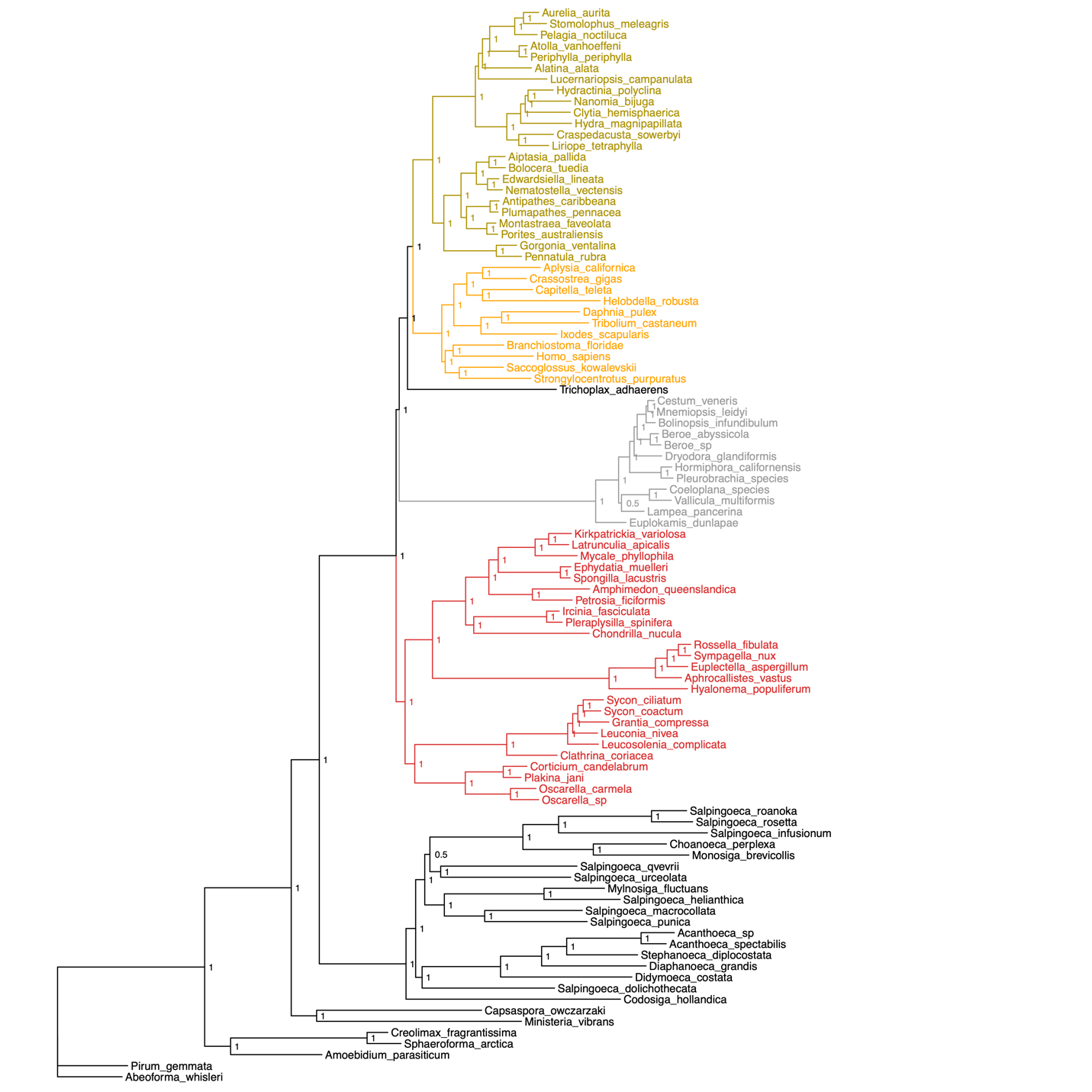

**Figure S18.** **Simion2017_filtered CAT-GTR+G4 phylogeny.** Animal subgroups coloured by taxonomy. Posterior probabilties given at each node.

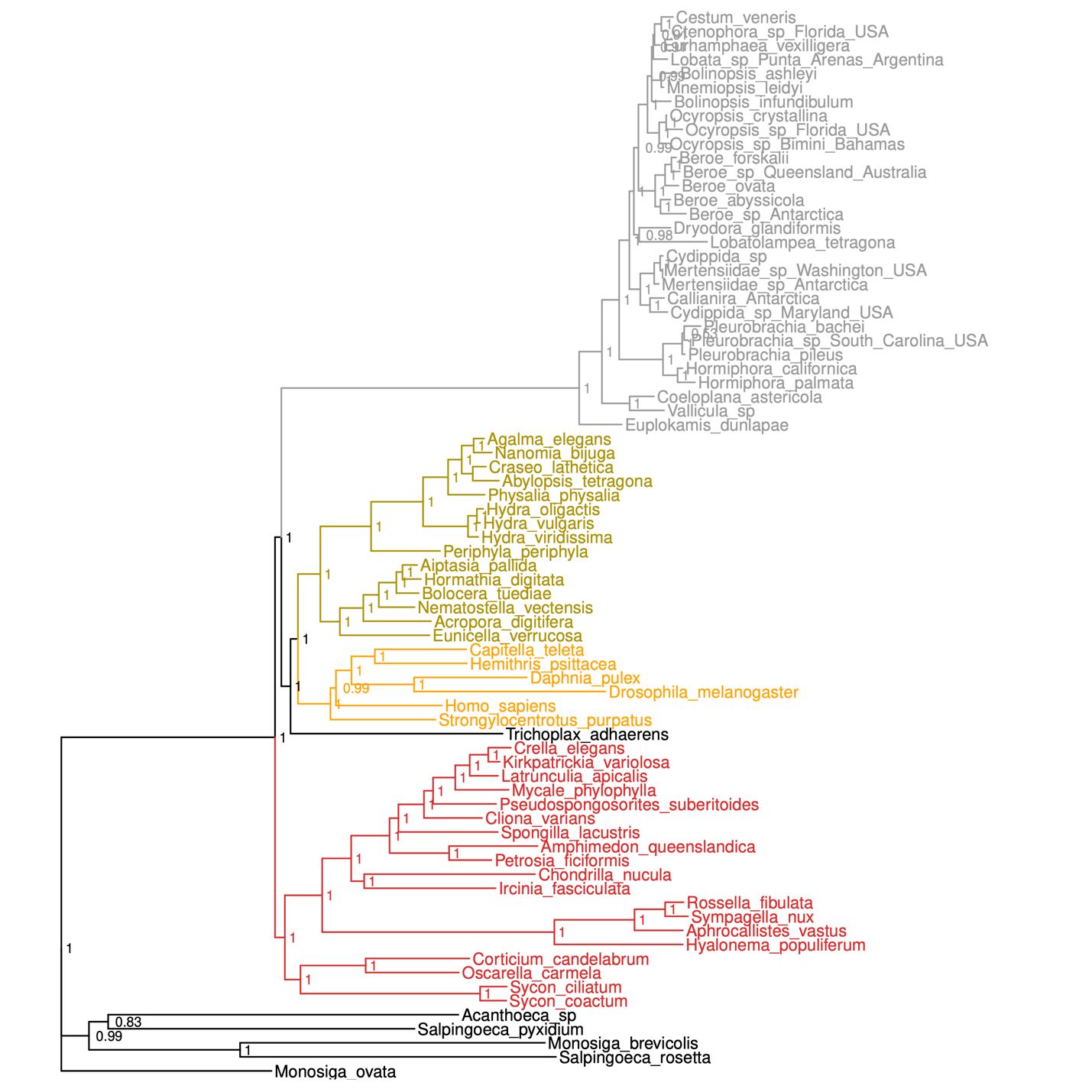

**Figure S19.** **Whelan2017MCRS_filtered CAT-GTR+G4 phylogeny.** Animal subgroups coloured by taxonomy. Posterior probabilties given at each node.

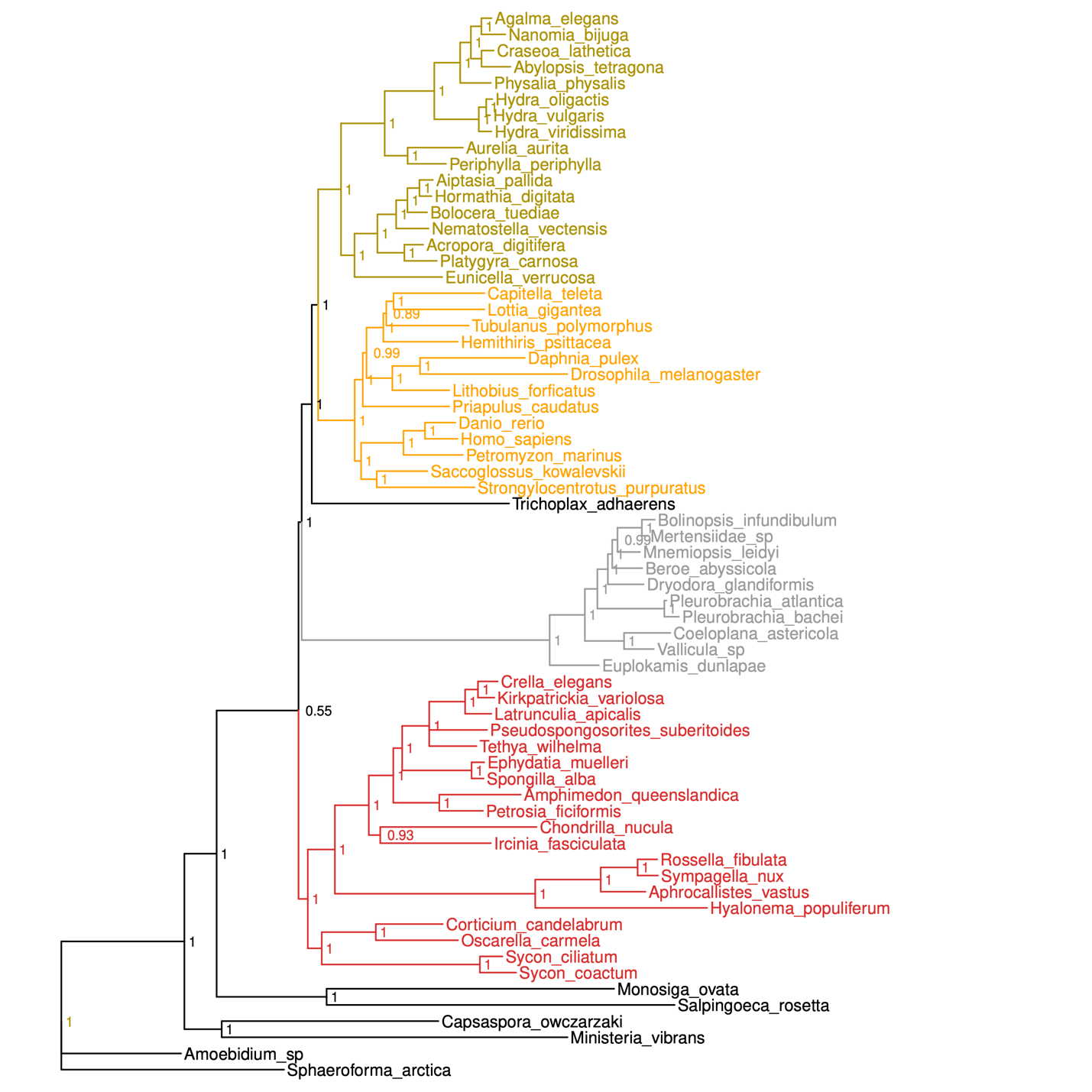

**Figure S20.** **Whelan2015_D10_filteredHolo CAT-GTR+G4 phylogeny.** Animal subgroups coloured by taxonomy. Posterior probabilties given at each node.

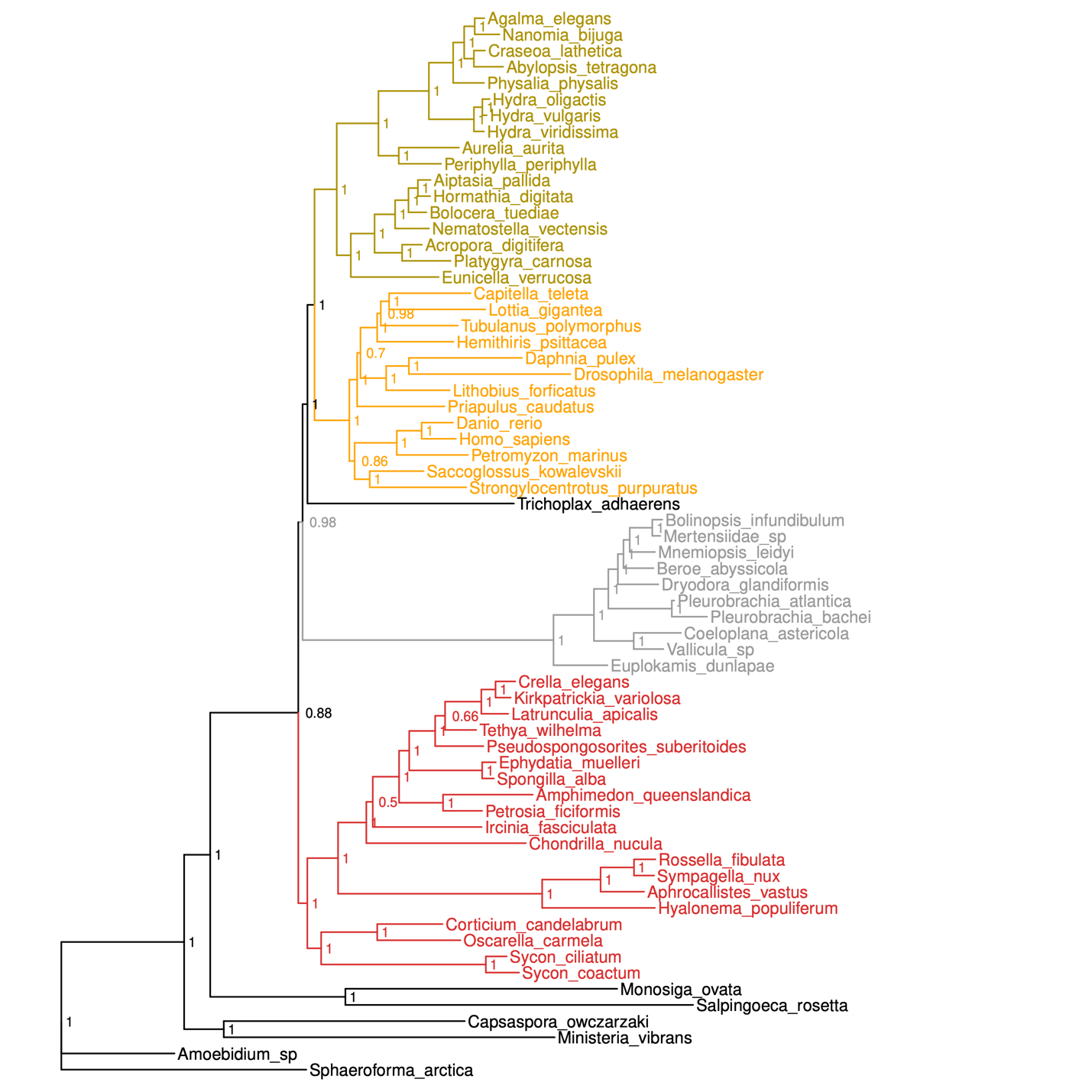

**Figure S21.** **Whelan2015_D20_filteredHolo CAT-GTR+G4 phylogeny.** Animal subgroups coloured by taxonomy. Posterior probabilties given at each node.

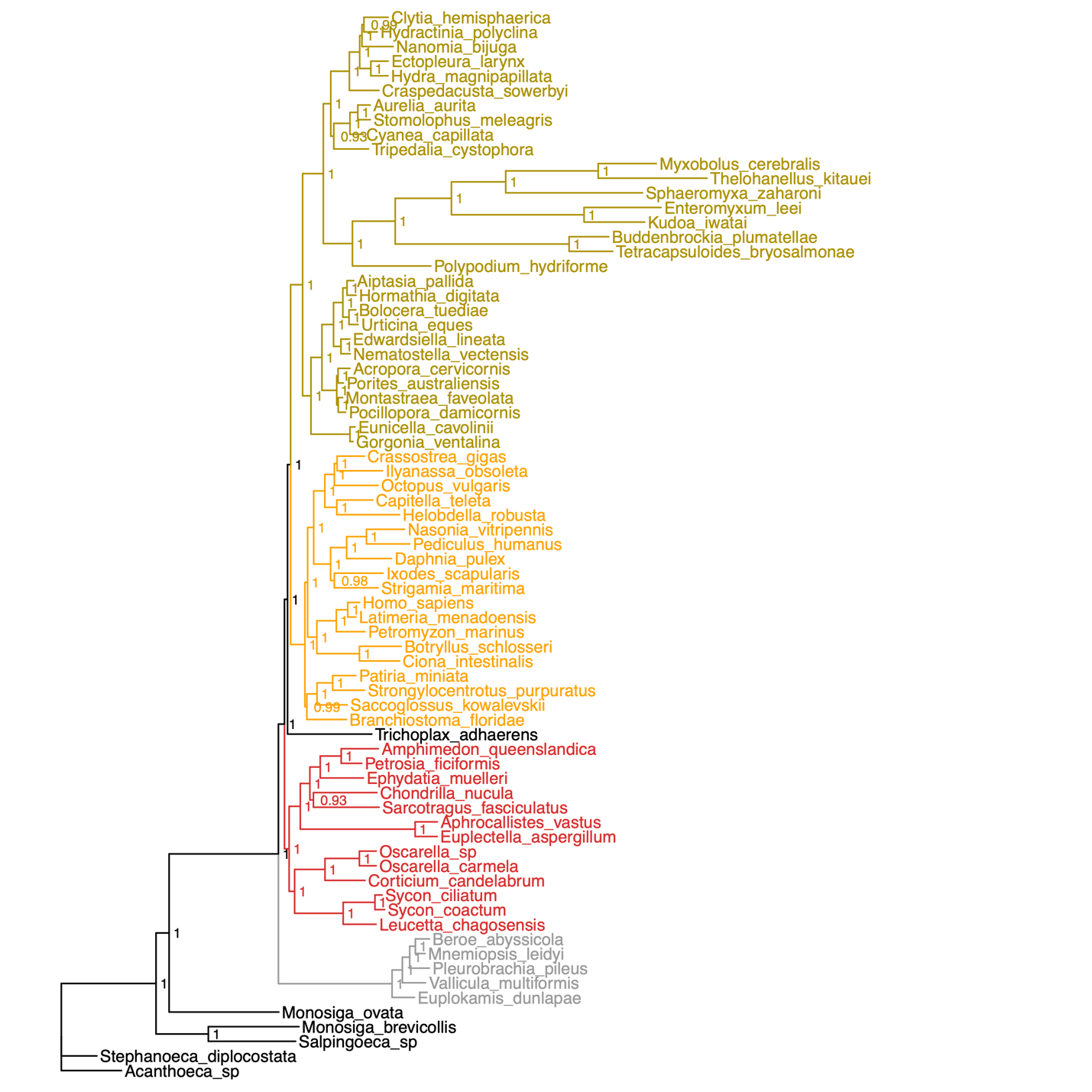

**Figure S22.** **Chang2015_filteredChoano CAT-GTR+G4 phylogeny.** Animal subgroups coloured by taxonomy. Posterior probabilties given at each node.

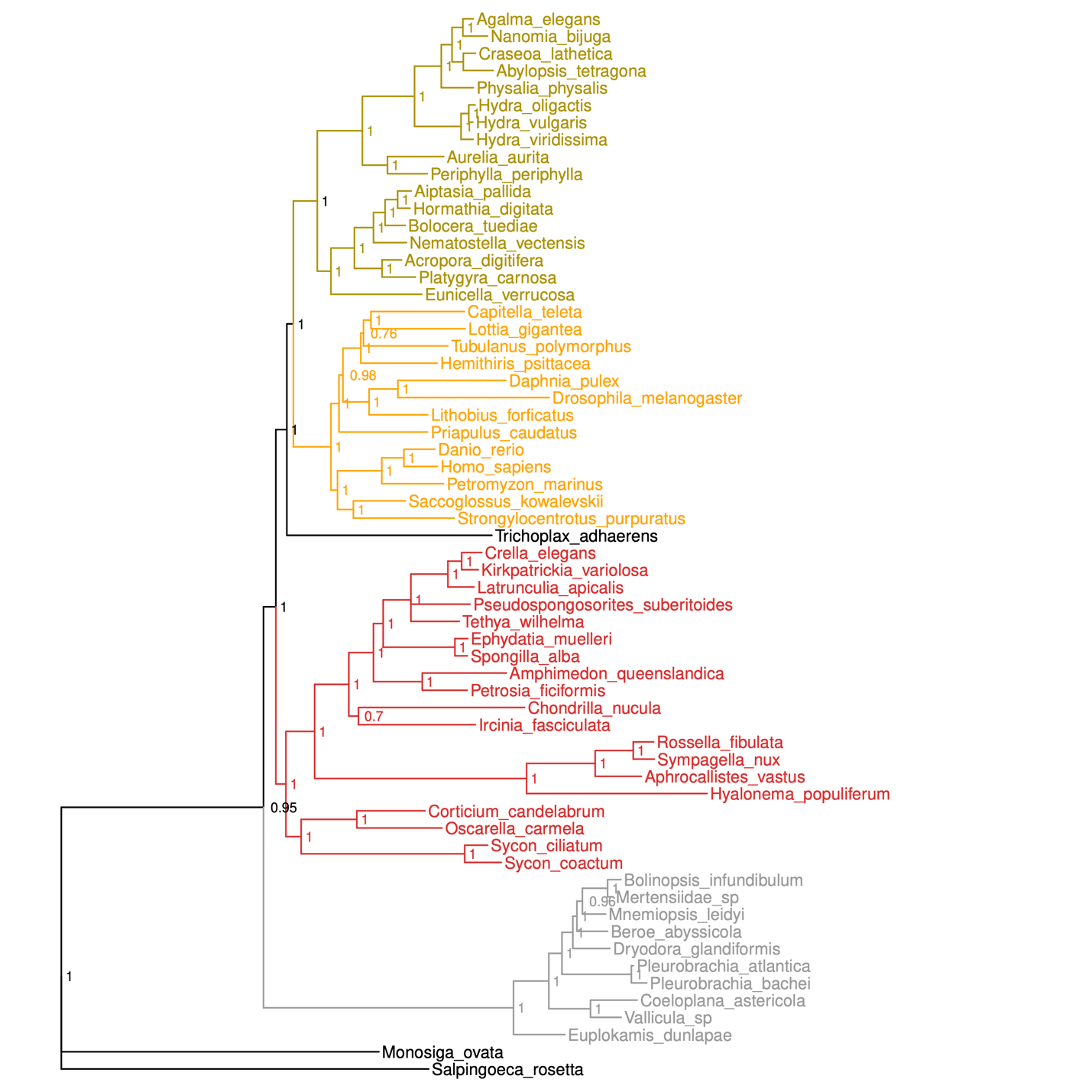

**Figure S23.** **Whelan2015_D10_filteredChoano CAT-GTR+G4 phylogeny.** Animal subgroups coloured by taxonomy. Posterior probabilties given at each node.

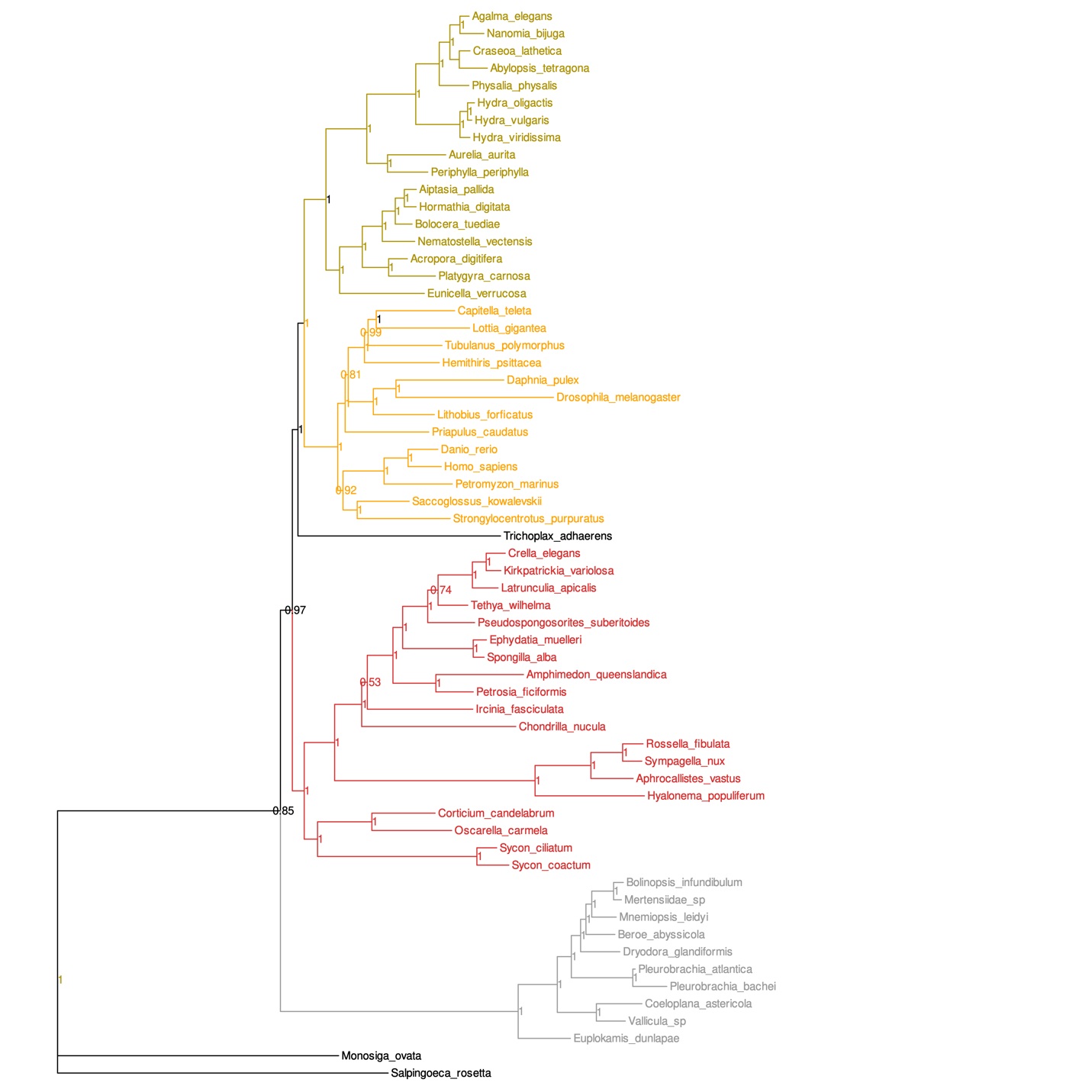

**Figure S24.** **Whelan2015_D20_filteredChoano CAT-GTR+G4 phylogeny.** Animal subgroups coloured by taxonomy. Posterior probabilties given at each node.

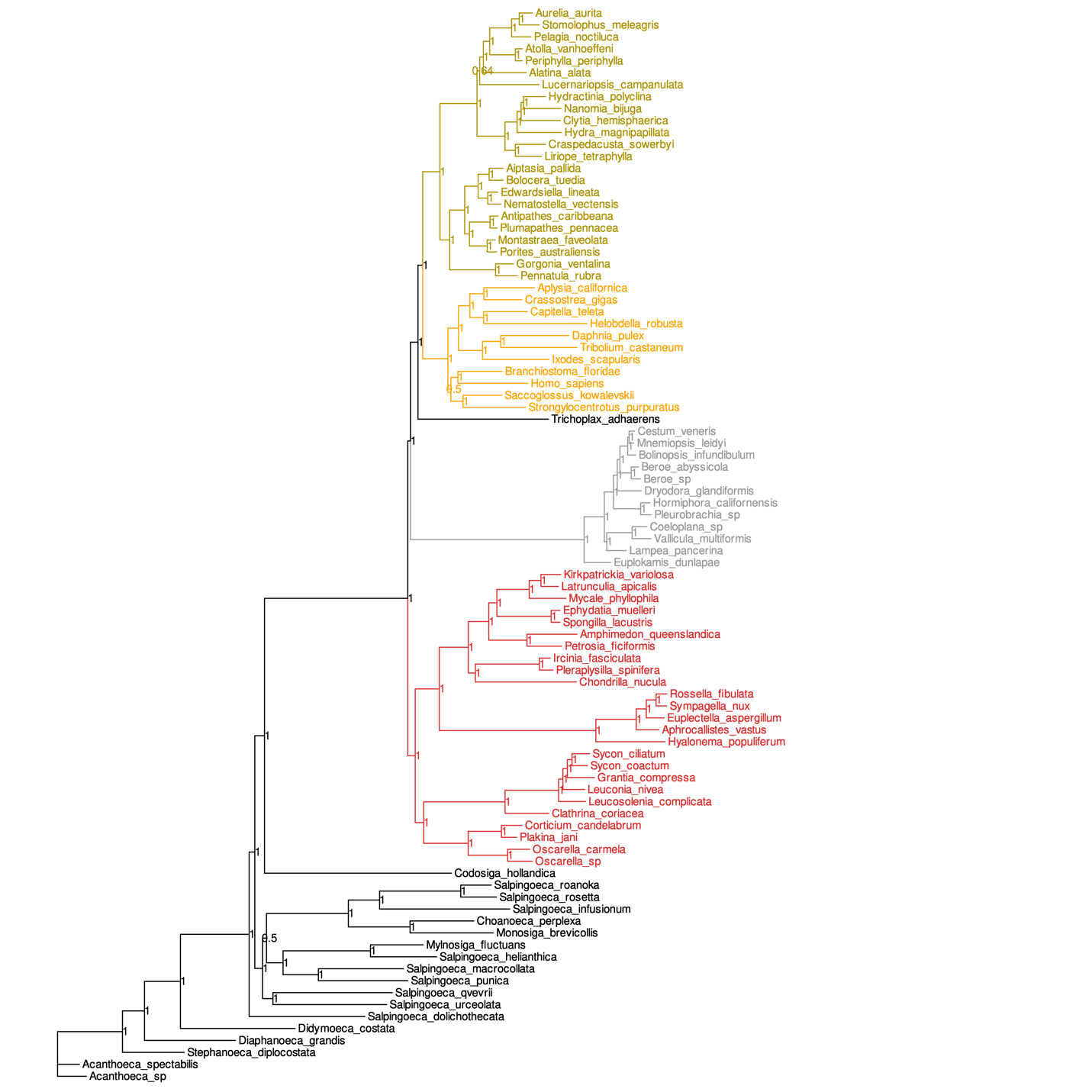

**Figure S25.** **Simion2017_filteredChoano CAT-GTR+G4 phylogeny.** Animal subgroups coloured by taxonomy. Posterior probabilties given at each node.

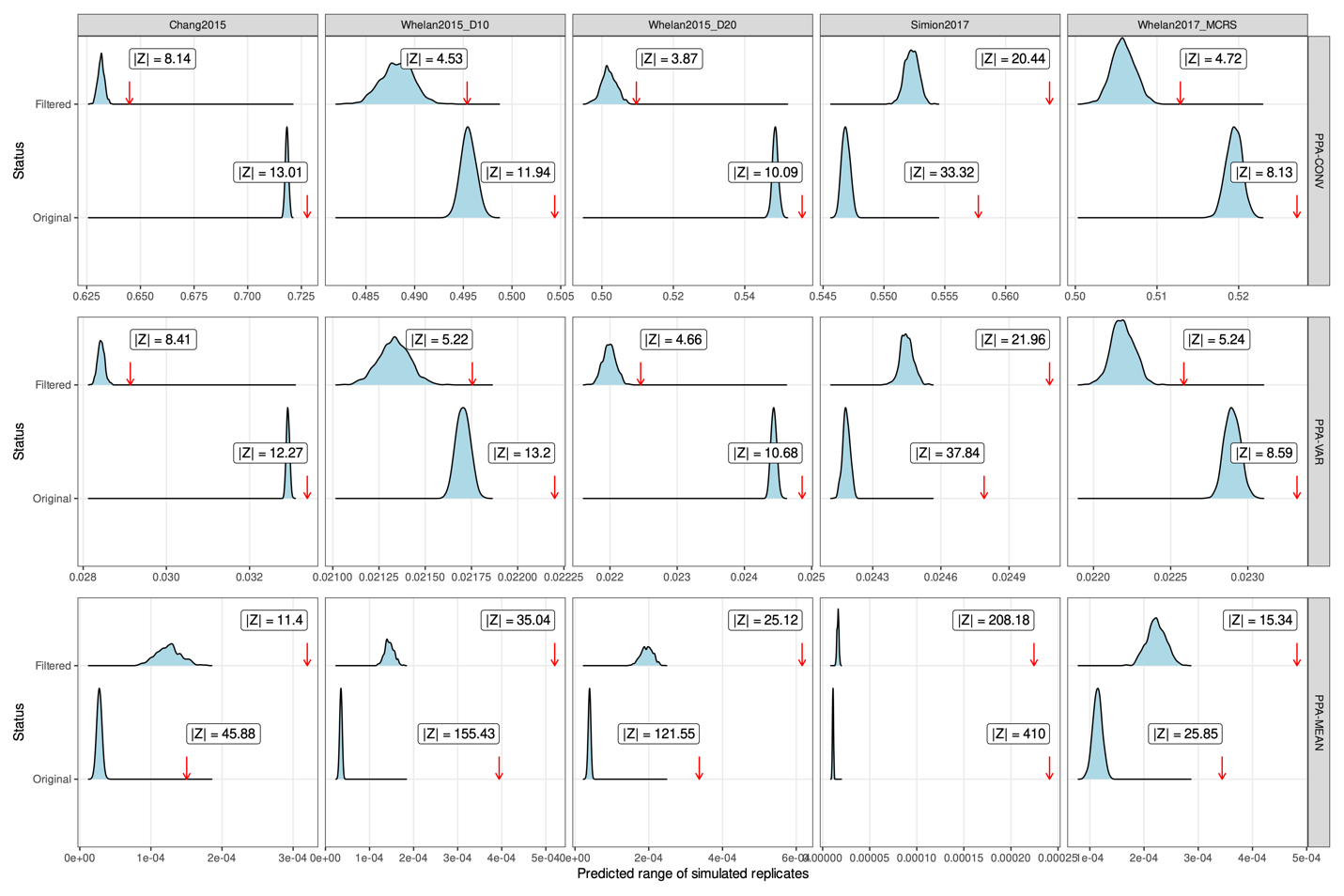

**Figure S26**. **Model fit assessment for CAT-GTR+G reconstructions of filtered animal datasets.** Red arrows indicate the observed mean for each PPA calculated for original and filtered datasets. Blue ridgeline curves represent the range of values for each PPA estimated from the CAT-GTR+G model using 500 simulated replicates, given a predicted mean and standard deviation estimated from PhyloBayes-MPI. Values for the original Chang2015 and Whelan2015_D20 datasets taken from Feuda et al. (2017). PPA statistics shown: probability of convergence towards the same amino acid in distantly related taxa (PPA-VAR), variance in empirical frequencies of each amino acid (PPA-CONV) and squared mean compositional heterogeneity (PPA-MEAN).

### **Supplementary Datasets**

Spreadsheet files are also available from https://github.com/chmccarthy/ATOLRootStudy.

**Dataset S1.** Taxon sampling across five previously-published animal datasets.

**Dataset S2.** Clan_Check results for five previously-published animal datasets.
